# Integrated transcriptomic and epigenomic profiling of human cartilage reveals coordinated changes in fibronectin expression, splicing, and receptor signaling during osteoarthritis progression

**DOI:** 10.64898/2026.10.29.762608

**Authors:** Hannah Swahn, Takuya Sakamoto, Chelsea Kenvisay, Merissa Olmer, Yutaro Uchida, Hiroshi Asahara, Rachel E. Miller, Anne-Marie Malfait, Dana E. Orange, Bella Mehta, Miguel Otero, William D. Bugbee, The RE-JOIN Consortium Investigators, Martin K. Lotz

**Author notes:** Corresponding author: Martin Lotz, Scripps Research Institute, 10550 North Torrey Pines Road, La Jolla, CA 92037.

## Abstract

Osteoarthritis (OA) is defined by progressive loss of articular cartilage, but how chondrocytes change as the disease advances is unclear because most studies have focused on end-stage tissue. We profiled human knee articular cartilage across the full range of macroscopic OA grade, from young healthy to aged severe OA donors, by bulk RNA sequencing of 92 knees, single-nucleus RNA sequencing of 108,600 chondrocyte nuclei from 30 donors and single-nucleus multiome sequencing of 24 of these donors, treating grade as a continuous variable. With increasing grade, cartilage increased extracellular matrix (ECM) synthesis and remodeling genes, lost glucocorticoid-responsive, lipid-metabolic and antioxidant programs, and changed the splicing of a small set of ECM transcripts, led by the extra domain B (EDB) exon of fibronectin (*FN1*). Nine chondrocyte clusters were present in every disease stage and formed a continuum ordered by cartilage zone and hypertrophic differentiation, with *PRG4*+ superficial and *COL10A1*+ hypertrophic chondrocytes as the only distinct cell types at its ends and a pre-hypertrophic state between them. With grade, all clusters shared a response in which ECM production rose and homeostatic programs fell, and they differed in which of its components was strongest: ECM production in the superficial cells, the secretory pathway in the hypertrophic and *CYTL1*+ cells, and inflammatory signaling in the *ANXA1*+ and *CRISPLD2*+ cells. Inferred communication between chondrocytes was remodeled with stage. *FN1* was induced in every cluster in excess of its fibril-assembly machinery, together with the fibronectin-cleaving protease *HTRA1*, and *FN1* signaling shifted in receptor use from integrin α5β1 toward *CD44* and, less robustly, αvβ8, accompanied by a rise in TGF-β/SMAD response genes. In four early-OA donors, tenascin-C was already largely induced and the *FN1* receptor shift from α5β1 toward *CD44* was partly present, whereas the rise of *FN1* outside superficial chondrocytes, of *FGF1* and of developmental and ossification programs came with progression. Chromatin accessibility changed in the same direction across clusters, with regions linked to *FN1* opening and regions linked to *ITGA5* closing. OA progression in human cartilage therefore appears to be a change of chondrocyte state that differs in emphasis along a zonal and hypertrophic continuum and includes changes in fibronectin regulation, production, processing, and receptor engagement.

## Introduction

Osteoarthritis (OA) is the most common joint disease and a leading cause of pain and disability, affecting an estimated 595 million people worldwide in 2020 (1). OA involves every tissue of the joint (2), but progressive loss of articular cartilage remains its defining feature, and no treatment has yet been conclusively shown to modify its structural progression (3). Chondrocytes are the only resident cells of articular cartilage and are solely responsible for synthesizing and maintaining its extracellular matrix (ECM), so the changes in chondrocyte function that accompany disease progression are central to understanding how cartilage fails.

Transcriptomic studies of human cartilage have identified dysregulated ECM, PI3K-Akt, HIF-1 and FOXO pathways and a network of transcription factors that is suppressed in OA (4,5). Most of these studies compared macroscopically normal cartilage with end-stage cartilage obtained at arthroplasty, and they therefore describe the endpoint of the disease rather than changes across disease severity. Single-cell RNA sequencing has since resolved human chondrocytes into populations described as proliferative, pre-hypertrophic, hypertrophic, fibro-, regulatory and inflammatory, and several studies have proposed that OA is driven by the emergence or expansion of particular pathogenic populations (6–9). These studies have typically profiled fewer than fifteen donors and compared OA with normal cartilage or damaged with intact cartilage, so it remains unclear whether disease progression is carried by specific chondrocyte populations or by changes shared among them, and what distinguishes the response of each population.

The ECM of articular cartilage comprises collagens, proteoglycans and glycoproteins together with the enzymes that modify and degrade them and the growth factors they bind, collectively termed the matrisome (10). Chondrocytes are embedded individually in this matrix and communicate largely through the molecules they secrete into it. Growth factors such as *FGF2* are sequestered in the pericellular matrix and released on loading or injury (11), and matrix proteins themselves can signal. Fibronectin accumulates in OA cartilage (12–14), and fibronectin fragments, which are present in human OA cartilage and synovial fluid, induce matrix degradation, whereas intact fibronectin has little effect (15–17); many of these matrix signals are received through integrins (18). How communication between chondrocytes changes as OA progresses has not been examined in human cartilage across the spectrum of disease severity.

The transcriptional changes of OA are likely to be accompanied by changes in chromatin. Bulk chromatin accessibility profiling of human knee and hip cartilage has mapped chondrocyte regulatory elements and identified accessibility changes associated with OA (19,20), but it is not known whether such changes are shared among chondrocyte populations or confined to some of them, or how they relate to the transcriptional response across the spectrum of disease severity.

Here we addressed these questions by profiling human knee articular cartilage across the full range of macroscopic OA grade (Outerbridge-based grade 0-4) (21) at three levels: bulk RNA sequencing of 92 knees, single-nucleus RNA sequencing of 108,600 chondrocyte nuclei from 30 donors, and single-nucleus multiome sequencing (paired chromatin accessibility and RNA) of 24 of these donors. Treating grade as a continuous variable rather than comparing normal with end-stage tissue, we asked what distinguishes the chondrocyte populations, which programs change with OA progression in the tissue and in each population, how communication between chondrocytes and their matrix changes, and whether these changes are reflected in chromatin. Our data suggest that OA progression is carried largely by a change of state in which all chondrocyte populations take part, with differing emphasis, and that it includes changes in fibronectin transcription, splice forms and signaling as a putative intercellular signaling molecule.

## Results

### Increasing macroscopic OA grade is associated with ECM remodeling and loss of homeostatic programs in cartilage

To define how the cartilage transcriptome changes as OA advances, we first profiled whole cartilage across the full range of macroscopic disease. Bulk RNA-seq of articular cartilage from 92 knees spanning the OA macroscopic knee grade spectrum (Outerbridge-based grade 0-4) separated samples along PC1, which accounted for 41.2% of the variance among the 500 most variable genes (Figure 1a). Macroscopic knee grade was the strongest correlate of this axis (R² = 0.36). Tissue source and post-mortem interval also correlated with it, but both differ between grade 4 knees, nearly all obtained at arthroplasty, and lower-grade knees from organ donors, so their contributions cannot be distinguished from those of end-stage disease.

**Figure 1.**
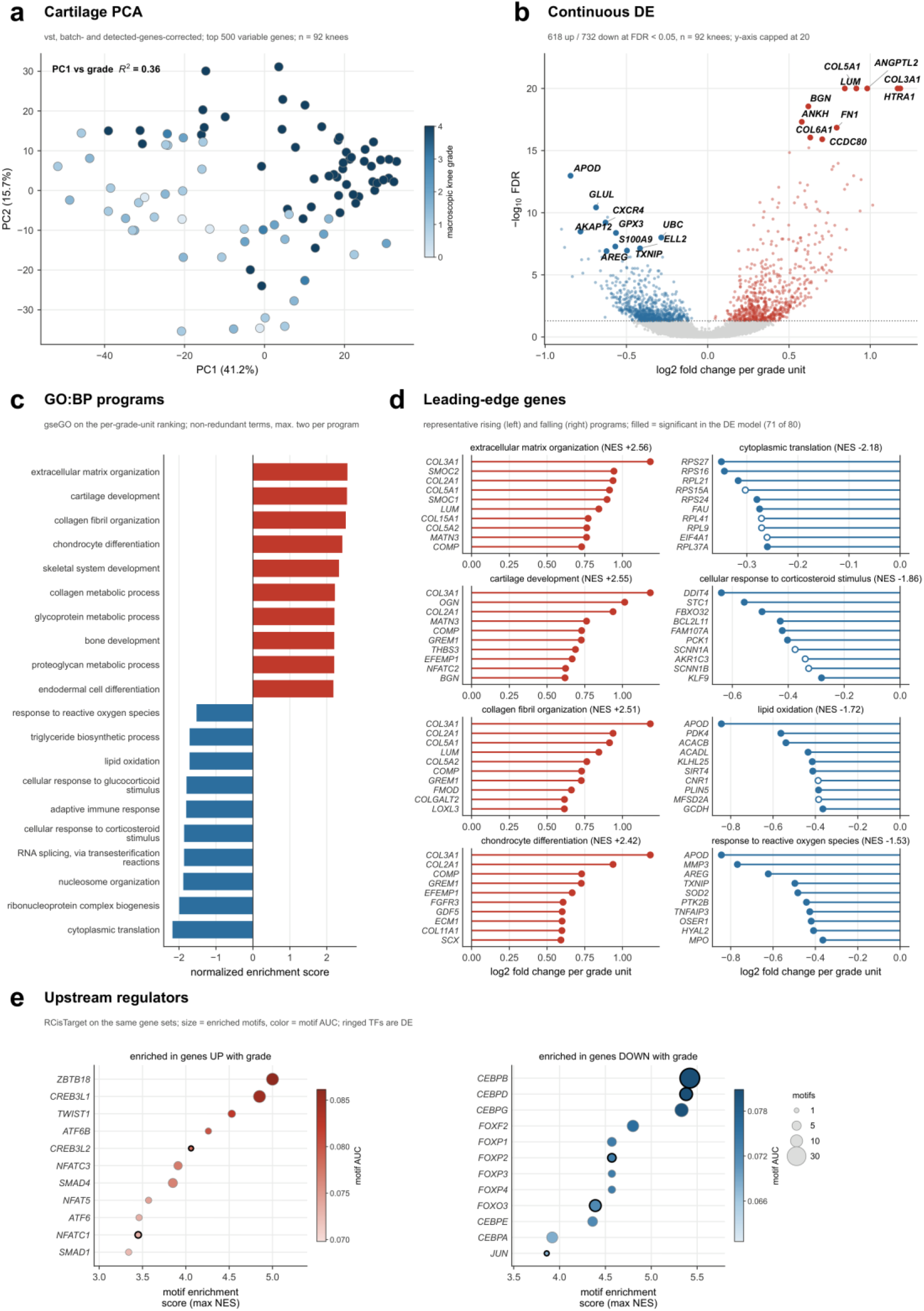
Bulk RNA sequencing of human articular cartilage across the OA severity spectrum. **(a)** Principal component analysis of 92 knees, colored by macroscopic knee grade; PC1 accounts for 41.2% of the variance among the 500 most variable genes and correlates with grade (R² = 0.36). **(b)** Genes changing with continuous grade; 618 rise and 732 fall at adjusted p < 0.05, and the most significant in each direction are labeled. The y axis is capped at 20. **(c)** GO biological processes enriched on the same ranking, ten non-redundant terms in each direction (rising, the highest normalized enrichment scores among terms of at least 30 genes; falling, selected two per biological program), as normalized enrichment score. **(d)** Leading-edge genes of four rising and four falling processes; filled points are also significant in the gene-level analysis, 71 of the 80 shown. **(e)** Transcription factor motifs enriched within 10 kb of the transcription start sites of the genes rising and falling with grade (RcisTarget); bubble size is the number of enriched motifs, fill is motif AUC, and a black ring marks a transcription factor that is itself differentially expressed with grade.

Treating grade as a continuous variable, 1,350 of 17,951 expressed genes were differentially expressed (DE) at an adjusted p value < 0.05 (618 upregulated, 732 downregulated) (Figure 1b). Among the most significantly upregulated genes were matrix and matrix-associated genes (*HTRA1*, *ANGPTL2*, *COL3A1*, *FN1*, *BGN*), and among the most significantly downregulated were genes of chondrocyte homeostasis and antioxidant defense (*APOD*, *GLUL*, *GPX3*) and the stress-responsive *TXNIP*. The direction of the signature was robust to RNA quality covariates and to removing low-quality libraries (90% to 100% of genes kept their direction, and 49% to 87% remained significant; fold-change ρ = 0.93 to 0.99) (Figure S1a-b). Comparing early OA (grade 1.5-2, n = 15) with Normal knees (grade 0-1, n = 23) did not identify an early-OA signature: 287 of the 320 differing genes formed one module of low-level, largely tissue-inappropriate transcripts that tracked library complexity (ρ = 0.88 with detected genes) and were largely absent (fewer than 5% of module genes expressed) from 10 of the 15 early-OA knees but from only one of 23 Normal knees, suggesting a technical rather than a disease-related difference (Figure S1c-d). With the module as a covariate, only *CDC20*, detected in nine knees, still differed, whereas *C4A*, a secreted gene among the uncorrected early-OA differences, did not (Figure S1c).

The grade signature was reproduced in direction in two public cartilage datasets and was enriched at OA risk loci (Figure S2). In an earlier cartilage cohort from our group (4), 80% of the 1,109 grade-associated genes measured changed in the same direction (53% also significant), and in paired lesioned and preserved cartilage from the RAAK study (22), 65% of 1,093 (14% significant); *FN1*, *HTRA1*, *ANGPTL2*, *COL5A1*, *IGFBP1* and *IL11* replicated in both (Figure S2a-b). Of the 77 high-confidence OA effector genes from genome-wide association (23), 72 were measured here and 14 changed with grade, and these 14 genes fell at 14 different risk loci out of 57, whereas only about 5.5 would be expected by chance (p = 8×10⁻⁴). However, the effector genes were no more likely to change with grade than the other genes at the same risk loci (p = 0.81) (Figure S2c-d). Thus, OA risk loci as a whole are enriched for genes that change with grade, but this does not single out the gene prioritized as causal at each locus. Finally, the grade response was compatible with a linear change: when the grades were compared as separate groups instead of as a continuous trend, no gene departed significantly from a straight-line change, for example by rising early and then leveling off (Figure S2e).

Gene set enrichment analysis identified 370 gene ontology (GO) biological processes associated with grade (176 upregulated, 194 downregulated) (Figure 1c). Upregulated processes involved ECM organization, collagen fibril organization, cartilage development and chondrocyte differentiation. Downregulated processes included translation-related terms (cytoplasmic translation, ribonucleoprotein complex biogenesis, RNA splicing; ribosome biogenesis in Figure S3) and stress resilience programs: 16 lipid and fatty acid metabolism processes spanning both oxidation and biosynthesis (lipid oxidation, NES = −1.72), cellular response to corticosteroid stimulus (NES = −1.86) and response to reactive oxygen species (NES = −1.53). With increasing grade, the cartilage transcriptome thus shifted toward ECM synthesis and remodeling, while stress resilience programs declined. The translation-related terms reflected a lower share of ribosomal-protein and translation-machinery transcripts in grade 4 libraries, which we interpret as a library-level feature of end-stage cartilage rather than as evidence of altered protein synthesis (Figure S3).

Leading-edge genes (Figure 1d) of the upregulated programs were fibrillar collagens and cartilage matrix constituents (*COL3A1*, *COL2A1*, *COL11A1*, *COMP*, *FMOD*) together with the chondrogenic regulators *FGFR3*, *GDF5* and *SCX*; those of the declining programs were ribosomal proteins and glucocorticoid-responsive (*DDIT4*, *KLF9*), lipid-oxidative (*PDK4*, *ACADL*) and antioxidant and redox-regulating (*SOD2*, *TXNIP*) genes.

Among the motifs enriched near regulated genes (top factors in Figure 1e; *XBP1*, *ATF4* and *NR3C1* rank below those shown), motifs near upregulated genes included those of *CREB3*/*ATF6* family members (*CREB3L2*, *ATF6*), *XBP1*, *ZBTB18*, *TWIST1* and NFAT and SMAD factors, and those near downregulated genes included those of C/EBP factors, *ATF4*, the glucocorticoid receptor *NR3C1* and FOXO/FOXP factors; *CREB3L2* and *NFATC1* were themselves upregulated, and *CEBPB*, *CEBPD* and *FOXO3* downregulated. The induced matrix program may therefore be regulated by *CREB3*/*ATF6* family factors, including the ER stress transducers *CREB3L2*, *ATF6* and *XBP1*, together with SMAD and NFAT, whereas loss of C/EBP, *ATF4*, FOXO and glucocorticoid receptor activity may underlie the decline of homeostatic programs.

Macroscopic Outerbridge grade describes the whole joint cartilage surface, whereas the Osteoarthritis Research Society International (OARSI) score measures the histological changes of the cartilage sampled (24). With the OARSI score in place of grade (91 knees), 660 genes changed (baseMean ≥ 10; 1,346 for grade under the same filter) (Figure S4); 545 of them also changed with grade, none in the opposite direction, and fold changes and pathway enrichment were closely correlated (r = 0.95 and 0.99) (Figure S5a-f). Processes associated with grade alone included the lipid-oxidation, corticosteroid-response and reactive-oxygen-species terms of Figure 1c, which fell with the OARSI score as well but did not reach adjusted p < 0.05 (adjusted p 0.07 to 0.12), and many immune terms, which may reflect the arthroplasty source of grade 4 tissue rather than cartilage damage. The two measures were correlated (ρ = 0.77) but diverged in arthroplasty knees, all of grade 4, whose OARSI scores ranged from 1 to 6 (Figure S5g), consistent with a small biopsy representing one standardized site rather than the whole joint. As expected for end-stage tissue, whose scores were concentrated at the severe end (41 of 47 knees scored 3 to 5), no gene changed robustly with the OARSI score within these 47 knees (Figure S5h-i). The OARSI score therefore recapitulated much of the grade-associated signature.

### A small set of matrix transcripts, led by the fibronectin EDB exon, changes its splicing with grade

Because ECM remodeling led the grade signature and many ECM proteins are made as functionally distinct splice isoforms, we next asked whether OA also changes which isoforms cartilage expresses. Of 3,657 alternative splicing events tested (Figure 2a), 12 events in 11 genes changed robustly with grade. Four were in ECM genes: the extra domain B (EDB) exon of *FN1*, the most statistically significant event genome-wide; a 5′ non-coding exon of *DCN* (ΔPSI −0.42 from grade 0 to grade 4); and two exons of *ABI3BP* (Figure 2a; one shown in Figure 2b). The others were exons of *SRSF3*, *BCLAF1*, *AKAP9*, *RPS9* and *RPS27* and retained introns of *MT1E*, *MT2A* and *ADM*. Testing the 309 matrisome events separately added an in-frame exon of *PRG4* (ΔPSI −0.57), events in *COL6A3*, *COL1A2*, *COLEC12* and *EGLN1*, and a shift within the *FN1* IIICS region (Figure 2a-b, Figure S6b). Apparent mutually exclusive exon events in *COL1A2*, *COL2A1*, *COL3A1* and *FN1* paired two constitutive exons and were excluded (Figure 2a, Figure S6b, d). All four ECM events survived a relaxed coverage filter, whereas a pooled-read model gave 15 to 46 times more calls (Figure S6a).

**Figure 2.**
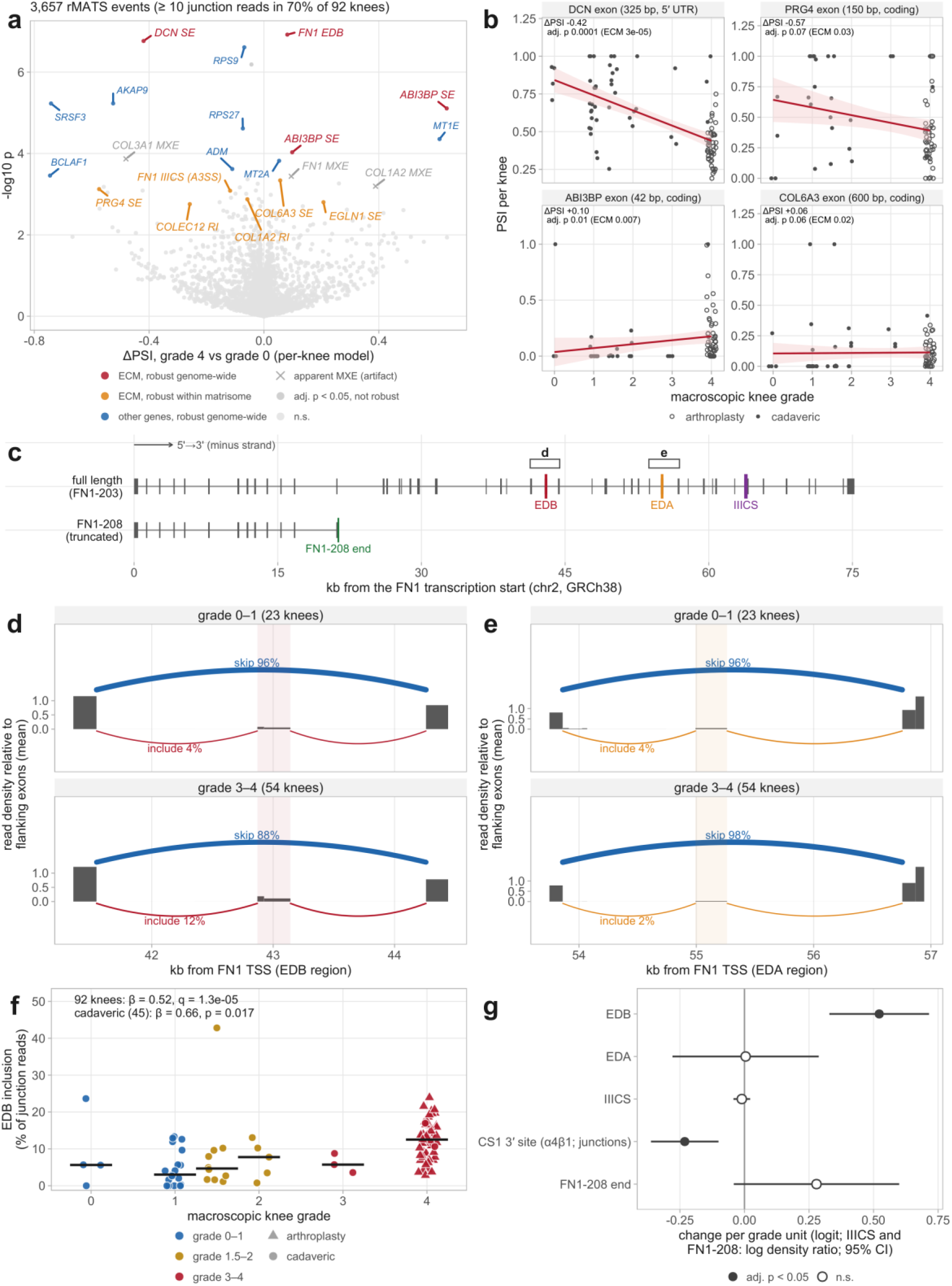
A small set of matrix transcripts, led by the fibronectin EDB exon, changes its splicing with grade. Bulk cartilage RNA-seq of the 92 knees analyzed for differential expression. **(a)** Alternative splicing events (rMATS: skipped and mutually exclusive exons, alternative 5′ and 3′ splice sites, retained introns) with at least 10 junction reads in 70% of knees, a mean PSI between 0.02 and 0.98 and a PSI standard deviation above 0.02 (3,657 events), tested per knee as logit inclusion against grade with the differential expression covariates; ΔPSI is the fitted change from grade 0 to grade 4. Robust events have adjusted p < 0.05 (Benjamini-Hochberg within each event type), |ΔPSI| ≥ 0.05 and the same sign at p < 0.05 with library strandedness as an additional covariate and after excluding the libraries flagged for low quality. Red, matrix (matrisome) genes robust genome-wide; orange, matrix genes robust when the false discovery rate is computed within the 309 matrisome events; blue, other robust events; crosses, apparent mutually exclusive exon events that pair two constitutive exons (Figure S6d). Dark gray, adjusted p < 0.05 but not robust. **(b)** Inclusion per knee of matrix exons whose splicing changed with grade (filled, cadaveric; open, arthroplasty), with ΔPSI and the adjusted p computed genome-wide and within matrisome events (ECM); lines are linear fits for display. **(c)** *FN1* gene model (GENCODE v44), drawn 5′ to 3′: the full-length transcript *FN1*-203, which contains the extra domain B (EDB) and A (EDA) exons and the complete IIICS region, and the truncated *FN1-208* transcript (ENST00000426059) with its unique 3′ end. Boxes mark the regions shown in **(d)** and **(e)**. **(d)** Sashimi-style view of the EDB region in the bulk cartilage RNA-seq of grade 0-1 (23) and grade 3-4 (54) knees: read density in each exonic (dark) and intronic (light) segment relative to the two flanking constitutive exons, mean across knees, and junction arcs for skipping (above) and inclusion (below) of EDB, labeled with the median share of junction reads; arc width is proportional to that share. **(e)** As in **(d)**, for EDA. **(f)** EDB inclusion in each of the 92 knees analyzed for differential expression, as the share of junction reads that include the exon, against macroscopic knee grade; triangles are arthroplasty and circles cadaveric knees, bars are medians. β is the change in logit inclusion per grade unit, adjusted for batch and the number of detected genes as in the differential expression analysis, for all knees (q, Benjamini-Hochberg across the 15 targeted measures: the *FN1*, *FGFR1-3*, *CD44* and *FGF1* splice measures, CS1 read density and the *FN1-208* intron control) and for the 45 cadaveric knees. **(g)** Change per grade unit in inclusion of EDB, EDA and the IIICS region, in use of the 3′ splice site that retains the 25 codons of the α4β1-binding CS1 segment (junction reads; adjusted p within matrisome events), and in the unique 3′ end of *FN1-208*, adjusted as in **(f)**, in all 92 knees (95% confidence intervals; filled, adjusted p < 0.05). IIICS and the *FN1-208* end are log ratios of read density to that of adjacent constitutive exons rather than logits.

Within *FN1* (Figure 2c), EDB inclusion rose from a median of 4% of *FN1* junction reads at this exon in grade 0-1 knees to 12% in grade 3-4 knees (adjusted p = 1.3×10⁻⁵; Figure 2d, f-g), robust to library quality and present among cadaveric knees (nominal p = 0.017) (Figure 2f) and in an exon-level model (Figure S6c), in agreement with the accumulation of EDB-containing fibronectin in OA cartilage (25). Extra domain A (EDA), the isoform associated with fibrosis, stayed low and unchanged (Figure 2e, g), as did total IIICS inclusion, but within IIICS splicing shifted away from the site that retains the integrin α4β1-binding CS1 segment (ΔPSI −0.12) (Figure 2a, g). The truncated *FN1-208* transcript encoding migration-stimulating factor, increased in lesioned OA cartilage (26), was rare and unchanged with grade (Figure 2c, g).

Thus, *FN1* transcripts shift with grade toward the EDB isoform associated with chondrogenic differentiation and away from the α4β1-binding CS1 segment, without the rise in EDA that marks fibrotic matrix.

### Nine chondrocyte clusters are detected across the OA severity spectrum

Bulk tissue cannot show which chondrocytes carry these changes, so we next profiled cartilage at single-nucleus resolution. Single-nucleus RNA sequencing (snRNA-seq) of medial femoral condyle cartilage from 30 donors (mean age 56.7 ± 21.1 years; Outerbridge-based grade 0-4, Figure S7a-b) yielded 108,600 chondrocyte nuclei in nine clusters, c0–c8 (Figure 3a), ranging from 40.4% (c0) to 1.2% (c8) of nuclei (Figure 3b). Each cluster contained nuclei from at least 26 donors, no donor contributed more than 12.2% of any cluster (Figure S7c), and all nine clusters were present in every disease stage (Figure S8a).

**Figure 3.**
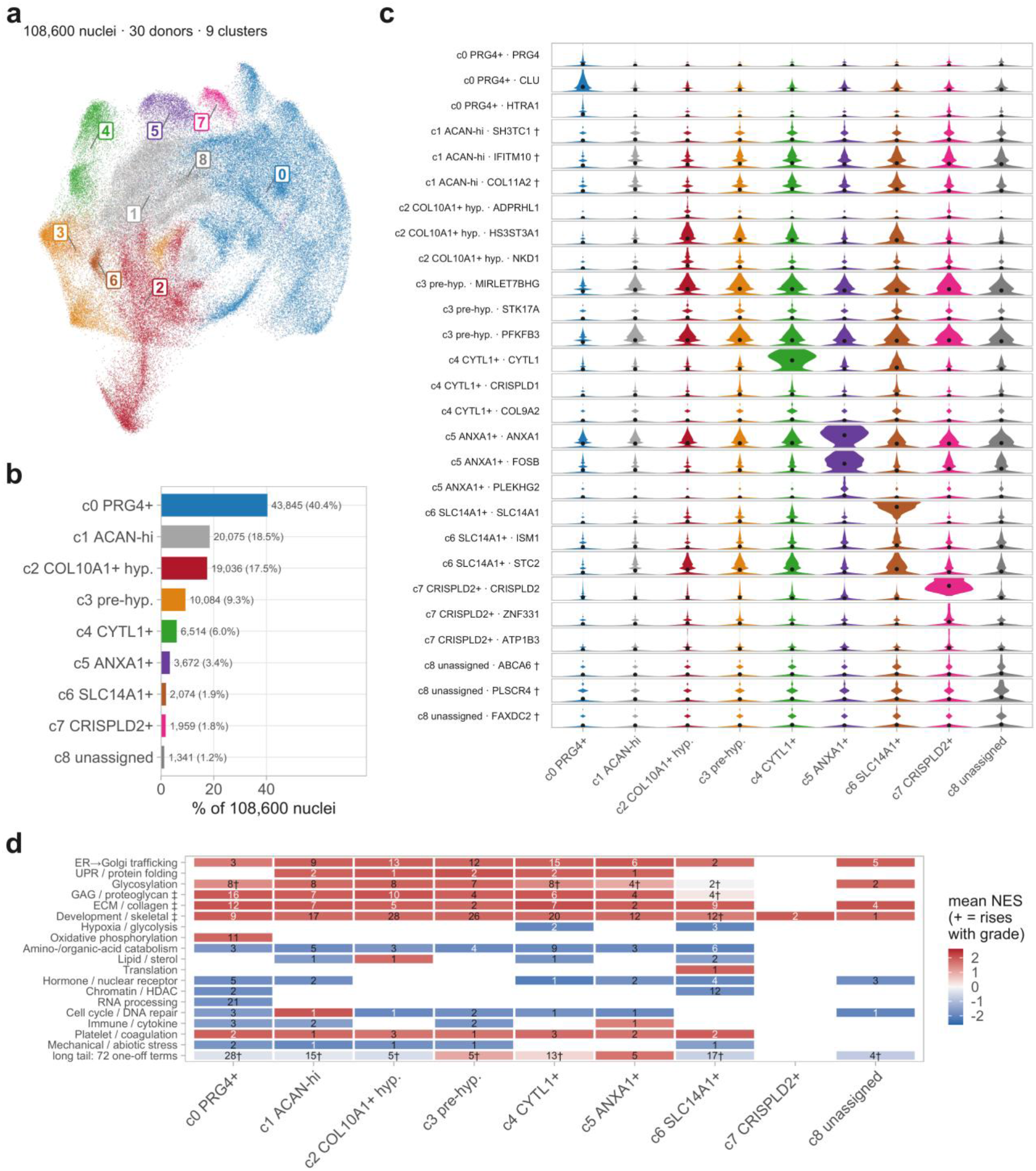
Nine chondrocyte clusters in human knee articular cartilage. **(a)** UMAP of 108,600 chondrocyte nuclei from 30 donors, clustered into nine clusters and labeled at cluster medians. **(b)** Nuclei per cluster, ranging from 40.4% (c0) to 1.2% (c8). **(c)** Per-nucleus expression of three markers per cluster; one row per gene, one violin per cluster, black dot the mean, each row on its own scale. † marks genes that are the best available for c1 and c8 rather than specific to them; the c3 genes shown are likewise only weakly enriched (see text). **(d)** Grade-associated GO biological process and Reactome terms grouped onto 18 keyword themes, with terms matching no theme and themes with few terms pooled in the bottom row; the number in each tile is how many terms are significant there and the fill is their mean normalized enrichment score. † marks tiles containing terms that change in both directions, so that the mean understates both; ‡ marks matrix-related themes, whose enrichment depends in part on the exclusion of matrix genes from the capture-burden covariate.

Six clusters had specific markers (Figure 3c): c0 *PRG4*/*CLU*, c2 *ADPRHL1*/*HS3ST3A1*, c4 *CYTL1*/*CRISPLD1*, c5 *ANXA1*/*FOSB*, c6 *SLC14A1*/*ISM1* and c7 *CRISPLD2*/*ZNF331*. c1, c3 and c8 had only weakly enriched genes (c1 *SH3TC1*, with high *COL2A1* and *ACAN*; c3 *MIRLET7BHG*/*STK17A*; c8 *ABCA6*/*PLSCR4*), each detected in 5 to 11 percentage points more nuclei than in other clusters; the genes most enriched in c3 were long-gene loci (*GPC5*, *FGF14*). Marker enrichment (Figure S9g) linked c0 to ECM organization, c4 to collagen biosynthesis, c5 and c7 to TNFα/NF-κB signaling, c2 to membrane transport and hypoxia, and c6 to skeletal system development; c3 had no enriched term, and the terms for c1 (collagen and ECM) and c8 (hormone and calcium response) each rested on two to four genes.

### Chondrocyte subsets form a continuum ordered by zone and hypertrophy, with distinct cell types at either end

The nine clusters could represent distinct cell types or interchangeable states of one cell type, so before examining how they change with disease we asked what defines each of them. To separate identity from the disease response, we first defined subset identity in the 12 healthy donors (grade 0-1) from markers, recurring programs, cross-donor recognition (whether a classifier trained on 11 of the healthy donors identified the cluster in the twelfth), chromatin, published populations and zonal signatures, checking every result in all 30 donors (Figure 4, Figure S9). On genes that distinguish superficial from deep cartilage (27), the subsets fell in the same order in every donor (median ρ = 0.92), from c0 at the superficial end through c5, c8, c7, c1, c4, c3 and c6 to c2 at the deep end (Figure 4a). A second axis, defined by *RUNX2*, *COL10A1* and *IBSP*, separated c2 from all other subsets (highest in c2 in 11 of 12 donors), with c3 and c6 intermediate. The subsets did not show a clear transcriptional separation: silhouette widths were near zero and the number of clusters depended on resolution (Figure S9).

**Figure 4.**
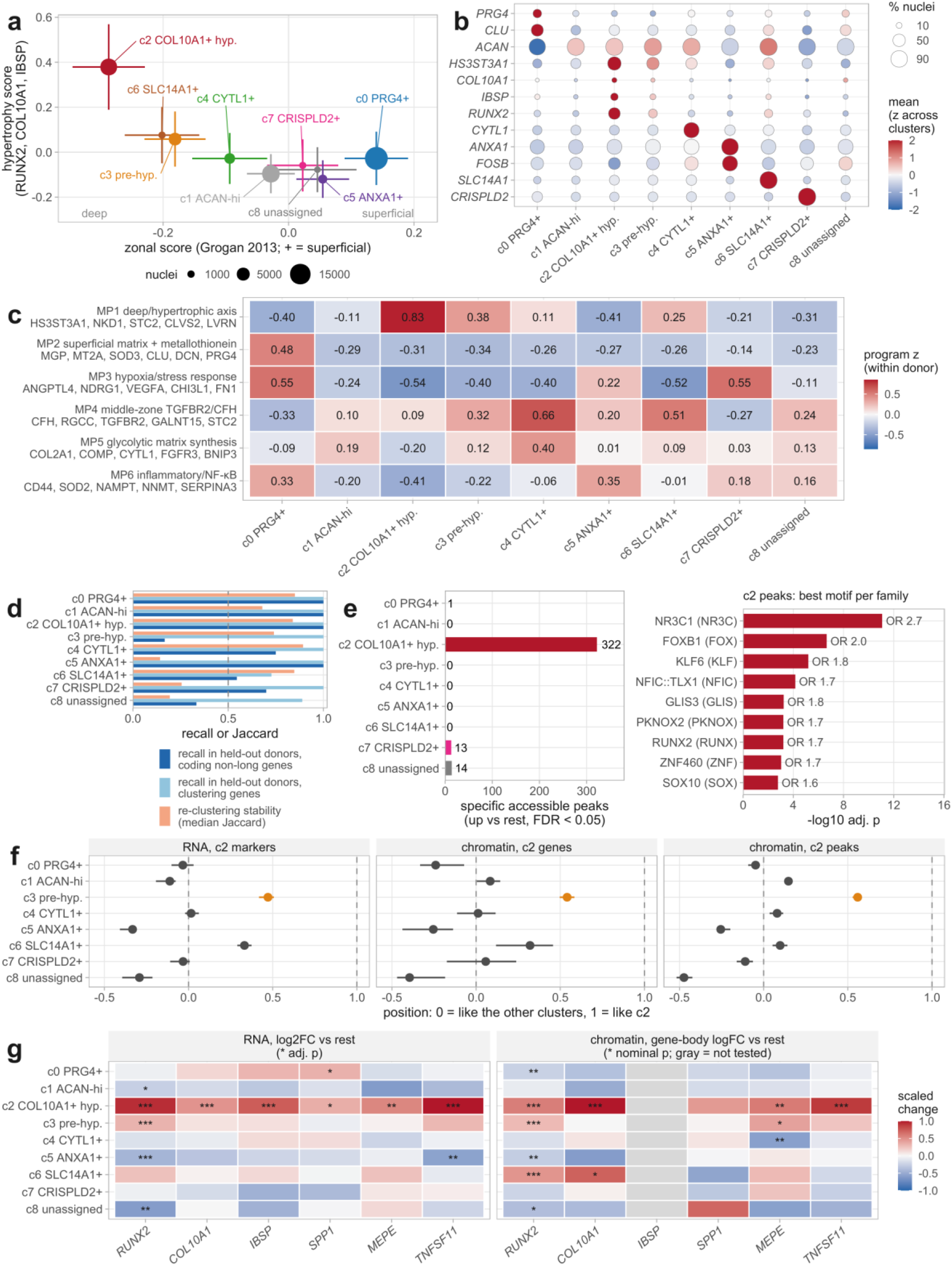
Chondrocyte subsets form a continuum ordered by zone and hypertrophy, with distinct cell types at either end. All panels use the 12 donors with healthy cartilage (macroscopic grade 0-1) unless stated; clusters are labeled with the names used in the text (c0 *PRG4*+ superficial, c1 *ACAN*-high low-signal, c2 *COL10A1*+ hypertrophic, c3 pre-hypertrophic, c4 *CYTL1*+, c5 *ANXA1*+, c6 *SLC14A1*+, c7 *CRISPLD2*+, c8 unassigned). **(a)** Zonal score (genes distinguishing superficial from deep human cartilage, ref. 27; positive = superficial) against a hypertrophy score (*RUNX2*, *COL10A1*, *IBSP*; *RUNX2* and *IBSP* are also among the deep-zone genes of the zonal score); mean ± standard error of donor means, point size proportional to the number of nuclei. **(b)** Expression of the genes used to name the clusters and of hypertrophy markers; dot size is the percentage of nuclei expressing the gene and color the mean expression scaled across clusters. **(c)** Six gene programs that recur across donors (non-negative matrix factorization within each donor; programs found in at least four of 12 donors, protein-coding genes excluding long genes); program scores z-scored within donor and averaged over donors. **(d)** Cross-donor recall of each cluster’s pseudobulk profile, with each donor in turn classified against the other donors (leave-one-donor-out, nearest-centroid assignment after centering each donor) using the 2,000 most variable protein-coding genes excluding long genes or the 956 features used for clustering, and stability on reclustering (median best-match Jaccard index over ten subsamples of 80% of all 108,600 nuclei from the 30 donors at the original resolution); dashed line, 0.5. **(e)** Number of regions more accessible in each cluster than in the others (donor-blocked pseudobulk of the single-nucleus multiome, adjusted p < 0.05), and motif enrichment among the 322 c2 regions against all tested accessible regions, stratified by GC content (Cochran-Mantel-Haenszel) (best motif per family; odds ratio shown). **(f)** Position of each cluster between the other clusters (0) and c2 (1), for the RNA of c2 marker genes (log2 fold change > 1) in the multiome nuclei, the gene-body accessibility of the same genes and the accessibility of the 539 regions more accessible in c2 at nominal p < 0.001; median and 95% confidence interval from 1,000 bootstrap resamples of genes or regions; c3 in orange. **(g)** Hypertrophy-associated genes: RNA log2 fold change of each cluster against the others (donor-blocked; stars, adjusted p) and gene-body accessibility log fold change (stars, nominal p); gray, not tested. * p < 0.05, ** p < 0.01, *** p < 0.001.

Only the subsets at the ends of these axes were distinct by every expression-based criterion, and c2 also by chromatin. c0 expressed *PRG4*, *CLU*, *CHI3L1* and *ANGPTL4* (Figure 4b-c), carried a superficial matrix and metallothionein program highest in c0 in all 12 donors (Figure 4c), matched the FC-2 population of an earlier OA cartilage atlas (8), and was recognized in every donor in cross-donor tests (Figure 4d). c2 expressed *RUNX2*, *COL10A1*, *IBSP*, *MEPE* and *TNFSF11*, headed a deep program in all 12 donors, matched published hypertrophic chondrocytes (8,9), and was the only subset with a substantial set of accessible regions of its own (322 regions, enriched for *RUNX2* and glucocorticoid receptor motifs, compared with 0 to 14 in the other subsets); 62% of c2 marker genes had concordantly opened regions (Figure 4e, g). We therefore refer to c0 as *PRG4*+ superficial and to c2 as *COL10A1*+ hypertrophic chondrocytes.

c3 lay between c2 and the other subsets: it expressed *RUNX2* at an intermediate level (log2 fold change 0.80, compared with 2.25 in c2) but not *COL10A1*, *IBSP* or *MEPE* (Figure 4b-c, g), and its chromatin was about halfway to c2, with the *RUNX2* and *MEPE* loci partly opened but *COL10A1* closed (Figure 4f-g). We therefore refer to c3 as pre-hypertrophic chondrocytes. Because c3 was separated from c2 in only a quarter of donors in pairwise cross-donor tests and was recognized as a population in only 17% of donors (Figure 4d), we regard c2 and c3 as consecutive states on one *RUNX2*-high axis rather than as separate cell types. The other subsets were reproducible states rather than distinct types (Figure 4b-d, Figure S9). c4 (*CYTL1*) combined middle-zone and matrix-synthesis programs and was the most stable on reclustering, c5 (*ANXA1*) carried immediate-early genes and an NF-κB program, and c6 and c7 were each defined by a single, nearly exclusive gene, *SLC14A1* and *CRISPLD2* (AUROC 0.96 and 0.97). c1 and c8 had no specific gene or program (best AUROC 0.59 and 0.61). c1 behaved as a low-signal state, and c8, recognized in only a third of donors in cross-donor tests and merged with c7 on reclustering, is not supported as a distinct population. We therefore refer to c4 to c7 by their marker genes, to c1 as *ACAN*-high low-signal and to c8 as unassigned chondrocytes; the evidence for every subset is summarized in Table S1. These analyses support two relatively distinct chondrocyte populations in healthy cartilage, *PRG4*+ superficial and *COL10A1*+ hypertrophic chondrocytes, at the ends of a continuum of intermediate states ordered by zone and hypertrophy, and these subsets provide the reference for the disease responses examined below.

### OA is associated with shifts within subsets rather than major changes in subset abundance

We then asked whether OA changes the subset composition of cartilage. No cluster’s abundance correlated significantly with grade (all adjusted p values ≥ 0.48) (Figure S8b); c0 rose non-significantly (37.8% of nuclei in Normal and 43.8% in severe OA donors; ρ = 0.34). Neighborhood-level testing with Milo (28) found 71 of 3,388 neighborhoods that changed in abundance with grade (10 gaining and 61 losing), mostly drawn from one cluster (median 95% of nuclei), most often c1 and c0, a pattern reproduced in single-chemistry donors (Figure S8c-d, f). Across neighborhoods, the change in abundance with grade tracked expression of the bulk grade signature of Figure 1 (ρ = 0.50), consistent with shifts in cell state within clusters rather than the emergence or loss of entire populations (Figure S8e, g).

### Chondrocyte subsets differ in the strength of components of a shared OA response

Having defined the subsets in healthy cartilage, we next asked how transcriptomes change in each as OA advances, and whether the bulk grade signature reflects a response shared by all chondrocytes or one confined to particular subsets. With increasing grade, 4,461 genes changed in at least one subset (Figure S10a), and their pathways grouped into shared themes (Figure 3d). Much of this response was shared: 17 genes changed in every subset in the same direction, including *HTRA1*, *FGF1*, *FN1* and *LTBP2* (up) and *ABCA1* and *MET* (down), and the most widely shared pathways were epithelial-mesenchymal transition (all nine subsets), ER-to-Golgi transport and ECM organization (eight), with the unfolded protein response and TGF-β signaling in six (Figure S10a-e). All 17 genes kept their direction in every subset tested after ambient RNA correction with SoupX (29) or DecontX (30); 12 (SoupX) and 14 (DecontX) of them remained significant in all nine subsets, and 16 in at least seven with either method.

Removing predicted doublets (31) left effect sizes unchanged (Figure S11). *HTRA1* and *FN1* also rose in the bulk cohort (Figure 1b). Of the 618 genes that rose in bulk, 53% rose in at least one subset and half were highest in *PRG4*+ superficial chondrocytes, whereas 27% of the 732 genes that fell in bulk, including immune genes such as *S100A9* and *CD79A*, were barely detected in chondrocyte nuclei (below 1 count per million in every subset), so part of the falling bulk signal probably reflects cells other than chondrocytes (Figure S1g). Grouping donors as Normal (grades 0-1), early OA (grades 1.5-2) and severe OA (grades 3-4) reproduced the continuous result in every subset (Figure S12); we return to the timing of these changes below.

Cluster-specific statistical significance was strongly influenced by statistical power: of 3,165 genes significant in one cluster, 2,754 were in c0, the largest. Tested directly with a cluster-by-grade interaction, the grade response of 905 of 9,051 genes (10.0%) differed between clusters (Figure 5b), most often among the widely shared genes, and no gene or pathway changed in opposite directions in different clusters. Requiring significance within the cluster, a significant interaction and a same-direction deviation from the other clusters identified 102 cluster-distinctive associations in 100 genes, 60 in c0, 18 in c6 and 14 in c5 (Figure 5a, c).

**Figure 5.**
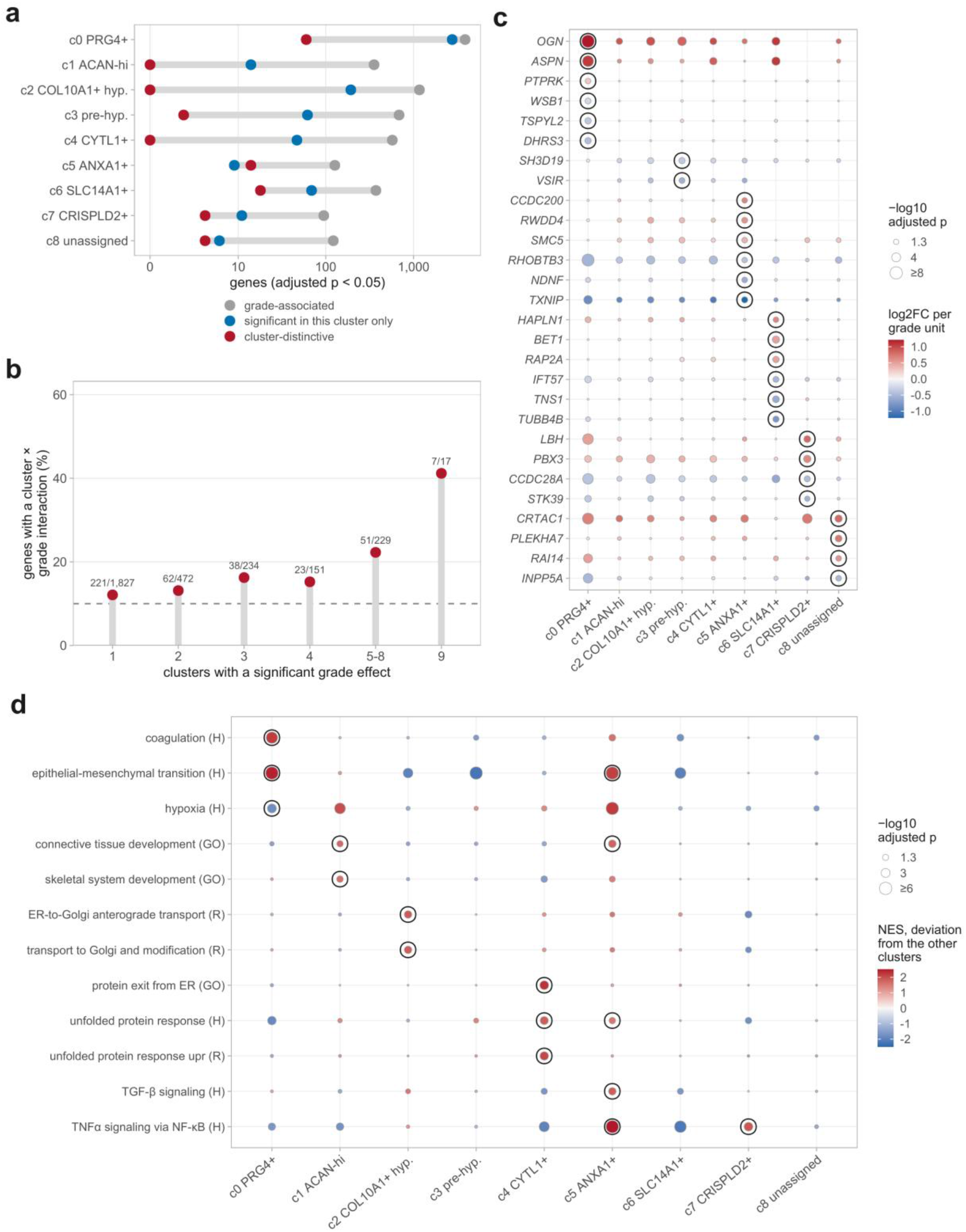
Cluster-distinctive responses to OA grade are differences in magnitude, not direction. **(a)** For each cluster, the number of genes associated with macroscopic knee grade (adjusted p < 0.05; continuous-grade pseudobulk model of Figure S10), the number significant in that cluster only, and the number that are cluster-distinctive, on a pseudo-logarithmic axis. A gene is cluster-distinctive in a cluster when it is grade-associated there, its grade response differs between clusters (cluster × grade interaction, adjusted p < 0.05) and it deviates from the mean of the other eight clusters in the same direction as its own change (adjusted p < 0.05), with its mixed-model grade slope in that cluster of the same sign. **(b)** Share of grade-associated genes with a significant cluster × grade interaction, by the number of clusters in which the gene is grade-associated; numbers are genes with an interaction over genes tested. The dashed line marks the 10.0% of all 9,051 broadly expressed genes with an interaction. The interaction was tested in a linear mixed model of variance-stabilized pseudobulk expression with technology, the ECM-excluded capture burden and cluster × grade as fixed effects and donor as a random intercept. **(c)** Cluster-distinctive genes, up to three per cluster and direction, ranked by deviation from the other clusters (c1, c2 and c4 have none): log2 fold change per grade unit in every cluster (color), with dot size showing −log10 adjusted p; the black ring marks the cluster in which the gene is distinctive. **(d)** Cluster-distinctive pathways, up to three per cluster: gene set enrichment analysis of each cluster’s deviation from the other eight (Hallmark, H; GO biological process, GO; Reactome, R), as normalized enrichment score (color) and −log10 adjusted p (size). The black ring marks pathways that deviate significantly and are also significant, in the same direction, in that cluster’s own grade analysis. No gene or pathway changed with grade in opposite directions in different clusters.

These distinctive responses changed the balance of the shared programs without reversing them (Figure 5c-d). c0 showed a stronger ECM response (*OGN*, *ASPN*, *FSTL1*, *COL6A1*; ECM organization and proteoglycan pathways; Table S1) and distinctively lost mRNA-processing and hypoxia-response genes. c4 showed the most distinctive increase in the unfolded protein response and c2 the strongest ER-to-Golgi transport, whereas the epithelial-mesenchymal transition response was attenuated in c2, c3 and c6. c1 showed stronger induction of skeletal and connective tissue development genes. TNFα/NF-κB signaling rose distinctively only in c5 and c7 (in c5 with TGF-β signaling and a stronger loss of *TXNIP*), and c6 distinctively lost the hypoxia-responsive glucose transporter *SLC2A1* (Table *S1*) and was the only cluster in which glycolysis genes fell. c7 and c8 contained too few nuclei to exclude distinctive responses.

### Each chondrocyte subset produces a distinct matrix, and OA shifts all of them toward fibronectin and small proteoglycans

Because ECM remodeling dominated the bulk grade signature and matrix genes were prominent in the shared response, we next examined which matrix each subset produces and how OA changes it. In the 12 healthy donors, all subsets devoted 2.8% to 3.3% of their transcriptome to matrisome genes (10), so they differed in matrisome composition rather than in matrisome share (Figure 6a, c). *PRG4*+ superficial chondrocytes (c0) made a glycoprotein- and small leucine-rich proteoglycan (SLRP)-rich matrix (*PRG4*, *DCN*, *MGP*, *TNC*, *COMP*) in which fibronectin was as abundant as aggrecan; *ACAN*-high chondrocytes (c1) the classical aggrecan and type II collagen matrix; *COL10A1*+ hypertrophic chondrocytes (c2) a mineralizing matrix (*COL10A1*, *IBSP*, *TNFSF11*, *P3H2*); pre-hypertrophic (c3), *CYTL1*+ (c4) and *SLC14A1*+ (c6) chondrocytes indistinguishable aggrecan-dominated matrices; *ANXA1*+ chondrocytes (c5) the highest share of ECM-affiliated transcripts; and *CRISPLD2*+ chondrocytes (c7) a glycoprotein-rich matrix with *CRISPLD2* and *COL18A1*.

**Figure 6.**
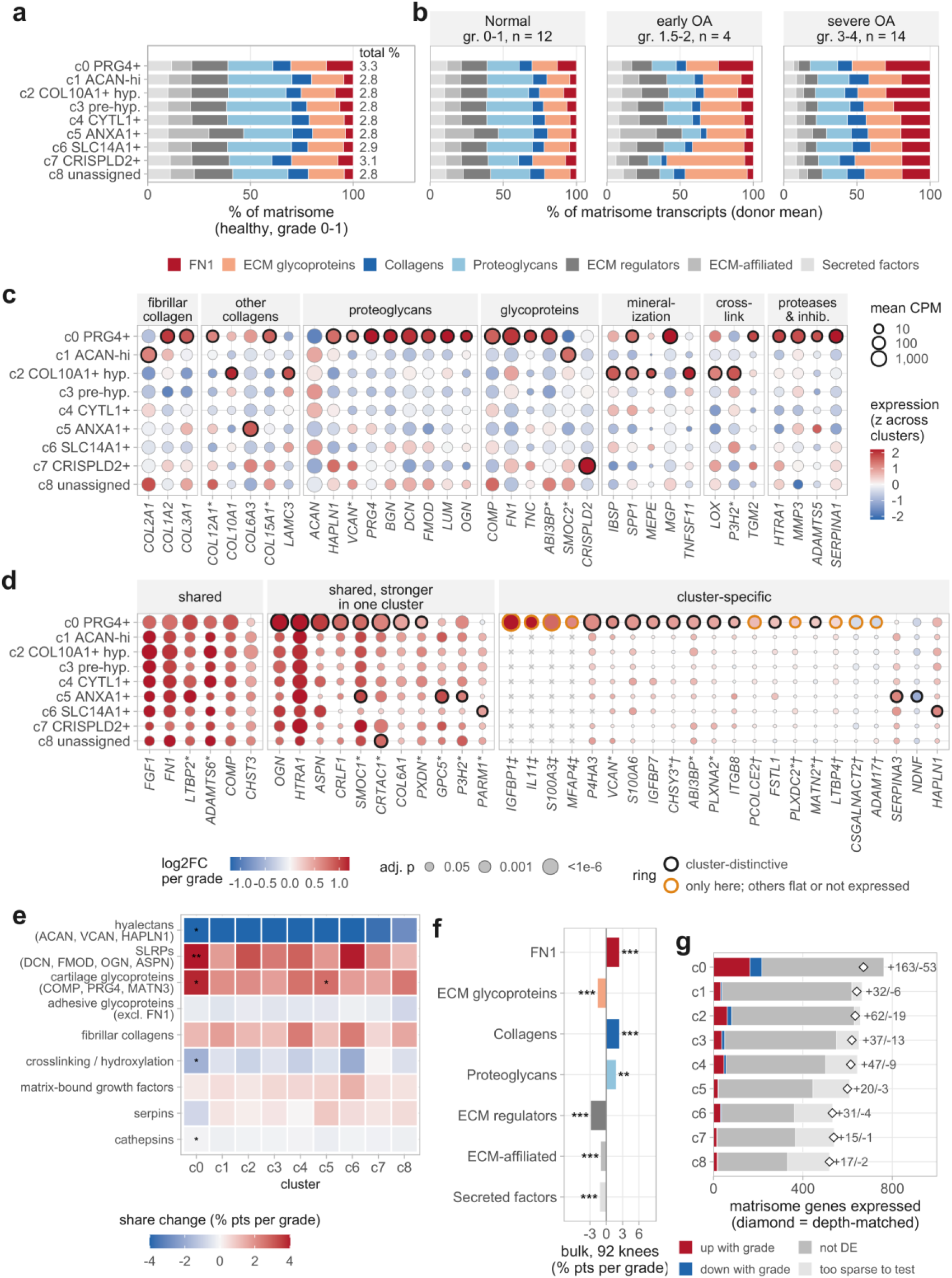
Each chondrocyte subset produces a distinct matrix, and OA shifts all of them toward fibronectin and small proteoglycans. Single-nucleus data are donor-by-cluster pseudobulks (at least 20 nuclei), with long genes (span at least 8 times the 13.8 kb background) excluded; matrisome categories follow the Naba matrisome (ref. 10) and matrix families are curated by function (Supplementary Methods). **(a)** Composition of each cluster’s matrisome transcripts in the 12 healthy donors (grade 0-1), with *FN1* shown separately from the other ECM glycoproteins; numbers give the matrisome as a percentage of all transcripts. **(b)** The same composition in Normal (grade 0-1, 12 donors), early OA (grade 1.5-2, 4 donors; two to four per cluster) and severe OA (grade 3-4, 14 donors) knees. **(c)** Matrix gene repertoire in healthy donors: dot size is mean counts per million and color the expression scaled across clusters; a black ring marks a gene specific to that cluster (donor-blocked limma, adjusted p < 0.05 and log fold change > 0.5) and an asterisk a long gene. **(d)** Shared and cluster-specific grade responses of matrisome genes: color, log2 fold change per grade unit (per-cluster pseudobulk model with technology and capture burden as covariates; genes with at least 10 counts in at least half of a cluster’s donors, and otherwise in at least three donors, refitted with the same model and size factors); dot size, adjusted p; cross, too few counts to test in that cluster. Shared, the matrisome genes significant in the most clusters; shared but stronger in one cluster, cluster-distinctive genes significant in at least three clusters; cluster-specific, the remaining cluster-distinctive genes and genes changing in one cluster only. A black ring marks a cluster-distinctive response (significant in that cluster, cluster-by-grade interaction and deviation from the other eight clusters adjusted p < 0.05, same sign; interaction adjusted across all 9,051 genes in the model, 346 of them matrisome genes) and an orange ring a gene that changes only in that cluster while every other cluster is flat (95% confidence interval including 0 and absolute log2 fold change < 0.2) or does not express it; dagger, flat in all eight other clusters; double dagger, restricted to that cluster (mean counts per million at least threefold above every other cluster, all of which have too few counts to test; Figure S13f); asterisk, long gene. **(e)** Change per grade unit in the share of nine selected matrix families, with *FN1* removed, per cluster (donor-level linear model with technology and capture burden; stars, adjusted p across all cluster-family tests); all families are shown in Figure S13. **(f)** Bulk cartilage RNA-seq: change per grade unit in each category’s share of matrisome transcripts (transcripts per million), with *FN1* shown separately as in a, across the 92 knees (share ∼ batch + detected genes + grade; stars, adjusted p). **(g)** Matrisome genes expressed in each cluster (mean counts per million of at least 1 over donors and at least three counts in at least three donors, or testable in the per-cluster grade model), divided by their change with grade in that cluster (adjusted p < 0.05); diamonds, genes expressed after thinning each cluster’s counts to the median depth of c1 to c8; numbers, genes up and down. Every matrix gene that changes in at least one cluster is shown in Figure S14. * p < 0.05, ** p < 0.01, *** p < 0.001.

With grade, the matrisome share rose in every subset and every subset shifted toward fibronectin: from Normal to severe OA, *FN1* rose from 13% to 31% of matrisome transcripts in *PRG4*+ superficial chondrocytes, from 9% to 30% in hypertrophic chondrocytes and from 4-7% to 19-25% in the other subsets (Figure 6b), consistent with the increased fibronectin content of OA cartilage, where it is found mainly in the surface zone (12,13). In early OA, the *FN1* share had already reached 24% in *PRG4*+ chondrocytes but changed by less than 4 percentage points in the other subsets (Figure 6b; two to four early-OA donors per subset). Excluding *FN1*, hyalectans fell and SLRPs and cartilage glycoproteins rose in all nine subsets, significantly in *PRG4*+ chondrocytes (hyalectans from 28% to 10%, SLRPs from 12% to 25%) and, for cartilage glycoproteins, in *ANXA1*+ chondrocytes (Figure 6e). Aggrecan expression itself did not fall but was diluted by induction of *FMOD*, *OGN*, *ASPN* and *COMP*, and *PRG4* did not change (Figure 6d, Figure S13e). *COL10A1* and *IBSP* did not rise in hypertrophic chondrocytes, so this matrix reflects subset identity and not a disease response (Figure S13e). In bulk (transcripts per million), the fibronectin, collagen, SLRP and cartilage glycoprotein shares rose with grade and the hyalectan and other ECM glycoprotein shares fell (Figure 6f, Figure S13); the SLRP, cartilage glycoprotein and hyalectan changes matched those in chondrocyte nuclei. The SLRP, collagen crosslinking and fibrillar collagen responses differed between subsets (Figure S13j-m), and matrix genes were enriched among grade-associated genes in eight of nine subsets (Figure S14d).

Most matrix responses were shared: *FN1*, *FGF1* and *LTBP2* rose in all nine subsets and *COMP*, *ADAMTS6* and *CHST3* in eight (Figure 6d). Of 26 cluster-distinctive matrix responses in 24 genes, all but one (*NDNF*, in c5) were increases and 18 were in *PRG4*+ superficial chondrocytes (Figure 6d), including *IGFBP7*, *VCAN*, *CHSY3*, *P4HA3* and *MATN2*, which rose only there, and *OGN* and *CRLF1*, which rose most steeply there; separately, *CSGALNACT2* and *ADAM17* fell only in *PRG4*+ chondrocytes, with every other subset flat (Figure 6d). No matrix gene was consistently expressed in one subset and silent in all others in more than two donors, but four were restricted to *PRG4*+ chondrocytes (at least threefold higher than in any other subset) and rose with grade (Figure 6d, Figure S13f): *IGFBP1*, essentially absent in Normal and early OA donors but expressed in 11 of 14 severe OA donors, *IL11*, *S100A3* and *MFAP4*. Elsewhere, *SERPINA3*, *GPC5*, *NDNF*, *P3H2* and *SMOC1* (*SMOC1* also in c0) responded distinctively in *ANXA1*+ chondrocytes (c5), *HAPLN1* and *PARM1* in *SLC14A1*+ chondrocytes (c6) and *CRTAC1*, as in c0, in unassigned chondrocytes (c8); no matrix gene changed in opposite directions between subsets, and no distinctive response was found in the other subsets, although power was greatest in the abundant *PRG4*+ subset. Each subset expressed 519 to 763 of the 1,111 matrisome and curated matrix-family genes, 487 of them in all nine (Figure 6g, Figure S14); *PRG4*+ chondrocytes, the most deeply sequenced, expressed the most, and 216 of their matrix genes changed with grade, compared with 16 to 81 in each other subset (Figure 6g, Figure S14).

Chondrocyte subsets thus are characterized by transcriptomes encoding ECM of distinct composition at a similar level, and OA shifts all of them toward fibronectin and SLRPs and away from hyalectans, with the earliest detectable changes in superficial chondrocytes, which also add a matrix response of their own.

### Chondrocyte communication is remodeled with OA stage, and inferred *FN1* signaling shifts its receptors

Because the matrix and the shared response consisted largely of secreted products, including *FGF1* and *FN1* (Figure S10b), which act on neighboring cells, we next asked how OA changes communication between the subsets. We inferred communication between the nine subsets at each disease stage with CellChat (32,33). Significant ligand-receptor links rose from 3,198 in Normal cartilage to 3,337 at early OA and 3,853 at severe OA, whereas summed communication probability peaked at early OA (Figure 7a, Figure S15), and the matrisome-specific method MatriCom (34) reproduced the progressive expansion (Figure S16a).

**Figure 7.**
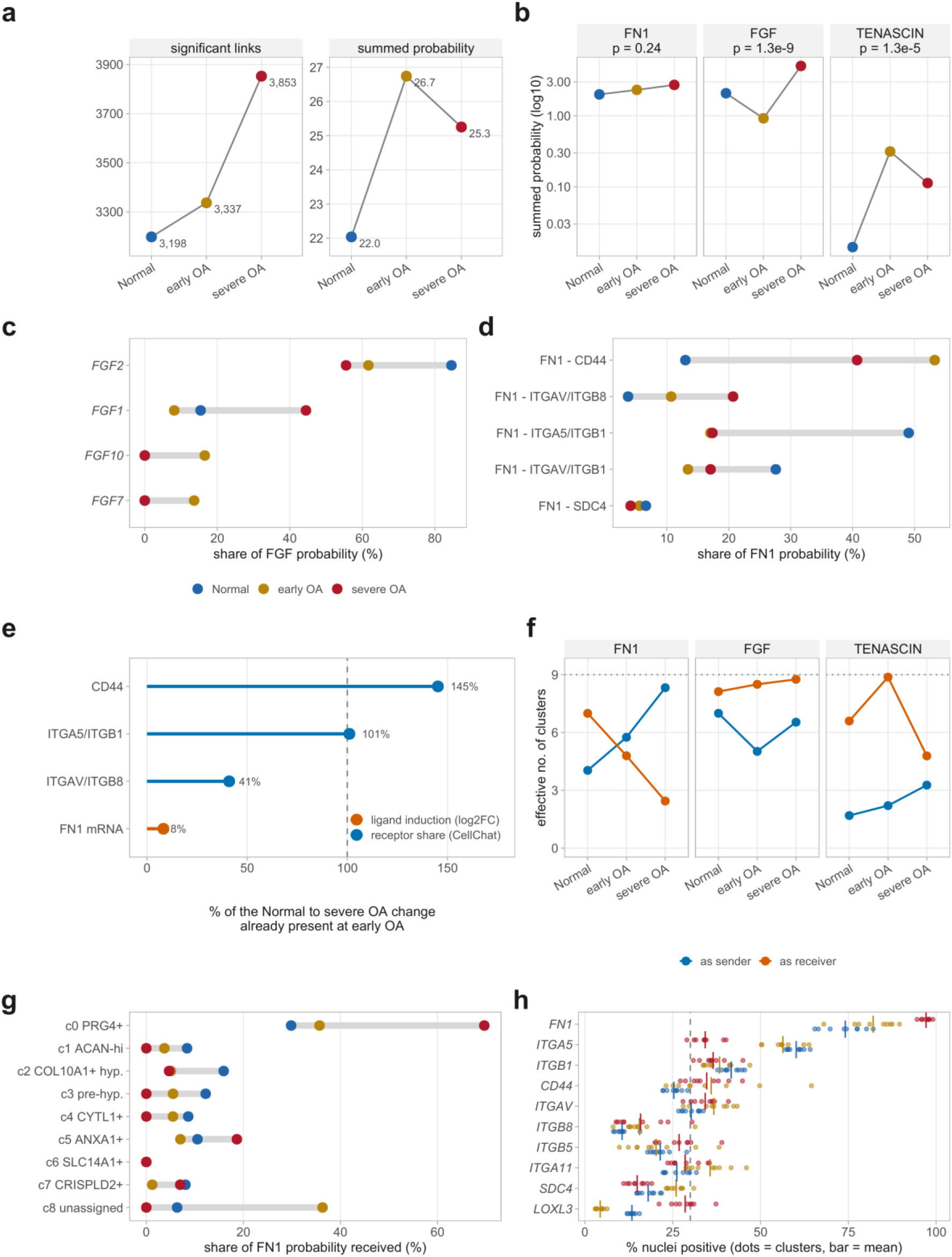
Inferred matrisome signaling between chondrocyte clusters is remodeled across OA stage, and *FN1* signaling shifts its receptors. CellChat communication probabilities (truncated mean, trim = 0.1), computed separately at each stage (Normal, grades 0-1; early OA, 1.5-2; severe OA, 3-4); color is stage throughout. **(a)** Significant ligand-receptor links and summed communication probability at each stage, shown separately because the link count rises monotonically while summed probability peaks at early OA. **(b)** Summed probability of the *FN1*, FGF and TENASCIN pathways on a log scale; strip labels give the rankNet p value for Normal against severe OA. **(c)** Each ligand’s share of FGF pathway probability. **(d)** Each receptor’s share of *FN1* probability. In c and d, gray bars span the three stages. **(e)** The percentage of each quantity’s Normal-to-severe-OA change already present at early OA, for the three *FN1* receptors whose share changed and for *FN1* mRNA (median early-OA log2 fold change against Normal over the seven clusters with at least three early-OA donors (c0 to c5, c7), as a percentage of the median severe-OA log2 fold change over all nine clusters, and the per-cluster value ranges from −3% to 51%, highest in c0); the dashed line is 100%. **(f)** Effective number of clusters sending and receiving each pathway, the exponential of the Shannon entropy of probability over the nine clusters; the dotted line marks all nine. Receiver convergence largely reflects CellChat’s significance filtering (Figure S21). **(g)** Each cluster’s share of *FN1* probability received. **(h)** MatriCom corroboration: percentage of nuclei expressing *FN1* and its receptor and gating partners in each cluster (points) and their mean (bar), with MatriCom’s 30% positive-population threshold dashed. Only the direction of change is compared between CellChat and MatriCom, not absolute magnitude.

Three matrisome pathways changed on different timescales (Figure 7b, Figure S17). Inferred TENASCIN signaling rose by early OA and remained above Normal levels (p = 1.3×10⁻⁵; here and below, paired Wilcoxon test of Normal against severe OA over sender-receiver cluster pairs, with nuclei pooled within each stage, so these p values do not account for between-donor variation). Inferred FGF signaling fell at early OA and rose between early and severe OA (Normal against severe OA, p = 1.3×10⁻⁹), with a change in ligand composition: *FGF2* carried most FGF signaling at every stage, but the *FGF1* share nearly tripled, from 15.4% in Normal cartilage to 44.5% at severe OA (Figure 7c). The donor-level *FGF1* to *FGF2* ratio rose with grade in all nine clusters (ρ = 0.57 to 0.81), whereas the *FGFR1* to *FGFR3* balance and a score of ERK feedback targets of FGF signaling did not change, although *SPRED1* and *SPRED2* fell in some clusters (Figure S18a-c). Two features may limit the effect of increased *FGF1*: its release and the sulfation of the heparan sulfate it binds. *FGF1* lacks a signal peptide and is released with *S100A13* and synaptotagmin-1 (35), of which only *S100A13* rose, modestly and in c0 alone (Figure S19a, c), and the expression balance of heparan sulfate 6-O sulfotransferases and 6-O endosulfatases, which control the sulfation that FGF-FGFR complexes require, fell in six of seven clusters, mainly through lower *HS6ST3* and higher *SULF1* (Figure S19a, b), while *SULF2* tracked *FGF1* across donors independently of grade (Figure S19c, d). *FGF18*, an *FGFR3* ligand whose recombinant form, sprifermin, has been developed as a therapeutic, rose modestly in c0 and c1, but *FGFR3* did not change (Figure S19a), and at severe OA *FGF1* was 10 to 180 times more abundant than *FGF18* (pseudobulk counts per million). In bulk, *FGFR1* and *FGFR2* were almost entirely the mesenchymal IIIc form, and their splicing, like that of *CD44* and the *FGF1* first exons, did not change with grade (Figure S19e). Total *FN1* signaling rose by 36% but not significantly in the pooled analysis (p = 0.24), although *FN1* expression rose in all nine subsets (Figure S10b).

Inferred *FN1* signaling instead showed a shift in receptor usage (Figure 7d, Figure S20): from Normal cartilage to severe OA, the share of *FN1* signaling through *ITGA5*/*ITGB1* fell from 49.1% to 17.4%, while that through *CD44* rose from 13.0% to 40.7% and through *ITGAV*/*ITGB8* from 3.7% to 20.7%. The shift from *ITGA5*/*ITGB1* toward *CD44* was already present at early OA (four donors), when *FN1* expression had reached a median of 8% of its eventual increase (−3% to 51% across the seven clusters with at least three early-OA donors), whereas *ITGAV*/*ITGB8* rose progressively (Figure 7e). *FN1* signaling also converged on fewer recipients, from 7.0 to 2.5 effective receiving subsets (Figure 7f), with c0 receiving 69.7% of *FN1* signal at severe OA compared with 29.8% in Normal cartilage (Figure 7g). In MatriCom, *FN1* was already expressed in 65.5 to 82.2% of nuclei of every subset in Normal cartilage (Figure 7h), and all 225 *FN1*-containing interactions gained by severe OA depended on expression of the partner, principally *CD44*, *LOXL3*, integrin subunits and *TNC*, rather than on *FN1* (Figure S16b-c).

Because the stage analysis pools nuclei, we repeated CellChat in each donor with identical settings (Figure S21). With the donor as the unit, the *FN1*-*ITGA5*/*ITGB1* share fell from 48% to 21% (Normal against severe OA, 12 and 14 donors; adjusted p = 6.4×10⁻⁴) and the *FN1*-*CD44* share rose from 13% to 36% (adjusted p = 6.4×10⁻⁴); the *ITGAV*/*ITGB8* share also rose (6% to 19%, adjusted p = 0.003) but less robustly. Both main shifts held in multiome-only donors and in a permutation-free expression score. In CellChat, 87% and 90% of each was already present in the four early-OA donors (nominal p = 0.045 for each; adjusted p = 0.068), but in the expression score only about half was (50% and 46%; p = 0.20 and 0.50). The *FGF1* share also rose (20% to 43%, adjusted p = 0.008), and c0 received more of the *FN1* signal (32% to 77%), although by less in the expression score (15% to 23%), in which the convergence on fewer receiving subsets was largely absent (8.2 to 7.0 effective receivers; Figure S21p). Total *FN1* signaling rose modestly (+41%, 95% confidence interval −1% to +81%, p = 0.11). Thus, as OA progressed, inferred *FN1* signaling changed less in overall magnitude than in inferred receptor usage: its reception shifted from integrin α5β1 toward *CD44* and, less robustly, αvβ8, a shift that held with the donor as the unit and, in CellChat, was largely apparent in the four early-OA donors, before most of the rise in *FN1* expression outside superficial chondrocytes.

### Tenascin-C changes by early OA, whereas developmental and ossification programs build with progression

Because the stage groups span early as well as end-stage disease, we next asked which changes are already present in early OA and which are added with progression (Figure 8). In the seven subsets with at least three early-OA donors (four donors, aged 55 to 71 years, compared with 18 to 52 years for Normal donors), 21.5% of the 8,168 changes between Normal and severe OA had reached at least half their size by early OA, 43.7% built up progressively and 34.8% appeared only late, with similar proportions in every subset and with long genes excluded (Figure 8a). Tested directly, 3.0% were established by early OA and 35.4% changed further between early and severe OA; the rest could not be resolved with three or four early-OA donors.

**Figure 8.**
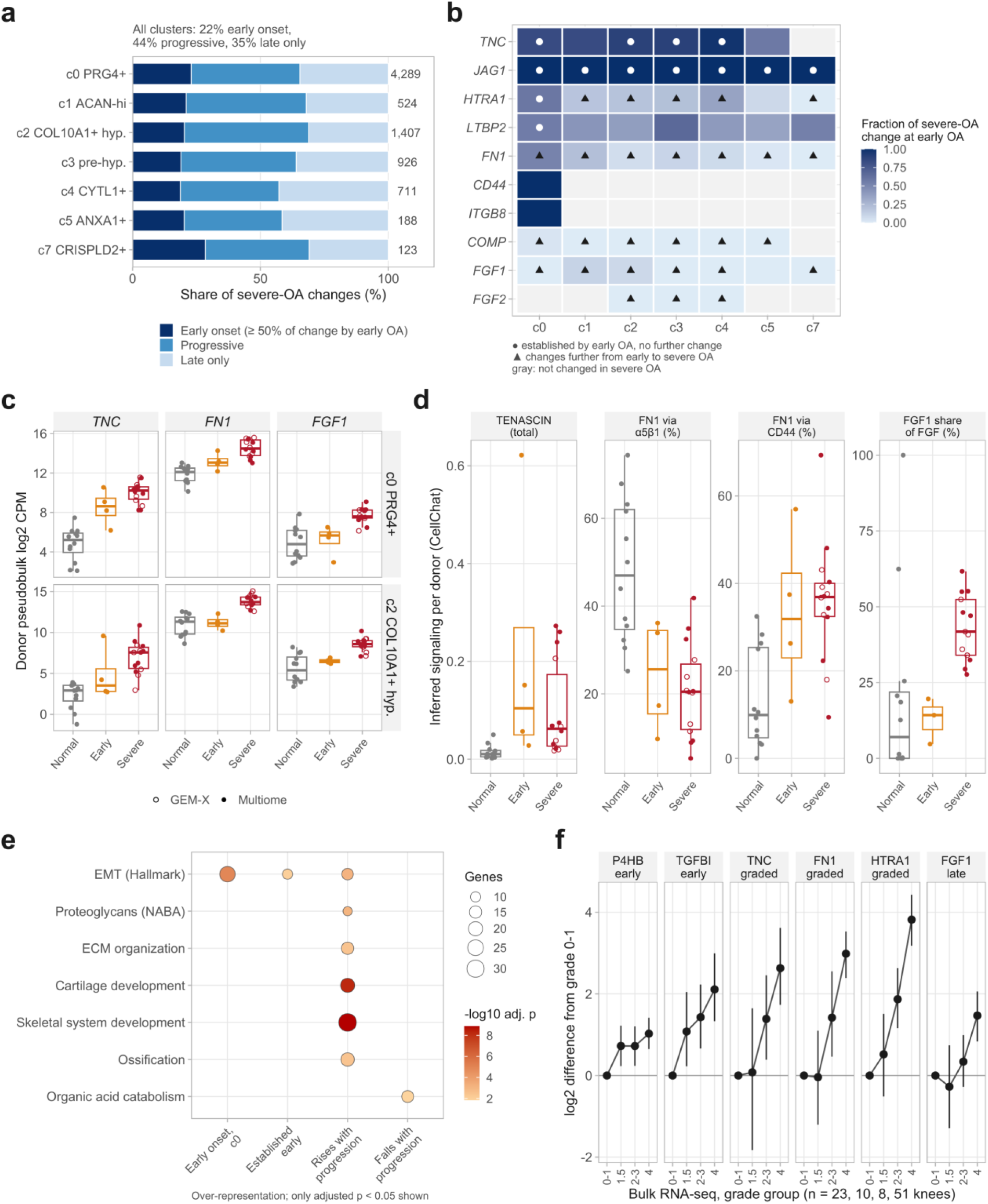
Tenascin-C changes by early OA, whereas developmental and ossification programs build with progression. Panels a-c and e use the three-stage single-nucleus pseudobulk model (Normal, grade 0-1, 12 donors; early OA, grade 1.5-2, 4 donors; severe OA, grade 3-4, 14 donors; technology and capture burden as covariates; Figure S12), in the seven clusters with at least three early-OA donors. **(a)** Genes changing between Normal and severe OA in each cluster (adjusted p < 0.05; number at right), classified by the fraction of the change present at early OA: early onset, at least 50% in the same direction; progressive, less than 50% in the same direction; late only, an early-OA change of the opposite sign. **(b)** Fraction of the severe-OA change present at early OA for selected genes (color, truncated to 0-1; gray, not changed in severe OA). Circle, established early (early OA against Normal adjusted p < 0.05 in the same direction, no change from early to severe OA); triangle, further change in the same direction from early to severe OA (adjusted p < 0.05). **(c)** Donor pseudobulk expression (log2 counts per million) of *TNC*, *FN1* and *FGF1* in *PRG4*+ superficial (c0) and *COL10A1*+ hypertrophic (c2) chondrocytes; boxes show median and interquartile range; open symbols, GEM-X donors. **(d)** Per-donor CellChat (as in Figure S21): total TENASCIN communication probability, shares of *FN1* signaling through *ITGA5*/*ITGB1* and through *CD44*, and the *FGF1* share of FGF signaling. **(e)** Over-representation of Hallmark, NABA matrisome and GO biological process gene sets among genes with early onset in c0 (rising), genes established early in at least one cluster, and genes that rise or fall further from early to severe OA in at least two clusters (background, genes tested in at least one cluster); only adjusted p < 0.05 shown; dot size, number of genes. **(f)** Bulk RNA-seq (92 knees): variance-stabilized expression adjusted for batch and detected genes, as the difference of each grade group from grade 0-1 (mean and bootstrap 95% confidence interval), with each gene’s timing class (early, at least half of the grade 4 change at grade 1.5; late, less than a quarter of it at grade 2-3; graded otherwise; mid-plateau, including the four rising overshoot genes, as defined in Methods). The four early-OA donors were aged 55 to 71 years.

Among the changes established early, *TNC* had reached 57% to 97% of its increase by early OA in c0 to c5 (log2 fold change 3.7 in c0, adjusted p = 8×10⁻⁵) without rising further (adjusted p ≥ 0.44), and the Notch ligand *JAG1* was established early in all seven subsets (Figure 8b-c). Inferred TENASCIN signaling followed the same course per donor (Wilcoxon rank-sum, nominal p = 0.009 for Normal against early OA and p = 0.56 for early against severe OA), whereas the shift in *FN1* receptor use toward *CD44* was only partly present at early OA (nominal p = 0.045; about half in the expression score; Figure 8d, Figure S21). *FN1* changed early only in *PRG4*+ superficial chondrocytes (half of its increase; adjusted p = 0.037), together with *HTRA1* and *LTBP2*; elsewhere *FN1* and *HTRA1* rose mainly between early and severe OA, as did *COMP* and *FGF1*, and the *FGF1* share of FGF signaling rose only at severe OA (Figure 8b-d).

Genes with early onset in *PRG4*+ chondrocytes and genes established early in any subset were enriched for epithelial-mesenchymal transition (EMT) genes (24 genes, adjusted p = 1.3×10⁻⁵, including *FN1*, *TNC*, *HTRA1*, *CD44*, *POSTN*, *VCAN* and *TGFBI*; and 10 genes, adjusted p = 0.0096), whereas genes that rose further with progression were enriched, beyond EMT, for skeletal system and cartilage development (adjusted p = 1.5×10⁻⁹ and 6.3×10⁻⁹), ossification, ECM organization and proteoglycans, and those that fell further for organic acid catabolism (Figure 8e); these enrichments held with long genes excluded.

In bulk cartilage grouped by grade (0-1, 1.5, 2-3 and 4; 23, 10, 8 and 51 knees; grade 2 grouped with grade 3), no gene departed from a linear trend, *TNC*, *FN1* and *HTRA1* did not detectably differ at grade 1.5 and rose from grade 2-3, and only 23 rising genes, among them collagen-processing and secretion genes (*P4HB*, *P3H2*, *CREB3L2*), had reached half their change by grade 1.5 by point estimate (Figure 8f); genes that fell only at grade 4 cannot be separated from the arthroplasty source of that tissue. Early OA was thus marked by tenascin-C and *JAG1* across subsets, a partial shift in *FN1* receptor use and, in superficial chondrocytes, an early matrix-remodeling response that included *FN1*, whereas *FN1* in the other subsets, *FGF1* and developmental and ossification programs came with progression; because *TNC* and *FN1* had not detectably changed in bulk at grade 1.5, these early changes rest on the four early-OA donors of the single-nucleus cohort.

### Chromatin accessibility changes with OA grade in the same direction across chondrocyte subsets

To determine whether these transcriptional changes are accompanied by changes in gene regulation, we measured chromatin accessibility together with RNA in the same nuclei. Single-nucleus multiome sequencing (paired ATAC and RNA) of 24 of the 30 donors yielded 81,670 chondrocyte nuclei labeled with the nine subsets (Figure 9a, Figure S22a). As in the transcriptome, the subsets were highly similar in chromatin (mean motif-activity correlation r = 0.975), differing mainly along an AP-1 axis high in c0 and c7 and low in c4 and c6 (Figure S22f).

**Figure 9.**
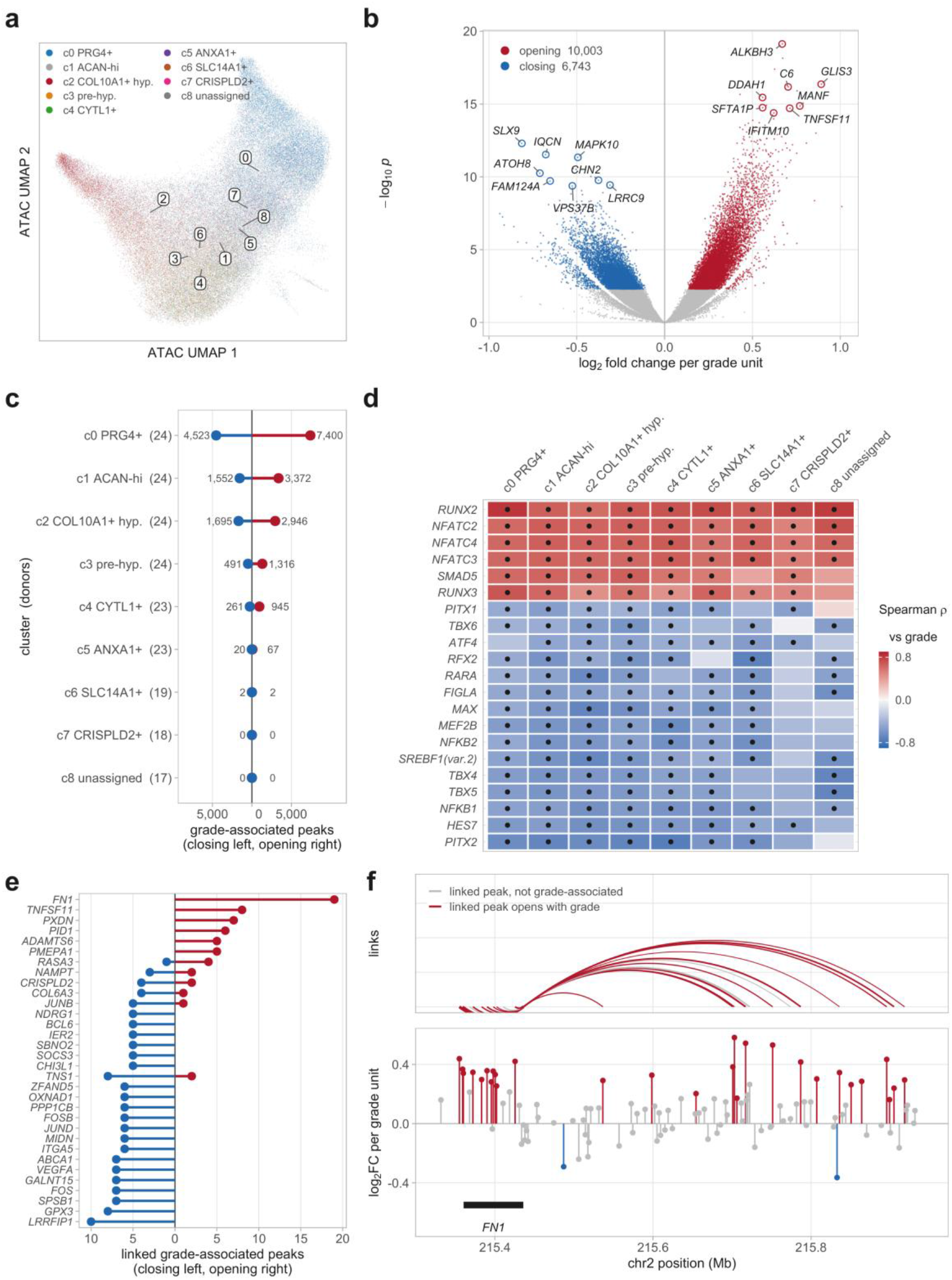
Chromatin accessibility changes with macroscopic knee grade in the same direction across chondrocyte clusters. Single-nucleus multiome of 81,670 chondrocyte nuclei from 24 of the 30 donors, labeled with the clusters of Figure 3 by barcode. **(a)** UMAP of the integrated ATAC embedding, colored by cluster, with cluster numbers as in Figure 3a. **(b)** Grade-associated accessibility pooled over clusters, from DESeq2 on per-donor pseudobulk against continuous grade; colored points are regions opening and closing at adjusted p value < 0.05, and labels are the nearest gene of the most significant regions. **(c)** Grade-associated regions tested within each cluster, closing left and opening right, with the number of contributing donors in brackets; clusters with fewer donors and nuclei have fewer testable regions. **(d)** Transcription factor motif activity against grade within each cluster, as the Spearman correlation of donor-mean chromVAR deviation with grade, for the 21 motifs associated with grade at nominal p < 0.05 in at least seven of the nine clusters; dots mark nominal p < 0.05. Of 5,697 cluster-motif tests, 625 reached nominal p < 0.05 and three remained significant after correction across 633 motifs within each cluster (*RUNX2* in c0 and c5, *PITX2* in c4). **(e)** Protein-coding genes with five or more linked grade-associated regions, from peak-gene links crossed with panel b, closing left and opening right. **(f)** The *FN1* locus: linked regions drawn as arcs from the transcription start site, width proportional to link z-score (top), and the grade effect of every tested region in the window (bottom). Peak-gene links are correlations between accessibility and expression across nuclei, not demonstrated regulation.

With grade, 16,746 of 175,071 accessible regions changed (10,003 opening, 6,743 closing; Figure 9b), robustly to leaving out any single donor. Opening regions were enriched in introns and distal intergenic regions and closing regions at promoters (Figure S22b). Within subsets, 15,474 regions changed in at least one, and none of the 5,204 significant in more than one subset changed in opposite directions (Figure 9c, Figure S22c-d).

*RUNX2* motif accessibility rose with grade in all nine subsets (nominal p < 0.05; ρ = 0.60 to 0.82; the most positively grade-correlated motif in seven), as did the *NFATC2*, *NFATC3* and *NFATC4* motifs (all nine subsets), *RUNX3* (eight) and *SMAD5* (seven), whereas *PITX2*, *HES7*, NF-κB and *MEF2B* motifs fell in seven or eight subsets (Figure 9d). Only *RUNX2* in c0 and c5 and *PITX2* in c4 remained significant after correction across 633 motifs, so the other motif changes are nominal; the NFAT and SMAD motif changes were concentrated in grade 4 knees, six of seven of which were obtained at arthroplasty.

Of 5,712 region-gene links, 922 ran through a grade-associated region, reaching 479 genes, and these were strongly one-directional: of genes with two or more such regions, 50 had all opening, 129 all closing and 21 were mixed (Figure 9e). *FN1* had the most, with 19 of its 23 linked regions opening and none closing (Figure 9f, Figure S23a), a pattern repeated in c0 to c4 (Figure S22e); *TNFSF11* had eight, all opening (Figure S23b). Conversely, all six grade-associated regions linked to *ITGA5* closed (Figure S24a-c), consistent with the lower *ITGA5*/*ITGB1* share of inferred *FN1* signaling (Figure 7d), as did those linked to *ABCA1* and *GPX3* (Figure S24d-i), which fell with grade in the snRNA-seq and bulk cohorts, respectively (Figure S10b, Figure 1b); their accessibility fell with grade across donors (ρ = −0.57 to −0.67) (Figure S24c, f, i). By contrast, *FGF1* promoter accessibility did not change although *FGF1* RNA rose in the same nuclei, and the *FGFR1* promoter closed modestly without a change in *FGFR1* RNA (Figure S18d).

### *FN1* is induced in excess of its assembly machinery, together with *HTRA1*

*FN1* recurred across these analyses: it was induced in every subset, changed its splicing, shifted its inferred receptor use and had the most linked regions opening with grade. We therefore asked whether the fibronectin assembly machinery rose with *FN1*. *FN1* rose with grade in all nine clusters (log2 fold change 0.78 to 1.20 per grade unit), but the machinery that assembles fibronectin into fibrils did not: *ITGA5*, *ITGB1*, *ITGAV*, *TLN1*, *FERMT2* and *SDC4* were unchanged, *ITGB3* and *TNS1* decreased in single clusters and *ILK* rose modestly in c0 only (Figure 10a). *ITGA5* did not rise in any cluster or in bulk, although increased *ITGA5* and *ITGB1* expression and α5β1 formation have been reported in cultured OA chondrocytes (36), and ERp57 (*PDIA3*), which promotes fibronectin fibril formation in cartilage (37), rose only modestly in bulk. Of the fibronectin-cleaving proteases, *HTRA1* rose in every cluster (Figure 10a), whereas *MMP2*, *MMP3* and *MMP14* did not and *MMP13* was not detected. *HTRA1* tracked *FN1* from donor to donor in all nine clusters independently of grade (partial ρ = 0.51 to 0.76; Figure 10b-c). *HTRA1*, like *FN1*, is induced by TGF-β through ALK5/SMAD2/3 signaling in chondrocytes (38,39), and adjusting for a SMAD3 early-response score weakened their coupling (median partial ρ from 0.63 to 0.50) but left it significant in eight of nine clusters, so a shared TGF-β response may account for part but not all of their co-induction.

**Figure 10.**
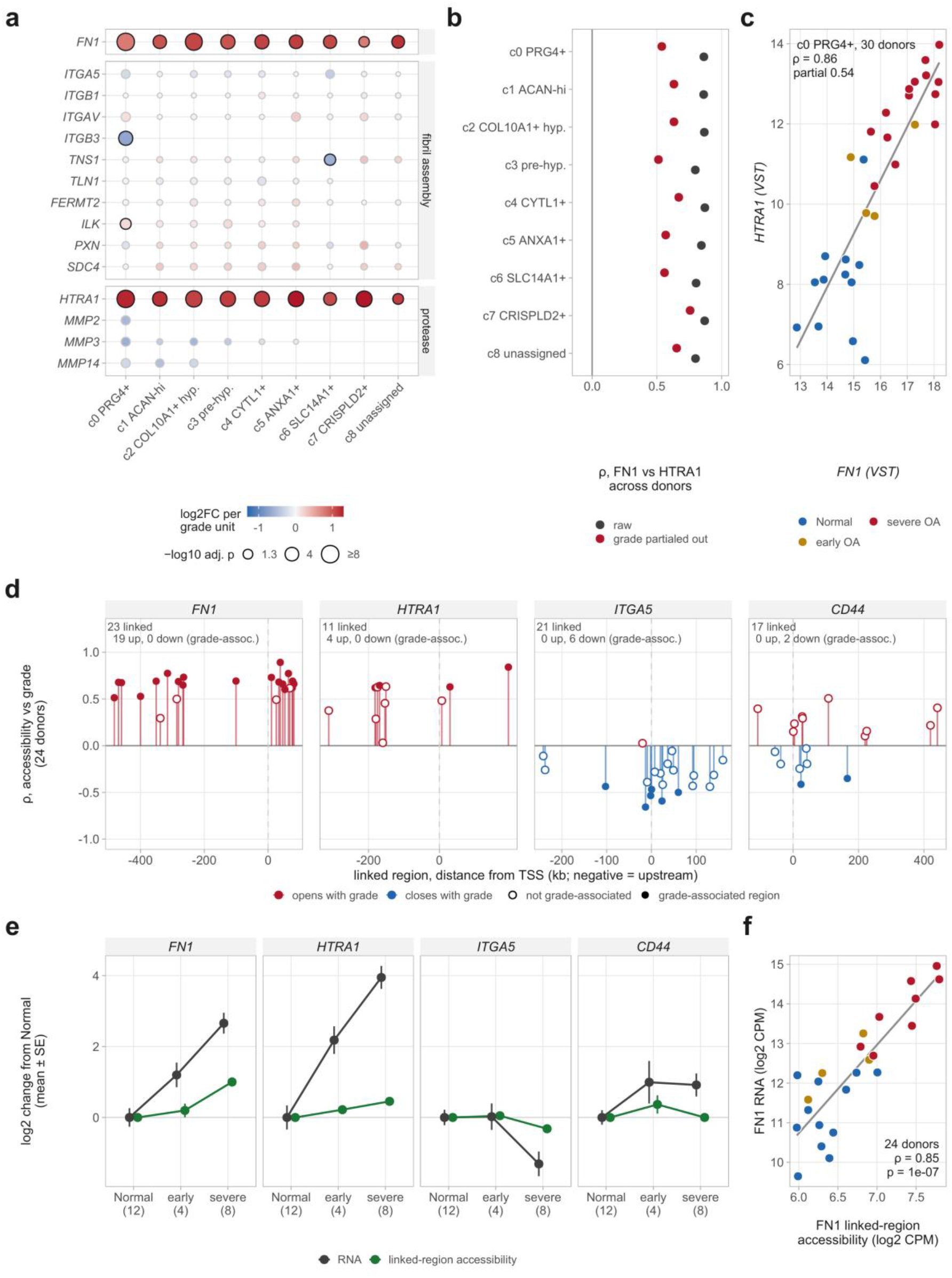
OA chondrocytes produce fibronectin in excess of its assembly machinery, together with *HTRA1*, and both are reflected in chromatin. **(a)** Grade association of *FN1*, the fibronectin fibril-assembly machinery and fibronectin-cleaving proteases in each cluster (continuous-grade pseudobulk model of Figure S10; 30 donors); color is log2 fold change per grade unit (capped at ±1.3), size −log10 adjusted p, and a black outline marks adjusted p < 0.05; a blank position means the gene was below the expression filter in that cluster. **(b)** Spearman correlation between *FN1* and *HTRA1* across donors within each cluster, before (gray) and after (red) partialing out grade; all are adjusted p < 0.05 (Benjamini-Hochberg across clusters). **(c)** *HTRA1* against *FN1* in each donor’s c0 pseudobulk (variance-stabilized expression), colored by stage; the line is a linear fit for display. **(d)** Every region linked to *FN1*, *HTRA1*, *ITGA5* and *CD44* by peak-gene linkage in the single-nucleus multiome, plotted by distance from the gene’s transcription start site (negative, upstream): Spearman correlation of each region’s accessibility (counts per million, per donor pseudobulk) with grade across 24 donors; filled points are regions in the grade-associated set of Figure 9b (adjusted p < 0.05). Counts give the linked regions and the grade-associated regions opening (up) or closing (down). **(e)** Accessibility summed over each gene’s linked regions, and RNA, in the same nuclei and per donor pseudobulk, as log2 change from the Normal mean; mean ± standard error, with donors per stage in brackets. (f) *FN1* linked-region accessibility against *FN1* RNA in each donor (same nuclei), colored by stage; Spearman correlation across 24 donors.

In the multiome data, all 23 *FN1*-linked and all 11 *HTRA1*-linked regions gained accessibility with grade (19 and four significantly), 20 of the 21 *ITGA5*-linked regions lost it, and *CD44*-linked regions changed in both directions (Figure 10d). Summed accessibility rose with RNA for *FN1* and *HTRA1*, fell with it for *ITGA5* at severe OA, and did not change for *CD44*, whose RNA rose in the multiome nuclei (Figure 10e) but not in the 30-donor snRNA-seq grade model. *FN1* linked-region accessibility tracked *FN1* RNA across donors (ρ = 0.85) and rose together with it rather than ahead of it (Figure 10e-f).

OA chondrocytes thus induce *FN1* in excess of the machinery that assembles it, together with a protease that cleaves it, and the fibronectin they produce also differs in isoform (Figure 2).

### TGF-β/SMAD response genes rise without more of the dominant ligand, and this signaling is a candidate regulator of *FN1*

Because TGF-β induces fibronectin and *HTRA1* in chondrocytes through ALK5/SMAD2/3 signaling (38,39), and integrin αvβ8, which gained a share of inferred fibronectin signaling in OA, activates latent TGF-β (40), we examined whether TGF-β signaling could account for the induction of *FN1* (Figure 11). TGF-β-responsive genes rose with grade in most clusters (*SKIL* and *LTBP2* in all nine, *COMP* in eight; Figure 11a), and the Hallmark TGF-β signaling gene set was positively enriched among grade-associated genes in all nine clusters, significantly in six (Figure S10d). A SMAD3 early-response score, excluding *FN1*, *COMP* and *LTBP2*, rose with grade in c0 and c2 to c5 (0.21 to 0.30 z per grade unit), whereas a BMP target score did not change (Figure 11b). The dominant ligand did not increase: *TGFB2*, which carried most inferred TGF-β signaling, was lower with grade in all nine clusters (significantly in two; *TGFB1* rose only in c2; Figure 11a), inferred TGF-β communication was lower at severe OA than in Normal cartilage, and *ITGB8*, which encodes the αvβ8 β subunit, rose only in c0. SMAD2/3 motif accessibility, scored over a family of eight SMAD2/3 motifs, rose with grade in seven subsets (ρ = 0.46 to 0.58; adjusted p = 0.015 to 0.036 across subsets; Figure 11c). The single JASPAR SMAD3 motif did not survive correction across 633 motifs, SMAD motifs do not distinguish SMAD2/3 from the BMP-responsive SMAD1/5, whose motif family rose similarly, and the SMAD motif rise was concentrated in grade 4 knees. Grade-associated regions linked to *TGFBR1* and the SMAD target *PMEPA1* opened, whereas the single region linked to *TGFB2* closed (Figure 11d). TGF-β/SMAD response genes and receptor-linked chromatin thus increased without an increase in the dominant ligand, consistent with increased activation of latent, matrix-bound TGF-β.

**Figure 11.**
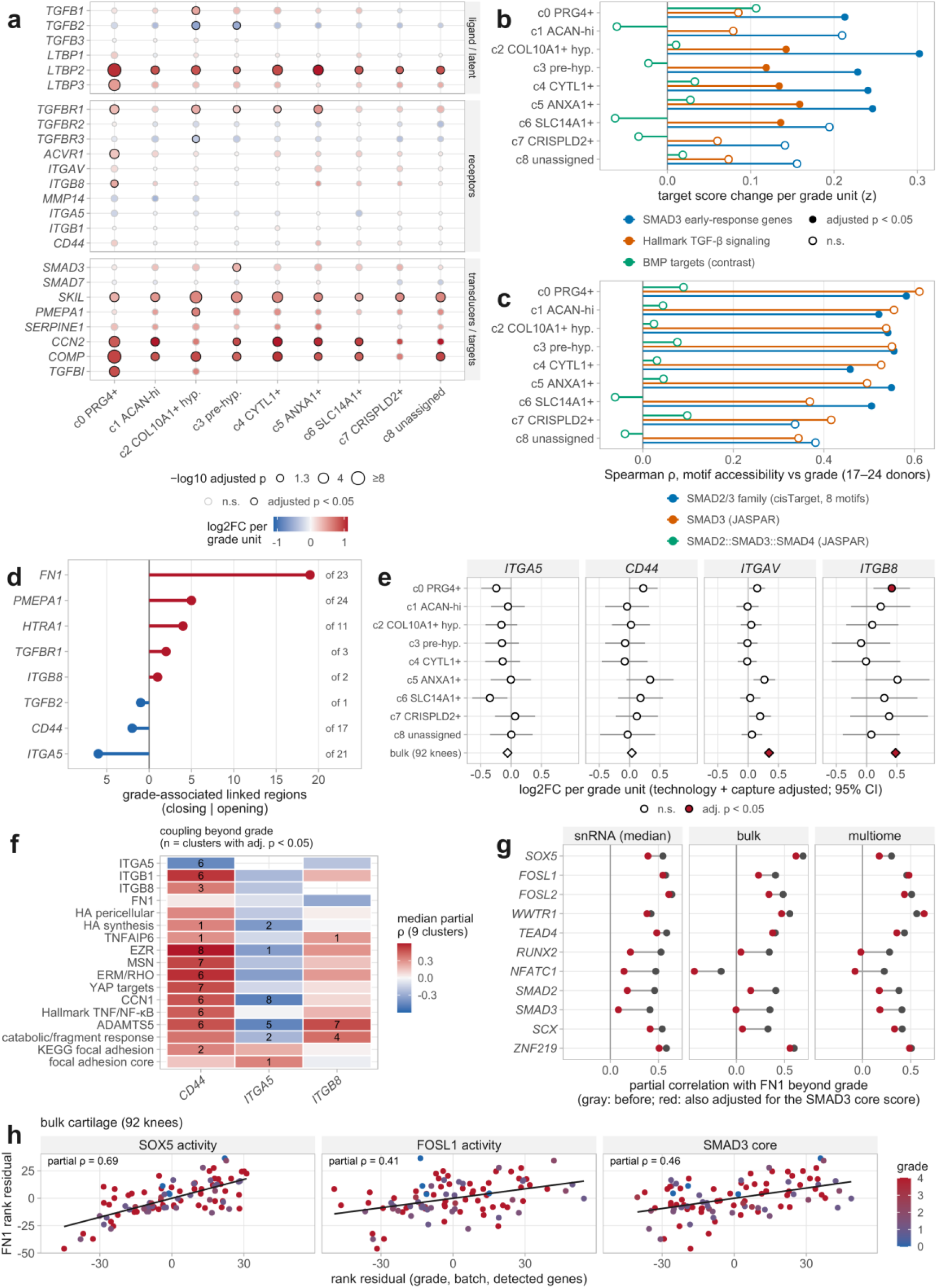
TGF-β/SMAD response genes rise without more of the dominant ligand, and this signaling is a candidate regulator of *FN1*. **(a)** Log2 fold change per grade unit (continuous-grade model of Figure S10) of TGF-β ligands and latent-complex genes, receptors and activators, and signal transducers and target genes in each cluster; size shows −log10 adjusted p and a black outline marks adjusted p < 0.05. A blank position means the gene was below the expression filter in that cluster. **(b)** Donor-level target scores (mean z-scored variance-stabilized expression of the set’s genes in each donor’s cluster pseudobulk) regressed on grade with technology and capture burden as covariates, as change in score per grade unit: a SMAD3 early-response set of 17 genes (*SERPINE1*, *SMAD7*, *SKIL*, *PMEPA1*, *TGFBI*, *CCN2*, *JUNB*, *BHLHE40*, *TGIF1*, *KLF10*, *SNAI2*, *SKI*, *LTBP1*, *COL1A1*, *TIMP3*, *CDKN2B* and *BAMBI*; *FN1*, *COMP* and *LTBP2* excluded to avoid circularity; 11 to 17 of them passed the expression filter per cluster), the Hallmark TGF-β signaling set, and a BMP target set as a specificity contrast. Filled points are adjusted p < 0.05 (Benjamini-Hochberg across clusters within each set). **(c)** Accessibility of SMAD motifs against grade within each cluster in the multiome cohort, as the Spearman correlation between each donor’s mean chromVAR deviation and grade (17 to 24 donors per cluster): the cisTarget SMAD2/3 motif family (eight motifs), and the JASPAR SMAD3 (MA0795.1) and SMAD2::SMAD3::SMAD4 (MA0513.1) motifs. Filled points are adjusted p < 0.05 (Benjamini-Hochberg across clusters for the cisTarget family and across the 633 JASPAR motifs within each cluster for the JASPAR motifs). **(d)** Grade-associated accessible regions (Figure 9b) linked to TGF-β pathway genes by peak-gene linkage (Figure 9e), as the number that open (right) or close (left) with grade; the number beside each gene is its total number of linked regions. *FN1*, *HTRA1*, *ITGA5* and *CD44* are included for reference (Figure 9f, Figure S24); *JUNB* (1 opening and 5 closing of 27 linked regions) is not shown. **(e)** Change per grade unit in *ITGA5*, *CD44*, *ITGAV* and *ITGB8* in each cluster (pseudobulk model with technology and capture burden as covariates, 19 to 30 donors per cluster) and in the bulk cohort (92 knees, differential expression covariates); 95% confidence intervals for clusters only, filled symbols adjusted p < 0.05. **(f)** Coupling across donors with *CD44*, *ITGA5* and *ITGB8* beyond grade, capture burden and technology: median partial Spearman correlation over nine clusters (single genes kept in every cluster regardless of the expression filter; program scores (rows) are mean z of the focal adhesion core, ERM/RHO, catabolic/fragment response, YAP targets, Hallmark TNF/NF-κB, KEGG focal adhesion, hyaluronan (HA) synthesis and HA pericellular gene sets defined in Supplementary Methods); numbers are clusters with adjusted p < 0.05. **(g)** Partial correlation with *FN1* beyond grade (and technology in snRNA-seq, or batch and detected genes in bulk; capture burden not included) before (gray) and after (red) additional adjustment for the SMAD3 core score, in each dataset. **(h)** Bulk cartilage (92 knees): *FN1* against *SOX5* activity, *FOSL1* activity and the SMAD3 core score, as rank residuals on grade, batch and detected genes; colored by grade.

The shift in *FN1* receptor share (Figure 7d-e) occurred without a change in the receptor genes: *ITGA5* and *CD44* did not change with continuous grade in any subset or in bulk (*CD44* differed between Normal and severe OA only in c0 in the stage model; Figure 8b), whereas *ITGB8* rose in c0 and in bulk, and *ITGAV* and, slightly, *ITGB1* in bulk (Figure 11e). Instead, *CD44* and *ITGA5* varied reciprocally from donor to donor independently of grade (median partial ρ = −0.45, significant in six of nine subsets; Figure S25b), a reciprocity not detected in bulk, and high-*CD44* donors also expressed more *EZR*, *MSN* and YAP target genes (Figure 11f). Of the hyaluronan-axis genes that act with *CD44*, *CEMIP* rose with grade in seven of nine subsets and *TNFAIP6* in c0 and in bulk (Figure S25a).

Combining prior regulatory evidence, motifs in the 19 *FN1*-linked regions that opened with grade, inferred transcription factor activity and its coupling with *FN1* beyond grade in the snRNA-seq, bulk and multiome data (Figure S26a) pointed to three candidate regulators of *FN1*. A SMAD3 early-response score tracked *FN1*-linked region accessibility beyond grade (partial ρ = 0.53, p = 0.008) and accounted for the coupling of *RUNX2* and *NFATC1* activity with *FN1* and partly for that of *SCX*. The coupling of *FOSL1* and *FOSL2* with *FN1* persisted after adjustment for the SMAD3 score in all three datasets. The *SOX5* motif occurred in 10 of the 19 opening regions (nominal p = 4×10⁻⁴, not significant after correction), and *SOX5* activity coupled most strongly with *FN1* in bulk (Figure 11g-h, Figure S26a). Coupling was not specific, however, as factors without known links to *FN1* coupled as strongly, and footprinting showed no increase in footprint depth with grade at *SOX5*, *FOSL1*, *FOSL2*, SMAD, TEAD or *FOXD1* motifs, whereas accessibility around *RUNX2* motifs (in c0) and around SMAD3 motifs (mainly in arthroplasty knees) rose with stage (Figure S26b-c). *FN1* induction is thus associated with TGF-β/SMAD, *FOSL1*/2 and *SOX5* activity, but its direct regulators remain to be established.

## Discussion

Profiling cartilage across the full spectrum of macroscopic OA grade, rather than comparing normal with end-stage tissue, suggests a different view of how chondrocytes respond to disease. OA progression appears to be carried less by the emergence or loss of particular chondrocyte populations than by a change of state in which every population takes part, with differing emphasis. This change appears to include a remodeling of how chondrocytes receive signals from their matrix, most clearly in inferred fibronectin signaling.

Earlier single-cell studies of OA cartilage proposed that disease is driven by the expansion of specific pathogenic populations (6–9). Our data point to a different emphasis. The chondrocyte clusters persisted across all disease stages, and the dominant disease signal was a common response shared across clusters, without opposing changes, in both the transcriptome and chromatin. Our analysis of cluster identity supports the view that the clusters are positions along a continuum of chondrocyte states that the disease shifts as a whole, rather than discrete lineages that replace one another. In healthy cartilage the clusters were ordered along a superficial-to-deep and a hypertrophy axis without gaps between them, and only the *PRG4*+ superficial and *COL10A1*+ hypertrophic chondrocytes at either end were distinct by expression and gene programs, and the hypertrophic cells also by chromatin, with a pre-hypertrophic state between the hypertrophic cells and the rest. The other clusters were reproducible states defined by single genes or by combinations of shared programs, which may explain why earlier studies have described differing numbers and kinds of chondrocyte populations (6–9). Where the clusters did differ in OA, they differed in the strength of components of the shared response, matrix production being strongest in the *PRG4*+ superficial cells, the secretory pathway in the hypertrophic and *CYTL1*+ cells, and inflammatory signaling in the *ANXA1*+ and *CRISPLD2*+ cells; none followed a distinct trajectory, although superficial chondrocytes reached part of the ECM response, including *FN1*, earlier (Figure 8). This does not exclude rare or transitional populations below the resolution of our clustering, and the shifts in abundance we did detect may matter locally. It does, however, suggest that interventions acting on the shared response would reach chondrocytes throughout the tissue rather than a minority population.

The shared response combines increased expression of matrix synthesis and secretion genes with loss of programs that are protective in cartilage. The rise in secretory and ER stress pathways, together with induction in bulk cartilage of *CREB3L2*, an ER stress transducer required for secretion of cartilage matrix (41), suggests that part of the stress signature of OA reflects increased secretory load rather than degeneration alone. At the same time, programs with protective roles in cartilage decline, including glucocorticoid-responsive and autophagy-related genes such as *DDIT4* (42), FOXO-dependent stress resistance (5,43) and cholesterol efflux, which is impaired in OA chondrocytes (44). These changes are consistent with a repair response that chondrocytes may not be able to sustain, in which matrix production increases while protective programs decline. The finding that the subsets make matrices of different composition at a similar overall level, and that OA shifts all of them toward fibronectin and small leucine-rich proteoglycans at the expense of hyalectans, indicates that OA changes the kind of matrix made as well as the amount. Increased synthesis may fail to restore cartilage, in part because the matrix produced differs in composition, which may alter its assembly and mechanical properties.

The fibronectin findings from several lines of evidence suggest that OA may affect the assembly, processing and reception of fibronectin as well as the amount made. Chondrocytes induced *FN1* in excess of the machinery that assembles fibronectin into fibrils, while α5β1, through which intact fibronectin transmits protective signals in cartilage (18,45), did not rise with it and lost part of its share of inferred fibronectin signaling. Fibronectin that is not incorporated into fibrils may be more accessible to proteolysis, and *HTRA1*, which generates fibronectin fragments that induce matrix metalloproteinases in synovial fibroblasts (46), was induced together with *FN1* donor by donor, possibly in part through the TGF-β response that induces both (38,39). Fibronectin fragments, which OA chondrocytes can generate through proteases such as ADAM-8 (47), accumulate in OA cartilage and synovial fluid and induce matrix degradation (15–17,48), and the receptors that gained a share of inferred fibronectin signaling can mediate fragment or TGF-β effects: *CD44* binds the heparin-binding domain of fibronectin (49) and contributes to MMP induction by a heparin-binding fibronectin fragment in human cartilage (50), and αvβ8 activates latent TGF-β (40). Apart from rises in *ITGB8* and, in bulk, *ITGAV* and slightly *ITGB1*, the receptor genes did not change with grade; rather, the rise in *FN1* met a receptor repertoire that differed between individuals along a reciprocal *CD44*-*ITGA5* axis independent of grade, which may contribute to differences between patients in how fibronectin and its fragments are received. In our data, TGF-β/SMAD response genes rose with grade, receptor-linked regions opened and SMAD motif accessibility rose, the last mainly in grade 4 knees, even though the dominant ligand tended to decline and the latent TGF-β-binding protein *LTBP2* was induced in every cluster, consistent with increased activation of matrix-bound latent TGF-β rather than increased ligand supply. Fibronectin signaling between chondrocytes has been noted in earlier single-cell studies of human cartilage (8,51), and fibronectin induces SMAD2/3 signaling in human chondrocytes (51). The fibronectin made in OA cartilage also changed in kind, toward the EDB-containing isoform, which is associated with chondrogenic differentiation (52) and accumulates as protein in OA cartilage (25), without the rise in the EDA isoform that marks fibrotic matrix. EDB inclusion was the most significant splicing change with grade in the cartilage transcriptome, and the few other matrix transcripts whose splicing changed, among them *DCN*, *ABI3BP* and, among matrisome events, *PRG4*, together with a shift away from the α4β1-binding CS1 segment of fibronectin, suggest that the matrix produced in OA differs in isoform as well as in amount. The regulators of *FN1* induction remain uncertain: TGF-β/SMAD, *FOSL1*/2 and *SOX5* activity tracked *FN1* across donors, but footprinting provided no direct evidence of increased binding. A truncated fibronectin variant that lacks the α5β1-binding site, increased in lesioned relative to preserved OA cartilage and impairing matrix deposition (26), was rare and did not change with grade in our cohort, suggesting that it marks focal lesions and is not part of the grade-wide program. Together these observations support a model (Figure 12) in which OA chondrocytes mount a developmental, repair-type fibronectin program but may not assemble all of its product, so that more fibronectin would be processed by *HTRA1* and received, intact or as fragments, through *CD44* and αvβ8, which are coupled to catabolic and TGF-β signaling, rather than predominantly through α5β1, although fragments can also signal through α5β1 itself (53,54). That much of the inferred receptor shift was already present in the four early-OA donors (nominally, and about half in the expression score), when *FN1* had risen in superficial chondrocytes but little in the other subsets, suggests that it is part of the chondrocyte response rather than solely a late consequence of matrix breakdown, although *HTRA1* was also already induced in superficial chondrocytes at this stage; the closing of chromatin at *ITGA5* is evident mainly at severe OA. Because these conclusions rest on mRNA, which cannot distinguish intact fibronectin from its fragments, the model requires direct experimental testing, such as domain-specific detection of fibronectin fragments and of EDB-fibronectin together with *HTRA1* in cartilage, and receptor blockade in human chondrocytes.

**Figure 12.**
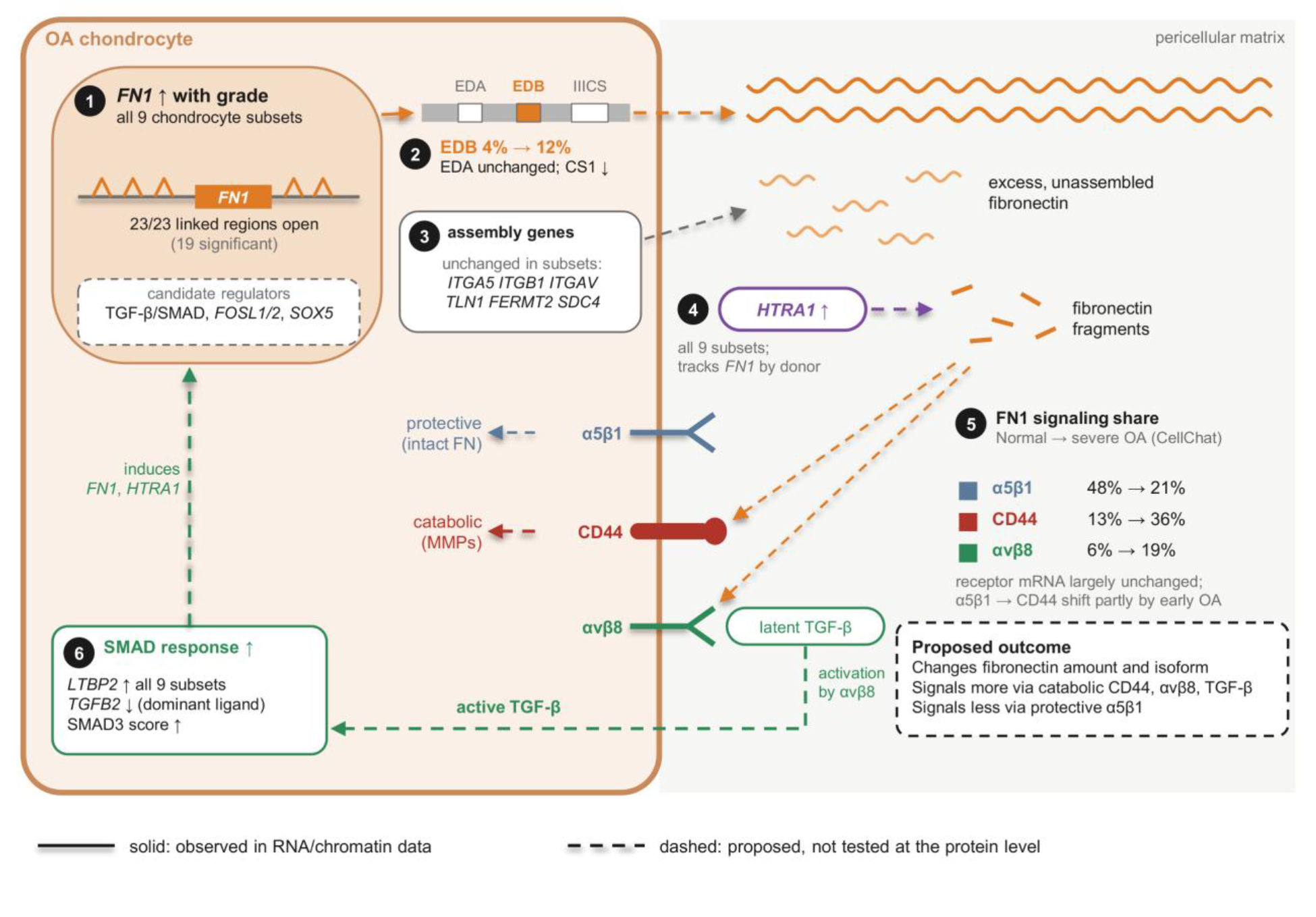
Proposed model of fibronectin production, processing and reception in OA chondrocytes. (1) *FN1* expression increased with grade in all nine chondrocyte subsets, and all 23 *FN1*-linked chromatin regions gained accessibility (Figure 10). (2) Inclusion of the EDB exon increased, EDA was unchanged, and IIICS splicing shifted away from the CS1 segment (Figure 2). (3) Expression of the core fibril-assembly genes (*ITGA5*, *ITGB1*, *ITGAV*, *TLN1*, *FERMT2*, *SDC4*) did not increase in chondrocyte subsets, suggesting that fibronectin is produced in excess of its assembly machinery (Figure 10). (4) *HTRA1* increased in all subsets and tracked *FN1* across donors, suggesting increased fibronectin fragmentation (Figure 10). (5) The inferred share of *FN1* signaling moved from α5β1 toward *CD44* and αvβ8, the shift toward *CD44* partly present at early OA (four donors, nominal; Figure 7, Figure 8d, Figure S21). (6) TGF-β/SMAD response genes increased without more of the dominant ligand, consistent with activation of latent TGF-β, which may in turn induce *FN1* and *HTRA1* (Figure 11). Percentages compare Normal with severe OA (donor-level CellChat), and EDB inclusion grade 0-1 with grade 3-4 knees. Solid lines, observed in RNA or chromatin data; dashed lines, proposed and not tested at the protein level.

Although no gene departed from a linear change with grade in bulk RNA-seq, the single-nucleus stage analysis separated changes already present at early OA, in tenascin-C, *JAG1* and, partly, *FN1* receptor use, from those added with progression, including *FGF1* and developmental and ossification programs (Figure 8). Inferred TENASCIN signaling, through tenascin-C, an endogenous *TLR4* activator (55), rose by early OA and remained elevated, whereas later stages showed a shift in FGF ligands from *FGF2*, which is chondroprotective in murine OA (11), toward *FGF1*, which is catabolic in chondrocytic cells (56). Whether this shift translates into signaling is less certain. *FGF1* leaves cells through a stress-induced route whose components barely rose with it, so in OA cartilage it may be released mainly when chondrocytes are injured, and expression of the SULF endosulfatases, which remove the 6-O sulfates that FGF-FGFR complexes require, rose with grade (*SULF1*) or tracked *FGF1* across donors (*SULF2*). *SULF1* and *SULF2* are overexpressed in OA cartilage, where they suppress FGF signaling while enhancing BMP signaling in chondrocytes, and they protect cartilage in mice (57). *SULF1* was higher with grade in all nine chondrocyte subsets, significantly in *PRG4*+ chondrocytes, in keeping with this earlier finding, and this induction may limit the consequences of the FGF ligand shift, consistent with the absence of an increase in ERK target expression. The balance of FGF ligands and receptors, rather than total FGF signaling, is thought to determine their effect on cartilage (58,59), and changes in heparan sulfate-modifying genes suggest that the matrix may further modulate this balance. In chromatin, increasing *RUNX2* motif accessibility across clusters (significant after correction in c0 and c5) is in keeping with its role in promoting OA in mice (60,61), and suggests that part of the hypertrophic program that distinguishes c2 and c3 in healthy cartilage may be engaged more widely in OA, while a rise in *RUNX3* motif accessibility, evident only at nominal significance, would be in keeping with a compensatory response, given that *RUNX3* protects cartilage and regulates lubricin and aggrecan (62).

Several limitations qualify these interpretations. The study is cross-sectional, so the sequence of events is inferred from between-donor differences rather than longitudinal observations. Within the bulk cohort, however, the grade response was consistent with a gradual, linear change, and it was reproduced in two public cartilage datasets, although one of them comes from an earlier cohort of our group (Figure S2). Intermediate grades were sparsely represented in the single-nucleus cohort. Because nearly all grade 4 cartilage was obtained at arthroplasty, and in the single-nucleus cohort was also sequenced with a different chemistry, tissue source is itself a marker of disease severity in this cohort, and for features confined to grade 4 the effects of end-stage disease and of tissue source cannot be separated. Age and grade are correlated in this cohort, as in most human OA cohorts, and their effects were not separated here. Cluster identity was defined in healthy donors profiled with a single chemistry, and part of the boundary between the hypertrophic and pre-hypertrophic subsets rested on long genes, which are detected preferentially in nuclei. Communication was inferred from mRNA; the stage networks pooled nuclei, and although the main receptor shifts were reproduced with the donor as the unit (Figure S21), the early-OA estimates rest on four donors.

In conclusion, OA severity in human knee cartilage is associated with a broadly shared change in chondrocyte state in which all chondrocytes take part, with differing emphasis along a zonal and hypertrophic continuum: matrix production rises while protective programs decline, the fibronectin produced shifts in isoform, and inferred fibronectin reception appears to be partly remodeled already at early OA, although this rests on four donors. These findings suggest that the assembly, processing and reception of fibronectin, and not only the amount made, merit investigation as targets in OA.

## Methods

### Human cartilage samples

Human tissue collection was approved by the Scripps Human Subjects Committee (protocol 20-7636). Informed consent/authorization was obtained for research use of donor tissue. Knees were obtained from organ donors without a history of knee osteoarthritis, rheumatoid arthritis or knee injury, through a tissue procurement program. Arthroplasty cartilage was obtained from patients undergoing total knee arthroplasty (TKA) at Scripps Clinic (La Jolla, CA). The entire knee cartilage surfaces from all compartments were scored macroscopically with the Outerbridge method (21) (macroscopic knee grade, 0 to 4). Osteochondral specimens were resected from the weight-bearing area of the medial femoral condyle and full-thickness cartilage was separated from the subchondral bone. Cartilage histopathology was assessed using the OARSI system and is referred to here as the OARSI score (24). For analyses by disease stage, donors were grouped as Normal (macroscopic knee grade 0-1), early OA (grade 1.5-2) and severe OA (grade 3-4).

Single-nucleus cohort. Thirty samples from 30 donors (one sample per donor; 22 right and 8 left knees) were profiled: 11 Normal, 4 Aged, 3 OA and 12 total knee arthroplasty (TKA) donors, aged 18-86 years (median 61), 19 male and 11 female, body mass index 19.5-47.6 kg/m², post-mortem interval 13.5-53.5 h for cadaveric donors (1.25-6 h from surgery to processing for arthroplasty samples). Macroscopic knee grades were 0 (n = 3), 1 (n = 9), 1.5 (n = 3), 2 (n = 1), 3 (n = 1) and 4 (n = 13), giving 12 donors at the Normal stage (grade 0-1), 4 at early OA and 14 at severe OA. Twenty-four samples were profiled with the single-nucleus multiome assay and six, all grade 4 arthroplasty samples, with the GEM-X single-nucleus assay; chemistry (the technology term) was therefore included as a covariate in every single-nucleus regression model that included GEM-X donors and, where relevant, analyses were repeated in the 24 multiome donors only.

Bulk cohort. Ninety-two cartilage samples from 92 knees (one per donor) were analyzed: 17 Normal, 16 Aged, 12 OA and 47 TKA, with macroscopic knee grades 0 (n = 4), 1 (n = 19), 1.5 (n = 10), 2 (n = 5), 3 (n = 3) and 4 (n = 51), aged 18-91 years, 56 male and 36 female, sequenced in two batches (81 and 11 samples). Of these, 45 were from cadaveric donors and 47 from total knee arthroplasty.

Knee identity, side, macroscopic grade and OARSI score of each knee used for bulk RNA sequencing were cross-checked against the donor tissue inventory. One knee (OA donor 7) had no score, leaving 91 knees (Normal 17, Aged 16, OA 11, arthroplasty 47). Agreement with macroscopic grade is reported as the Spearman correlation.

### Nuclei isolation, library preparation and sequencing

Nuclei were isolated from cartilage stored in liquid nitrogen using the Singulator S200 Automated Tissue Dissociation System (S2 Genomics), following a previously published protocol (protocols.io, https://doi.org/10.17504/protocols.io.ewov193molr2/v1). Briefly, 100-150 mg of frozen tissue was minced and processed in a Singulator cartridge containing RNase inhibitor with the low-volume nuclei isolation program, and the suspension was filtered (20 µm and 40 µm strainers) and centrifuged twice at 150 × g for 10 min at 4 °C. Nuclei were counted after acridine orange/propidium iodide (AO/PI) staining on a Countess II FL Automated Cell Counter, and nuclear integrity was inspected by microscopy. Nuclei were diluted to 4,000-5,000 nuclei/µL for a final recovery of 20,000 nuclei in 5 µL. Libraries were prepared with the Chromium Next GEM Single Cell Multiome ATAC + Gene Expression kit (10x Genomics, PN-1000283), with 33,000 nuclei loaded per reaction, and sequenced paired-end on an AVITI system (Element Biosciences). The six GEM-X libraries were prepared with the Chromium GEM-X Single Cell 3′ v4 kit (10x Genomics, PN-1000691), with 15,000 nuclei loaded per reaction (targeting 10,000 nuclei), and sequenced on an AVITI system.

Multiome libraries (24 donors) were aligned and counted with Cell Ranger ARC 2.0.2 against the refdata-cellranger-arc-GRCh38-2020-A-2.0.0 reference, and single-nucleus RNA-seq libraries from the remaining six arthroplasty donors (Chromium GEM-X Single Cell 3′ v4) with Cell Ranger 9.0.1 against refdata-gex-GRCh38-2024-A; intronic reads were included in both. Multiome and GEM-X gene-expression matrices were merged by gene symbol. Because the two Cell Ranger references (GRCh38-2020-A, GENCODE v32; GRCh38-2024-A, GENCODE v44) assign different symbols to 12,913 shared Ensembl genes (mostly unnamed long non-coding RNAs; 468 protein-coding), these genes were counted under different symbols in the two chemistries; no gene named in this study is affected. Integration features were restricted to symbols present in every sample.

### Bulk RNA sequencing and quantification

Total RNA was extracted from 100 mg of homogenized frozen cartilage with TRIzol reagent (Thermo Fisher Scientific), followed by on-column DNase treatment and cleanup (RNeasy MinElute Cleanup Kit, QIAGEN); integrity was assessed on an Agilent 4200 TapeStation. Libraries were prepared with the QIAseq Stranded mRNA Select Low Input Kit (QIAGEN, cat. 334205) with ribosomal RNA depletion (cat. 334235) and sequenced on an Illumina NovaSeq X (2 × 151-bp paired-end reads; target depth approximately 30 million read pairs per sample). Library strandedness was reverse (dUTP-type), as confirmed with RSeQC infer_experiment. Libraries were sequenced in two batches, and batch was included as a covariate in the bulk models.

Reads were trimmed with Trim Galore (Cutadapt) in paired mode with explicit adapter sequences, removing the first 16 bases of read 2, and aligned to GRCh38 with HISAT2 (prebuilt grch38 index, --dta). Gene-level counts were obtained with featureCounts against the GENCODE v44 comprehensive annotation (exon features summarized by gene_id; reverse-stranded, paired-end, both mates aligned, chimeric pairs not counted). The number of detected genes was the number of the 62,700 annotated genes with at least one count. Libraries were retained if at least 10% of reads were assigned to genes and at least 5 million reads were assigned. No analyzed cartilage library failed these thresholds; seven further libraries were not analyzed: two failed sequencing quality control, four could not be assigned unambiguously to their knee, and one was a second library from a knee already included. Genes on chromosome Y or the mitochondrial chromosome and *XIST* were removed, and genes were retained if their TPM exceeded 1 in at least 25% of the cartilage libraries. This left 17,951 genes, which formed the gene universe for all bulk analyses.

RNA quality was measured for every library with RSeQC (read_distribution.py and infer_experiment.py) as the reverse-strand share of strand-informative reads and the percentage of reads in coding sequence. A library was flagged if its reverse-strand share was below 0.75 or its coding fraction below 10% (6 of the 92 libraries), and high-quality libraries were defined as reverse-strand share of at least 0.85 and coding fraction of at least 30% (75 knees). Flagged libraries were kept in the primary analyses. Principal components (Figure 1a) were computed on the 500 most variable variance-stabilized genes after removal of batch and detected genes with limma removeBatchEffect.

### Bulk differential expression with continuous grade

Differential expression was tested with DESeq2. Grade was entered as a continuous variable with the design ∼ Batch + Detected_Genes_z + Macro_Knee_Grade, where Detected_Genes_z is the z-scored number of detected genes (z-scored across the samples of the original multi-tissue run; because this differs from z-scoring within the 92 knees only by a linear rescaling, the grade estimates are identical). Size factors, dispersions and the negative binomial GLM were fitted with DESeq() defaults. The grade coefficient was tested with the Wald test, and p values were adjusted with the Benjamini-Hochberg method using results() defaults (independent filtering, alpha = 0.1, Cook’s distance outlier handling). Reported log2 fold changes per grade unit were shrunk with lfcShrink(type = “normal”); p values and adjusted p values come from the unshrunk Wald test, because shrinkage does not alter them. Genes were called differentially expressed at an adjusted p value < 0.05 (Figure 1b). For the gene lists used in motif enrichment, a mean normalized count (baseMean) of at least 10 was also required.

For gene set enrichment analysis, genes were ranked by −log10(p) × sign(log2 fold change) using the nominal Wald p value of the grade coefficient, keeping one Ensembl ID per symbol. clusterProfiler gseGO was run on GO Biological Process terms (gene set size 10-500, fgsea algorithm with eps = 0, set.seed(42), Benjamini-Hochberg adjustment), and terms with adjusted p < 0.05 were considered enriched. Terms shown in Figure 1c-d were selected to be non-redundant and to represent distinct programs (Supplementary Methods). Transcription factor motif enrichment was tested with RcisTarget separately on the genes rising and falling with grade (adjusted p < 0.05, baseMean ≥ 10), using the hg38 RefSeq r80 motif rankings (10 kb upstream and downstream of the transcription start site, mc9nr) and the version 9 HGNC motif annotation, with cisTarget defaults (area under the recovery curve over the top 5% of the ranking, NES threshold 3, direct and orthology-inferred annotations). Motifs with NES > 3 were collapsed to their high-confidence transcription factors, each summarized by its maximum NES. The 12 factors with the highest maximum NES per direction are shown (Figure 1e), excluding *MYLK*, which is not a transcription factor, so that 11 are shown for rising genes.

### Bulk alternative splicing

Splicing events were counted from the HISAT2 alignments with rMATS-turbo 4.3.0, run with all libraries as a single group and without its built-in group test (--statoff), for paired-end, reverse-stranded (fr-firststrand) reads of 151 nt with --variable-read-length and --allow-clipping, using junction-read counts only. For each event and knee, the percent spliced in (PSI) was calculated from length-normalized inclusion and skipping junction counts. Events were tested if they had at least 10 junction reads in at least 70% of the 92 knees, a mean PSI between 0.02 and 0.98 and a standard deviation of PSI above 0.02. Each event was modeled with one value per knee, so that a few high-coverage libraries could not dominate the estimate: knees with fewer than 10 junction reads were excluded, and PSI was clipped to [0.01, 0.99], logit-transformed and regressed by ordinary least squares on grade with sequencing batch and the z-scored number of detected genes, testing the grade coefficient with a two-sided t test. ΔPSI was the fitted PSI at grade 4 minus that at grade 0. P values were adjusted by the Benjamini-Hochberg method within each event type. An event was called robust if its adjusted p was < 0.05, |ΔPSI| was ≥ 0.05, and its grade coefficient had the same sign and p < 0.05 both with strandedness as an added covariate and after excluding the 6 flagged libraries; a cadaveric-only fit was reported but not required. Matrisome events were also adjusted separately. The per-knee model was chosen because, on simulated counts, it recovered all true events and called none of the high-depth decoys, whereas a quasibinomial model of pooled junction counts called all decoys, and apparent mutually exclusive exon events that paired two constitutive exons were excluded from interpretation (Supplementary Methods).

Exon usage was measured independently of rMATS on disjoint exonic bins of protein-coding and lncRNA genes, in the style of DEXSeq, counted with featureCounts. In genes of the differential expression universe with more than one annotated transcript, bins with a mean of at least 10 reads and nonzero counts in at least 70% of knees were analyzed; the logit share of each bin within its gene was regressed on batch, detected genes and grade with limma (eBayes, robust = TRUE), with gene-level p values by the Simes method. Targeted measures of *FN1* (EDB, EDA, IIICS and the 3′ end of *FN1-208*), *FGFR1-3* IIIb/IIIc, *CD44* variant exons and *FGF1* first exons were obtained from exonic, intronic and junction counts. Knees with fewer than 20 informative reads were excluded, measures were transformed to the empirical logit or log density ratio and regressed on grade with batch and detected genes, and p values were adjusted across the 15 targeted measures (these, a CS1 read-density measure and the *FN1-208* intron background control). Use of the IIICS 3′ splice site that retains the CS1 segment was taken from the genome-wide rMATS analysis, with its matrisome-adjusted p value.

### Bulk OARSI score analyses

The OARSI analysis used the genes, model and thresholds of the grade analysis, with the OARSI score in place of grade: DESeq2 with the design ∼ Batch + Detected_Genes_z + OARSI_Score, the score entered as a continuous variable and Detected_Genes_z computed over all 92 knees, with Wald tests, Benjamini-Hochberg adjustment and lfcShrink(type = “normal”). Gene set and motif enrichment used the ranking, settings, databases and seed described for grade. Directional agreement with the grade signature used unshrunk estimates. To test whether the OARSI score tracked the transcriptome within end-stage tissue, the same model was fitted to the 47 arthroplasty knees, all of macroscopic grade 4. As an independent test, genes associated with the score in the 44 scored cadaveric knees formed a signature that was scored in each arthroplasty knee and correlated with its OARSI score (Spearman), against a null distribution from 1,000 expression-matched random gene sets.

### Bulk robustness and sensitivity analyses

The following analyses are described in the Supplementary Methods. The grade model was refitted with the reverse-strand share or coding fraction as covariates, without the flagged libraries and in the high-quality libraries only (Figure S1a-b). The translation-related signal was scored per knee and tested with additional adjustment for RNA quality, tissue source and post-mortem interval, within each tissue source, and in the single-nucleus data (Figure S3). Early OA (grade 1.5-2) was compared with Normal (grade 0-1) with the grade-model covariates and a library-quality module covariate (Figure S1c-d). Co-expression modules were identified with WGCNA after removal of batch and RNA-quality effects (Figure S1e). Single-nucleus cluster signatures (Figure S1f) and Naba matrisome (10) category shares (Figure 6f) were scored in the bulk knees and regressed on grade. The grade signature was compared with GSE114007 (4) and GSE57218 (22). Enrichment of grade-associated genes among the effector genes of a genome-wide association meta-analysis of OA (Table S12 of ref. 23) was tested at the gene and the association-signal level. Departure from a linear response to grade was tested with a DESeq2 likelihood-ratio test of grade group as a factor against a linear grade term.

### Single-nucleus RNA-seq processing and clustering

Filtered gene-expression count matrices were read per sample. Nuclei with 1,000-25,000 unique molecular identifiers (UMIs) and less than 5% mitochondrial reads were retained (137,919 nuclei). No gene-count, complexity or computational doublet filter was applied; contaminating and artifactual populations were instead removed at the cluster level. Each sample was normalized with SCTransform (3,000 variable features), regressing the mitochondrial fraction.

Nuclear RNA over-represents long genes. Gene spans were computed from the Ensembl GRCh38 release 110 annotation, and a gene was defined as long when its span was at least eight times the median span of the annotated genes in the count matrix (13.8 kb), that is at least approximately 110 kb. Capture burden was defined, for each donor-cluster pseudobulk (for each donor in the neighborhood analysis), as the fraction of UMIs from genes with known span that fell in long genes lacking any matrix-, cartilage- or skeletal-related GO biological process annotation (4,149 of 4,379 long genes; Supplementary Methods), so that the covariate does not absorb matrix biology.

Samples were integrated with Seurat (SCT normalization, 50 dimensions, canonical correlation analysis anchors). Because variable features selected on the SCT residual variance were biased toward long genes, features were re-selected per sample by their residual variance above a loess fit of variance on gene span, and all quality-filtered nuclei were re-integrated. In this space, clusters of a fine clustering (resolution 1.0) were removed when their highest score for nine non-chondrocyte lineage panels exceeded 0.10 and their cartilage core score was below 0.05 (fibrotic and hypertrophic chondrocyte genes were deliberately excluded from the contaminant panels), and three osteoblastic or mural subclusters of one mixed cluster were removed on marker expression. Clusters of the resolution-0.1 partition failing marker-based admissibility criteria (a single top marker exceeding the next by a log2 fold change of at least 4, or of at least 1.5 when the top 20 markers had a median span at least eight times background; more than 50% of nuclei from one donor; or fewer than 50 nuclei), together with one cluster defined by the long gene *PCDH9*, were removed, leaving 108,600 chondrocyte nuclei, which were re-clustered (50 principal components, Louvain clustering at resolution 0.1, seed 42) into nine clusters: c0 *PRG4*+ superficial, c1 *ACAN*-high low-signal, c2 *COL10A1*+ hypertrophic, c3 pre-hypertrophic, c4 *CYTL1*+, c5 *ANXA1*+, c6 *SLC14A1*+, c7 *CRISPLD2*+ and c8 unassigned. Cluster markers (Figure 3c) were identified on the SCT assay with Wilcoxon tests (Bonferroni-adjusted p < 0.05), and marker pathway enrichment (Figure S9g) used over-representation analysis with long and ribosomal protein genes removed (Supplementary Methods). Markers computed on the RNA assay were compared with the normal-donor populations of Swahn et al. (8) by hypergeometric tests of marker overlap.

### Single-nucleus grade, stage and abundance analyses

For each cluster, raw counts were summed per donor (pseudobulk); donor-clusters with at least 20 nuclei were retained, clusters were tested when at least 8 donors and three grade values were available, and genes with at least 10 counts in at least 50% of the retained donors were tested. DESeq2 was fitted with ∼ technology + capture burden + macroscopic knee grade, grade as a continuous variable and capture burden centered within cluster (technology was dropped where it was not estimable), and the grade coefficient was tested with the Wald test with Benjamini-Hochberg correction (adjusted p < 0.05; DESeq2 default independent filtering). Log2 fold changes per grade unit are maximum-likelihood estimates. Pathway analysis ranked the Wald statistic z-scored within span deciles, after removal of cytosolic and mitochondrial ribosomal protein genes (and genes without a span annotation), in fgsea (Hallmark, GO biological process and Reactome; 15-500 genes). Genes were called shared when they had adjusted p < 0.05 in all nine clusters.

For the three-stage analysis, the same per-cluster pseudobulk model was fitted with ∼ technology + burden + stage, testing early OA versus Normal and severe OA versus Normal with Wald tests and any stage effect with a likelihood-ratio test against ∼ technology + burden. At least 10 donor-clusters were required, a cluster was skipped if Normal or severe OA had fewer than three donors, and the early-OA contrast was reported only with at least three early-OA donors. Severe-OA genes were classed by their early-OA effect as early onset (same sign, |early|/|severe| log2 fold change ≥ 0.5), progressive (same sign, ratio < 0.5) or late only (early-OA change of the opposite sign). Severe OA was also tested against early OA on the same fit, and severe-OA genes were further classed as established early (early OA against Normal adjusted p < 0.05 in the same direction, with no change from early to severe OA) or as changing with progression (severe against early OA adjusted p < 0.05 in the same direction) (Figure 8a-b). Over-representation of GO biological process (simplified at a similarity of 0.7), MSigDB Hallmark and NABA matrisome gene sets among the genes of each class, separately for rising and falling genes, pooled over clusters (a gene counted when it was in that class in at least two clusters, or in one where stated), was tested with clusterProfiler against all genes tested in at least one cluster (for c0 alone, against genes tested in c0; gene sets of 10 to 500 genes, NABA up to 2,000; Benjamini-Hochberg correction; Figure 8e). Class shares were recomputed with long genes excluded, and the enrichments with genes and background restricted to protein-coding genes that are not long.

For the timing of bulk changes, knees were grouped by grade (0-1, 23; 1.5, 10; 2-3, 8; 4, 51), variance-stabilized expression was adjusted for batch and the number of detected genes (limma removeBatchEffect), and each of the 1,350 grade-associated genes was classed by the fraction of its grade 4 change reached at grade 1.5 and grade 2-3: early, at least half at grade 1.5; late, less than a quarter at grade 2-3; mid-plateau, at least three quarters at grade 2-3; overshoot, a change at grade 1.5 or 2-3 exceeding the grade 4 change by at least half (rising overshoot genes were merged with mid-plateau); otherwise graded. Group means were given bootstrap 95% confidence intervals over knees within each group (Figure 8f). A likelihood-ratio test of the grade-group factor against a linear grade score (DESeq2, ∼ batch + detected genes + group) found no gene departing from linearity, and these classes are therefore descriptive.

Over-representation of GO biological process (simplified at a similarity of 0.7), Hallmark and NABA matrisome gene sets in each class and direction was tested with clusterProfiler against the 17,951 tested genes (gene sets of 10 to 500 genes for GO and 5 to 2,000 for Hallmark and NABA; Benjamini-Hochberg). A gene was considered low-count or sparsely detected if it was non-coding, detected in fewer than half of the knees or had a median normalized count below 10.

Cluster abundance was analyzed per donor on the complete 30 × 9 donor-by-cluster grid of nuclei counts. Centered log-ratio-transformed proportions (pseudocount 0.5 nuclei) were tested by Spearman correlation with grade and by a Kruskal-Wallis test across stages, against 10,000 donor-label permutations, with Benjamini-Hochberg correction across clusters. Shifts within clusters were tested with miloR (28) in neighborhoods of a k-nearest-neighbor graph (k = 30) built on the 50 principal components used for clustering, modeling per-donor neighborhood counts with an edgeR negative binomial generalized linear model (∼ technology + capture burden + macroscopic knee grade) and controlling the spatial false discovery rate at 0.1. Both analyses were repeated in the 24 multiome donors. Neighborhood log fold changes were correlated with a per-nucleus bulk grade-signature score (Spearman; Supplementary Methods).

To test whether grade responses differed between clusters (Figure 5), all donor-clusters with at least 20 nuclei (239 from 30 donors) were analyzed jointly on variance-stabilized counts of genes with at least 10 counts in at least 50% of donor-clusters. For each gene, a linear mixed model, technology + burden + cluster + cluster × grade + (1 | donor), was compared with a model with a single common grade slope by likelihood-ratio test (8 df), and each cluster’s slope was contrasted with the unweighted mean slope of the other eight (Wald z). A grade association was called cluster-distinctive when it was significant in that cluster in the per-cluster DESeq2 model, the interaction and the deviation both had adjusted p < 0.05, and the deviation and the cluster’s own mixed-model slope had the sign of the cluster’s DESeq2 log2 fold change. Cluster-distinctive pathways were identified by fgsea on the deviation z (adjusted p < 0.05) and also had to be significant with the same sign in that cluster’s grade GSEA (Supplementary Methods).

Ambient RNA and doublets were assessed as sensitivity analyses on all 137,919 quality-filtered nuclei. Counts were corrected with SoupX (29) and with DecontX (30), doublets were predicted with scDblFinder (31), and the per-cluster grade model was refitted unchanged on each corrected count set, with and without predicted doublets (Figure S11; Supplementary Methods).

### Cluster identity in healthy donors

Cluster identity was defined in the 12 donors with healthy cartilage (macroscopic knee grade ≤ 1; 51,443 nuclei), so that identity was not confounded with the OA response; markers, pairwise contrasts, signature enrichment, classification and pseudobulk recall were repeated in all 30 donors, and chromatin analyses in all 24 multiome donors. Marker genes were tested on donor × cluster pseudobulk with limma-voom (design ∼0 + cluster + donor, each cluster against the unweighted mean of the other eight; adjusted p < 0.05 and log2 fold change > 0.5). Clusters were placed on a zonal axis using gene sets from microdissected human knee cartilage (27) and on a hypertrophy axis (*RUNX2*, *COL10A1*, *IBSP*), and enrichment of zonal and published signatures from Ji et al. (6), Fan et al. (9) and Swahn et al. (8) was tested by gene set enrichment analysis. Recurrent gene programs were identified by non-negative matrix factorization in each healthy donor and combined into meta-programs following Gavish et al. (63). Separability was assessed by leave-one-donor-out classification and stability by reclustering subsamples. In the healthy multiome donors, cluster-specific accessible regions and chromVAR motif accessibility were tested in the same donor-blocked design, and motif enrichment among cluster-specific regions was tested with a Cochran-Mantel-Haenszel test stratified by peak GC content. The identity of c5 and c8 was also checked. Details are given in the Supplementary Methods.

### Matrisome analyses

Matrisome genes and categories were taken from the Naba matrisome (10) as distributed in MSigDB (1,026 genes, each in one category), and genes were also assigned to 24 function-defined families curated from explicit gene lists. Using the donor-by-cluster pseudobulks of the grade model, the share of each category or family was computed as its summed counts per million (CPM) as a percentage of the annotation total, excluding long Naba genes and showing *FN1* separately, and change with grade was estimated per cluster with share ∼ technology + capture burden + grade (Benjamini-Hochberg across all tests of an analysis). Bulk shares were computed in the same way from transcripts per million (TPM), because full-length bulk reads over-weight long transcripts, and regressed on grade with share ∼ batch + detected genes + grade. Per-gene grade effects came from the per-cluster pseudobulk model; matrisome genes that failed its expression filter but had at least 10 counts in at least three donors were refitted with the same design and size factors and called changed only if supported by a donor-level Spearman correlation. Family-level responses were scored as mean Wald statistics z-scored within span deciles. Details are given in the Supplementary Methods.

### Cell-cell communication

Communication between the nine chondrocyte clusters was inferred with CellChat (32,33) separately at each disease stage (Normal, 12 donors; early OA, 4 donors; severe OA, 14 donors), using the log-normalized RNA expression of the 108,600 chondrocyte nuclei. A cluster entered a stage’s analysis only if it contained at least 25 nuclei at that stage. The complete human CellChatDB was used without curation or added interactions. Communication probabilities were computed with computeCommunProb using the truncated mean (type = “truncatedMean”, trim = 0.1) as the average expression per cluster, with population.size = FALSE, so probabilities were not weighted by cluster size, and significance was assessed with 100 permutations (seed 42). We used the truncated mean instead of CellChat’s default trimean because the trimean is zero unless a gene is detected in roughly 25% of a cluster’s nuclei, a single-cell threshold that discards lower-expressed receptors in nuclear data. Links supported by fewer than 10 nuclei in either cluster were removed, and significant links (p < 0.05) were extracted at ligand-receptor and pathway level.

Network size was summarized per stage as the number of significant ligand-receptor links and the sum of their probabilities (Figure 7a). For the *FN1*, FGF and TENASCIN pathways, each ligand’s share of FGF probability (Figure 7c) and each ligand-receptor pair’s share of *FN1* probability (Figure 7d) were expressed as percentages of the stage total. Pathways were compared between stages with rankNet, which compares each pathway’s probabilities over all sender-receiver cluster pairs by a paired Wilcoxon signed-rank test. Because nuclei were pooled within each stage and the test units are sender-receiver cluster pairs, these p values do not reflect donor-to-donor variation, and they were not corrected across pathways. The percentage of the Normal-to-severe OA change already present at early OA was computed for the *FN1* receptor shares and for *FN1* mRNA (Figure 7e; Supplementary Methods). The breadth of sending and receiving was the effective number of clusters, exp(−Σ p_i ln p_i), where p_i is cluster i’s fraction of the pathway’s summed probability as source or target.

To obtain donor-level statistics, CellChat was run separately for each of the 30 donors with the same settings. A donor-level expression product of ligand and receptor expression served as a check independent of CellChat’s permutation filter. With the donor as the unit, each measure was correlated with grade (Spearman) and compared between Normal (12) and severe OA (14) donors (Wilcoxon rank-sum test), with Benjamini-Hochberg correction within each set of measures; the three *FN1* receptor shares were the confirmatory tests (Figure S21). As an independent, threshold-based method, MatriCom (34) was run at each stage on the same nuclei, clusters and 25-nucleus floor with the application defaults (a gene expressed by a population if at least 30% of its nuclei had non-zero expression and its mean non-zero log-normalized expression was at least 1); because the two methods use different databases, counting units and inclusion rules, only the direction of change between stages was compared. FGF ligand and receptor ratios, an ERK feedback target score and heparan sulfate modification genes were analyzed from the per-cluster pseudobulks with the covariate-adjusted grade model (Figures S18-S19; Supplementary Methods).

### Single-nucleus multiome

Paired snRNA-seq and snATAC-seq libraries from 24 of the 30 donors were analyzed with Seurat and Signac. Nuclei were kept if they had 1,000-25,000 RNA UMIs, less than 5% mitochondrial reads, nucleosome signal < 2 and transcription start site (TSS) enrichment > 1. Per-sample peak calls were merged into a unified set of 247,013 peaks, fragments were re-counted against this set, and nuclei with more than 500 fragments in peaks were kept (104,181 nuclei). RNA was integrated with SCT-based CCA anchors and the peaks assay by integrated latent semantic indexing (dimensions 2-40). Because the multiome RNA libraries are the same libraries as the snRNA-seq data for these donors, cluster labels were transferred by exact barcode join rather than by statistical label transfer, and the 81,670 labeled chondrocyte nuclei were used for all analyses.

Differential accessibility against grade was modeled on fragment counts summed per donor, in DESeq2 with ∼ macro_knee_grade, grade continuous, testing the grade coefficient with the Wald test and calling peaks at Benjamini-Hochberg-adjusted p < 0.05 (Figure 9b). Peaks were kept if they had at least 10 counts in at least 6 donors (25% of donors). Because donor was the unit of replication, it could not be included as a covariate, and tissue source could not be included because it is aliased with grade 4: all six arthroplasty knees are grade 4, and only one grade-4 knee is cadaveric. Robustness was checked by leave-one-donor-out refits. The same model was fitted per cluster (donor-clusters with at least 20 nuclei, clusters with at least 8 donors; Figure 9c).

Peaks were linked to genes with Signac LinkPeaks (peaks within 500 kb of the gene TSS, Pearson correlation of accessibility with SCT expression against 200 GC-, accessibility- and width-matched background peaks; absolute score > 0.05 and z-test p < 0.05, without correction for multiple testing), computed across all 81,670 nuclei without reference to grade or cluster. Links are correlations of accessibility with expression and do not demonstrate regulation. Motif accessibility was estimated with chromVAR for 633 human JASPAR2020 CORE matrices; per-nucleus deviation z-scores were averaged per donor within each cluster (at least 20 nuclei, at least 8 donors per cluster) and correlated with grade by Spearman correlation, with Benjamini-Hochberg correction across motifs within each cluster (Figure 9d). Correlations from individual nuclei were not used for inference, because they are pseudoreplicated. SMAD motif families were scored (donor-cluster units with at least 10 nuclei) as the mean chromVAR z of their member motifs from the cisTarget v10nr_clust collection (the SMAD2/3 family from eight motifs) and correlated with grade per cluster, with Benjamini-Hochberg correction across the nine clusters within each family; the JASPAR SMAD3 and SMAD2::SMAD3::SMAD4 motifs were taken from the per-cluster motif analysis (Figure 11c). Coverage at selected loci was drawn with Signac CoveragePlot grouped by stage (Figures S23-S24). For display only, donors were grouped into stages; all statistical tests used continuous grade.

### FN1, TGF-β/SMAD and candidate regulator analyses

Per-cluster grade associations of *FN1*, fibril-assembly genes, fibronectin-cleaving proteases, TGF-β pathway genes and *FN1* receptor genes were taken from the per-cluster snRNA-seq grade model (Figures 10a, 11a and 11e), and bulk estimates from the bulk continuous-grade model. Within each cluster, donor pseudobulks were variance-stabilized (DESeq2 vst, blind = TRUE) and target-set scores (a SMAD3 early-response “SMAD3 core” set with *FN1*, *COMP* and *LTBP2* excluded, Hallmark TGF-β signaling and a BMP target set) were computed as the mean z over expressed genes and regressed on grade with technology and capture burden. Coupling beyond grade was computed across donors within each cluster as a partial Spearman correlation: the ranks of both variables were regressed on rank(grade) (and, for the receptor analyses, capture burden and technology) and the residuals were correlated (Pearson), with Benjamini-Hochberg adjustment across clusters (Figures 10b-c and 11f). Accessibility of peaks linked to *FN1*, *HTRA1*, *ITGA5* and *CD44* was correlated with grade across the 24 multiome donors (Spearman; descriptive, not tested; Figure 10d); summed linked-region accessibility and RNA were summarized by stage (Figure 10e), and *FN1* linked-region accessibility was correlated with *FN1* RNA (Figure 10f). Candidate transcriptional regulators of *FN1* (33 curated factors and pathways plus data-driven additions) were scored for prior regulatory edges in CollecTRI and DoRothEA, and transcription factor activity was inferred with decoupleR (univariate linear model, CollecTRI regulons with *FN1* removed) in the snRNA-seq, bulk and multiome data. Candidates were ranked on four evidence categories: a prior regulatory edge, motif enrichment in the 19 *FN1*-linked opening regions, association of activity with grade, and coupling with *FN1* beyond grade (Figure S26a). Tn5 footprint depth at selected motifs was analyzed separately (Figure S26b-c). The coupling statistics used to rank candidate regulators are nominal and were not corrected for multiple testing (Supplementary Methods).

### Statistical analysis

Unless stated otherwise, tests were two-sided, multiple testing was controlled by the Benjamini-Hochberg procedure, and the significance threshold was adjusted p < 0.05. Donors, not nuclei, were the unit of replication for all single-nucleus inferences; each bulk sample was one knee from one donor. For bulk RNA-seq, reported log2 fold changes are shrunk with a normal prior (DESeq2 lfcShrink, type “normal”) and p values come from the unshrunk Wald test; calls that depend on the direction of change (robustness to library quality, replication and response shapes) used the unshrunk estimates. Single-nucleus log2 fold changes are unshrunk maximum-likelihood estimates. Peak-gene links, rankNet pathway comparisons and candidate regulator coupling are reported as nominal statistics. Because grade 4 cartilage was obtained almost entirely at arthroplasty, analyses in cadaveric knees only are reported as sensitivity analyses and were not required for significance. Random seeds were fixed (42 unless stated otherwise).

## Data and code availability

Raw and processed sequencing data analyzed in this study (bulk RNA-seq, single-nucleus RNA-seq and single-nucleus multiome) are derived from the donors included in datasets that will be available through the SPARC Portal (RRID:SCR_017041) upon publication at the following DOIs: https://doi.org/10.26275/ew6a-yepx (64) and https://doi.org/10.26275/sd1m-hckw (65). Raw sequencing data will be available through the database of Genotypes and Phenotypes (dbGaP) under accession phs004628.

## Software

Bulk RNA-seq reads were processed with FastQC 0.12.1, Trim Galore 0.6.10 (Cutadapt 5.1), HISAT2 2.2.1, samtools 1.22.1 and featureCounts (Subread 2.1.1 for gene counts and 2.0.3 for splice-junction and exon-bin counts); splicing events were quantified with rMATS-turbo 4.3.0, and read distribution and strandedness with RSeQC 5.0.4. Single-nucleus multiome libraries were processed with Cell Ranger ARC 2.0.2 (refdata-cellranger-arc-GRCh38-2020-A-2.0.0) and GEM-X single-nucleus RNA-seq libraries with Cell Ranger 9.0.1 (refdata-gex-GRCh38-2024-A); genes were matched across the two references by gene symbol (see Nuclei isolation, library preparation and sequencing). Downstream analyses were performed in R 4.3.1 with Seurat 5.1.0, SeuratObject 5.0.2, sctransform 0.4.1, Signac 1.13.0, chromVAR 1.24.0, motifmatchr 1.24.0, TFBSTools 1.40.0, JASPAR2020 0.99.10, BSgenome.Hsapiens.UCSC.hg38 1.4.5, EnsDb.Hsapiens.v86 2.99.0, ChIPseeker 1.38.0, TxDb.Hsapiens.UCSC.hg38.knownGene 3.18.0, DESeq2 1.40.2, limma 3.56.2, edgeR 3.42.4, lme4 1.1-35.4, fgsea 1.28.0, clusterProfiler 4.10.1, GOSemSim 2.28.1, msigdbr 7.5.1 (MSigDB 7.5.1), org.Hs.eg.db 3.18.0, RcisTarget 1.20.0, WGCNA 1.73, miloR 1.10.0, CellChat 2.1.2 (CellChatDB v2), MatriCom 1.0, decoupleR 2.8.0, dorothea 1.14.1, presto 1.0.0, glmnet 4.1.8, RcppML 0.3.7, SoupX 1.6.2, decontX 1.0.0, scDblFinder 1.16.0 and data.table 1.17.0; figures were drawn with ggplot2 3.5.1, patchwork 1.2.0, ggrepel 0.9.5 and ggrastr 1.0.2 and with matplotlib in Python. R package versions are those of the analysis environment.

## Acknowledgments

The authors thank Michelle Payad, Zaira Flores, Jessica Nitta, and the team at Lifesharing, San Diego for providing access to the precious gift of tissue donation from the local San Diego & Imperial communities that made the research possible. We thank the Scripps Research Genomics core members Tony Mondala, Stephen Head and Sean Wang for their support with omics data generation.

## Author contributions

M.K.L. and H.S. conceptualized this study. H.S. and M.K.L. drafted the manuscript, with contributions from the other authors. H.S. performed data analysis. All authors approved the final version of the manuscript and figures.

## The RE-JOIN Consortium

Armen Akopian, Kyle Allen, Alejandro Almarza, Benjamin Arenkiel, Yangjin Bae, Bruna Balbino de Paula, Anita Bandrowski, Mario Danilo Boada, Jacqueline Boccanfuso, Jyl Boline, Dawen Cai, Carpio, Dellina Lane, Robert Caudle, Racel Cela, Yong Chen, Rui Chen, Brian Constantinescu, Cortez, Ibdanelo, Yenisel Cruz-Almeida, M. Franklin Dolwick, Chris Donnelly, Zelong Dou, Joshua Emrick, Malin Ernberg, Danielle Freburg-Hoffmeister, Spencer Fullam, Janak Gaire, Akash Gandhi, Benjamin Goolsby, Stacey Greene, Nele Haelterman, Michael Iadarola, Shingo Ishihara, Azeez Ishola, Sudhish Jayachandran, Zixue Jin, Frank Ko, Priya Kulkarni, Zhao Lai, Brendan Lee, Yona Levites, Carolina Leynes, Jun Li, Martin Lotz, Lindsey Macpherson, Tristan Maerz, Camilla Majano, Anne-Marie Malfait, Maryann Martone, Bella Mehta, Richard Miller, Rachel Miller, Michael Newton, Alia Obeidat, Merissa Olmer, Dana Orange, Miguel Otero, Kevin Otto, Folly Patterson, Marlena Pela, Sienna Perry, Theodore Price, Hernan Prieto, Russell Ray, Dongjun Ren, Margarete Ribeiro Dasilva, Alexus Roberts, Elizabeth Ronan, Oscar Ruiz, Shad Smith, Mairobys Socorro, Kaitlin Southern, Joshua Stover, Michael Strinden, Hannah Swahn, Evelyne Tantry, Sue Tappan, Luis Tovias Sanchez, Airam Vivanco-Estela, Joost Wagenaar, Lai Wang, Kim Worley, Joshua Wythe, Jiansen Yan, and Julia Younis.

