## Supplemental Information for "Integrated transcriptomic and epigenomic profiling of human cartilage reveals coordinated changes in fibronectin expression, splicing, and receptor signaling during osteoarthritis progression"

for

### Contents

**Supplementary Methods**
**Figure S1.** Robustness of the grade analysis to RNA quality, comparison of early OA with Normal cartilage and other analyses of the bulk cohort.
**Figure S2.** The bulk grade signature is reproduced in direction in public cartilage data, is enriched at OA risk loci and is consistent with a linear change with grade.
**Figure S3.** The decline of translation-related transcripts with grade is a library-level feature of end-stage cartilage.
**Figure S4.** The cartilage transcriptome changes with the histological OARSI score as with macroscopic grade.
**Figure S5.** Macroscopic grade and OARSI score signatures, and the OARSI score within arthroplasty knees.
**Figure S6.** Genome-wide splicing in bulk cartilage: model comparison, matrix events and FN1 exon usage.
**Figure S7.** Cohort composition.
**Figure S8.** Cluster abundance by grade and neighborhood differential abundance.
**Figure S9.** Cluster identity: zonal, published-population, motif, separability and marker-pathway analyses.
**Figure S10.** Advancing OA is associated with a response shared by all nine chondrocyte clusters.
**Figure S11.** The shared grade response persists after ambient RNA correction and doublet removal.
**Figure S12.** Staging donors reproduces the continuous grade result.
**Figure S13.** Matrix composition per cluster: families, categories, cluster-restricted genes, bulk detail and between-cluster differences.
**Figure S14.** Matrisome genes expressed in each chondrocyte cluster and their change with grade.
**Figure S15.** CellChat network topology across stage.
**Figure S16.** MatriCom analysis supporting Figure 7.
**Figure S17.** Pathway and FN1-receptor networks by chondrocyte cluster.
**Figure S18.** FGF1 rises relative to FGF2 with grade without a rise in ERK output.
**Figure S19.** Expression of FGF1 export and heparan sulfate-modifying genes with grade.
**Figure S20.** CellChat ligand-receptor composition and sender/receiver detail.
**Figure S21.** Donor-level replication of the stage-level communication results.
**Figure S22.** Chromatin accessibility detail.
**Figure S23.** Chromatin accessibility at FN1 and TNFSF11 by stage.
**Figure S24.** Accessibility at the regions linked to ITGA5, ABCA1 and GPX3 falls with grade.
**Figure S25.** Hyaluronan-axis genes and the donor-level CD44-ITGA5 axis.
**Figure S26.** Candidate transcriptional regulators of FN1: evidence and footprinting.
**Table S1.** Provided as a separate Excel file (Table_S1.xlsx).

### Supplementary Methods

*Full description of all analyses. Reference numbers refer to the reference list of the main text. The main-text Methods summarize these procedures.*

#### Human cartilage samples

Described in the main Methods.

#### Nuclei isolation, library preparation and sequencing

Nuclei were isolated from cartilage stored in liquid nitrogen using the Singulator S200 Automated Tissue Dissociation System (S2 Genomics). The procedure was performed according to a previously published protocol (protocols.io, https://doi.org/10.17504/protocols.io.ewov193molr2/v1). Before loading into the Singulator, 100-150 mg of frozen tissue was finely shaved and minced into small pieces using a scalpel on a pre-chilled petri dish. The tissue fragments were then transferred to a pre-cooled Singulator cartridge containing 75 µL of RNase inhibitor (40 U/µL). The cartridge was placed into the Singulator, and the low-volume nuclei isolation program was run. Following homogenization, the resulting suspension was passed through 20 µm and 40 µm strainers to remove tissue and cellular debris. The flow-through was collected into a 5 mL LoBind tube and centrifuged at 150 × g for 10 min at 4 °C. The supernatant was carefully removed, and the nuclei pellet was resuspended in 600 µL of PBS containing 0.02 mg/mL recombinant albumin and 1 U/µL RNase inhibitor. To further eliminate cellular debris, a second centrifugation was performed at 150 × g for 10 min at 4 °C. The supernatant was discarded, and the nuclei pellet was resuspended in 100 µL of diluted nuclei buffer. For nuclei counting and quality assessment, 18 µL of the nuclei suspension was mixed with 2 µL of acridine orange/propidium iodide (AO/PI) stain. Nuclei were counted using the Countess II FL Automated Cell Counter (AO/PI staining). Nuclear morphology and membrane integrity were also inspected under a microscope at 40× or 60× magnification (AO/PI staining), following the nuclear integrity check guidelines provided by 10x Genomics. Based on the estimated concentration, the nuclei were diluted in chilled diluted nuclei buffer to a target concentration of 4,000-5,000 nuclei/µL for a final recovery of 20,000 nuclei in 5 µL. Libraries were prepared with the Chromium Next GEM Single Cell Multiome ATAC + Gene Expression kit (10x Genomics, PN-1000283), with 33,000 nuclei loaded per reaction, and sequenced paired-end on an AVITI system (Element Biosciences). GEM-X library details are given in the main Methods. Multiome libraries (24 donors) were aligned and counted with Cell Ranger ARC 2.0.2 against the refdata-cellranger-arc-GRCh38-2020-A-2.0.0 reference, and single-nucleus RNA-seq libraries from the remaining six arthroplasty donors (Chromium GEM-X Single Cell 3′ v4) with Cell Ranger 9.0.1 against refdata-gex-GRCh38-2024-A; intronic reads were included in both.

#### Bulk RNA sequencing

Total RNA was extracted from knee articular cartilage. For RNA isolation, 100 mg of frozen cartilage was homogenized in a precooled mortar using a pestle, followed by further homogenization in a Precellys homogenizer at 6,500 rpm for four 20-s cycles. RNA was isolated using TRIzol reagent (Thermo Fisher Scientific), followed by phase separation in chloroform and ethanol precipitation. After isolation, on-column DNase treatment (Zymo) was performed using DNase I (QIAGEN) and the RNeasy MinElute Cleanup Kit (QIAGEN). RNA concentration and purity were assessed by NanoDrop spectrophotometry (A260/A280 and A260/A230 ratios), and integrity was assessed on an Agilent 4200 TapeStation (Agilent Technologies). RNA libraries were prepared using the QIAseq Stranded mRNA Select Low Input Kit (QIAGEN, cat. 334205) with ribosomal RNA depletion (cat. 334235). Libraries were spiked with 5% PhiX control and sequenced on an Illumina NovaSeq X (Illumina) using a 25B flow cell with 300-cycle chemistry, generating 2 × 151-bp paired-end reads, to a target depth of approximately 30 million read pairs per sample. Library strandedness was reverse (dUTP-type), as confirmed with RSeQC infer_experiment (median 94% of strand-informative read pairs in the reverse orientation across the analyzed cartilage libraries). Libraries were sequenced in two batches (81 and 11 of the analyzed knees) (see main Methods).

#### Read processing, alignment and quantification

Adapters and low-quality bases were trimmed with Trim Galore 0.6.10 (Cutadapt 5.1) in paired mode with the adapter sequences AGATCGGAAGAGCACACGTCTGAACTCCAGTCA (read 1) and AGATCGGAAGAGCGTCGTGTAGGGAAAGAGTGT (read 2) given explicitly. The first 16 bases of read 2 were removed (--clip_R2 16). All other settings were defaults (Phred quality cutoff 20, minimum read length 20 bp, adapter overlap stringency 1 bp). Trimmed reads were aligned to GRCh38 with HISAT2 2.2.1 using the prebuilt HISAT2 grch38 genome index (no splice-site annotation) and the --dta option, without a strandedness setting. Alignments were sorted with samtools 1.22.1. Gene-level counts were obtained with featureCounts (Subread 2.1.1) against the GENCODE v44 comprehensive annotation (primary chromosomes; gene models identical to Ensembl release 110). Counting was on exon features summarized by gene_id, reverse-stranded (-s 2), paired-end (-p), and both mates of a pair had to align (-B). Chimeric pairs were not counted (-C), and reads rather than fragments were counted. The index and the annotation named the mitochondrial chromosome differently, so mitochondrial genes received no counts; they were removed downstream in any case. Transcripts per million (TPM) were computed using as gene length the union of the gene's exons in the annotation (non-overlapping exonic length, as reported by featureCounts). Per-library metrics were the number of read pairs before trimming, the percentage of reads aligned, the percentage of reads assigned to genes and the number of assigned reads. The number of detected genes was the number of the 62,700 annotated genes with at least one count.

#### Sample and gene filtering

Libraries were retained if at least 10% of reads were assigned to genes and at least 5 million reads were assigned. No analyzed cartilage library failed these thresholds; seven further libraries were not analyzed (see main Methods). The analyzed cohort comprised 92 knees: grade 0 (n = 4), 1 (n = 19), 1.5 (n = 10), 2 (n = 5), 3 (n = 3) and 4 (n = 51). Of these, 45 were from cadaveric donors and 47 from total knee arthroplasty. Genes on chromosome Y or the mitochondrial chromosome (assigned with org.Hs.eg.db) and *XIST* were removed (62,396 genes remained). Genes were then retained if their TPM exceeded 1 in at least 25% of the cartilage libraries. This left 17,951 genes, which formed the gene universe for all bulk analyses below.

#### RNA quality assessment

Read distribution and strandedness were measured for every BAM file with RSeQC (read_distribution.py and infer_experiment.py, default settings) using a BED12 gene model built from the same GENCODE v44 annotation (chromosome names matched to the alignment index). RSeQC's feature categories overlap, so reads were partitioned as follows: coding (CDS exons), UTR (5' and 3' UTR exons), intronic, and intergenic (total tags minus the other three). The percentage of coding reads was CDS tags divided by total tags. Strandedness was summarized as the reverse-strand share, the fraction of strand-informative reads in the reverse-stranded configuration ("1+-,1-+,2++,2--") among those in either configuration. Low-quality libraries had near-unstranded, intergenic-rich profiles, consistent with genomic DNA contamination. A library was flagged if its reverse-strand share was below 0.75 or its coding fraction below 10%; 6 of the 92 cartilage libraries were flagged. High-quality libraries were defined more strictly as reverse-strand share of at least 0.85 and coding fraction of at least 30% (75 knees). Flagged libraries were kept in the primary analyses, and their influence was assessed in the sensitivity analyses described below.

#### Principal component analysis

Counts of the 17,951 genes were transformed with the DESeq2 variance-stabilizing transformation (blind = TRUE, design ~1, size factors estimated on cartilage samples only). Sequencing batch and the number of detected genes were then removed with limma removeBatchEffect, protecting donor category (Normal, Aged, OA, arthroplasty) in the design matrix. Principal components were computed with prcomp (centered, not scaled) on the 500 genes with the highest variance after correction. The association of PC1 with grade is reported as the squared Pearson correlation ($R^2 = 0.36$; PC1 explained 41.2% of the variance) (Figure 1a).

#### Differential expression with continuous grade

Differential expression was tested with DESeq2. Grade was entered as a continuous variable with the design ~ Batch + Detected_Genes_z + Macro_Knee_Grade, where Detected_Genes_z is the z-scored number of detected genes (z-scored across the samples of the original multi-tissue run; because this differs from z-scoring within the 92 knees only by a linear rescaling, the grade estimates are identical). Genes with zero total counts were removed before fitting (none in this set). Size factors, dispersions and the negative binomial GLM were fitted with DESeq() defaults. The grade coefficient was tested with the Wald test, and p values were adjusted with the Benjamini-Hochberg method using results() defaults (independent filtering, alpha = 0.1, Cook's distance outlier handling); no gene received a missing adjusted p value. Reported log2 fold changes per grade unit were shrunk with lfcShrink(type = "normal"). P values and adjusted p values come from the unshrunk Wald test, because shrinkage does not alter them. Genes were called differentially expressed at an adjusted p value < 0.05. This gave 1,350 genes, 618 rising and 732 falling with grade (Figure 1b). For the downstream gene lists used in motif enrichment, we additionally required a mean normalized count (baseMean) of at least 10 (618 rising, 728 falling).

#### Gene set enrichment analysis

Genes were ranked by $-\log10(p) \times \text{sign}(\log2 \text{ fold change})$, using the nominal Wald p value of the grade coefficient (p values floored at $10^{-300}$). Ensembl IDs were mapped to gene symbols. Where several IDs mapped to one symbol, the ID with the largest absolute ranking statistic was kept, giving 17,923 ranked genes. The sign was taken from the shrunk log2 fold change. Shrinkage reversed the sign of 598 genes, all with nominal $p > 0.08$, whose ranks lie near zero; none was in the leading edge of an enriched term. Gene set enrichment analysis was run with clusterProfiler gseGO on GO Biological Process terms (org.Hs.eg.db, keyType SYMBOL). Settings were gene set size 10-500, exponent 1, the fgsea algorithm with eps = 0 and a fixed seed (set.seed(42), seed = TRUE), Benjamini-Hochberg adjustment, and all terms retained (pvalueCutoff = 1). Terms with adjusted $p < 0.05$ were considered enriched.

Figure 1c shows rising and falling terms chosen as follows. For rising terms, enriched terms with at least 30 genes were ordered by normalized enrichment score (NES). A term was dropped if its leading-edge genes overlapped those of a higher-ranked term with a Jaccard index above 0.7, and the first ten remaining terms were shown. For falling terms, ten enriched terms were chosen to represent the distinct falling programs, two per program (translation and ribonucleoprotein biogenesis, chromatin and RNA processing, glucocorticoid response, immune response, lipid metabolism and oxidative stress). Terms naming the negative regulation of a process were avoided. For Figure 1d, four rising and four falling terms were chosen, and for each the ten leading-edge (core enrichment) genes with the largest absolute shrunk log2 fold change were plotted.

#### Transcription factor motif enrichment

Motif enrichment was tested with RcisTarget separately on the gene symbols rising and falling with grade (adjusted $p < 0.05$, baseMean ≥ 10). The motif ranking database was hg38 RefSeq r80, 10 kb upstream and downstream of the transcription start site (mc9nr, genes-vs-motifs rankings). Motif annotations were the RcisTarget version 9 HGNC annotation (motifAnnotations_hgnc_v9), matching the ranking database. cisTarget was run with defaults: area under the recovery curve computed over the top 5% of the ranking, NES threshold 3, high-confidence annotations = direct and orthology-inferred. For each direction, motifs with NES > 3 were collapsed to their high-confidence transcription factors. Each factor was summarized by its maximum NES, maximum AUC and number of enriched motifs. The 12 factors with the highest maximum NES per direction are shown (Figure 1e), excluding *MYLK*, which is not a transcription factor, so that 11 are shown for rising genes. A factor was marked as differentially expressed if it was itself among the genes changing with grade in the same direction.

#### Robustness of the grade analysis to RNA quality

The grade model was refitted in four ways, each restricted to the same 17,951 genes: (i) adding the z-scored reverse-strand share as a covariate; (ii) adding the z-scored coding fraction as a covariate; (iii) excluding the 6 flagged libraries (n = 86); and (iv) restricting to the 75 high-quality libraries. In each refit, Detected_Genes_z was recomputed within the knees analyzed. Log2 fold changes were shrunk with lfcShrink(type = "normal") as in the primary analysis. For each refit we computed, over the 1,350 genes of the primary analysis, (a) the percentage still at adjusted $p < 0.05$ and (b) the percentage with the same sign as the primary model and $p < 0.05$ (Figure S1a). Same-sign calls compared unshrunk (maximum-likelihood) log2 fold changes, because normal-prior shrinkage can reverse the sign of near-zero estimates. Agreement of effect sizes was the Spearman correlation of shrunk log2 fold changes between the primary and refitted models over the same 1,350 genes ($\rho$ = 0.93 to 0.99 across the four refits; Figure S1b shows the refit restricted to high-quality libraries).

#### Sensitivity of the translation-related signal

For each knee, cytosolic ribosomal-protein genes (77 RPL/RPS genes, pseudogenes excluded), the leading-edge genes of cytoplasmic translation and the ribosome biogenesis and RNA splicing GO gene sets were scored as the mean z-scored variance-stabilized expression, and the ribosomal-protein fraction of counts was also computed. Scores were regressed on grade with the covariates of the grade model (batch and z-scored detected genes) and then additionally adjusted for RNA quality (RSeQC reverse-strand share and percentage of coding reads, percentages of assigned and aligned reads), for tissue source (arthroplasty or organ donor) and for post-mortem interval (PMI, or time from surgery to processing for arthroplasty knees, from the donor records; available for all 92 knees), and the analysis was repeated after excluding libraries flagged by RSeQC. Because PMI was much shorter in arthroplasty knees (0.5-6 h, one knee 19.75 h) than in organ donors, its effect was also tested within each source: within cadaveric donors (score ~ PMI + grade, and grades 0-3 only) and within arthroplasty knees, all of grade 4, against the OARSI score (score ~ PMI + OARSI score, with and without the knee with the longest interval). Grade 4 knees from organ donors and from arthroplasty were compared directly. The DESeq2 grade model and gseGO were refitted with the same covariates and subsets (Figure S3). In the single-nucleus data, the grade effects of the same ribosomal-protein genes were summarized per cluster from the per-cluster pseudobulk models with and without the capture-burden covariate.

#### Early OA versus Normal

Knees of grade 0-1 (Normal, n = 23) were compared with knees of grade 1.5-2 (early OA, n = 15) using the same model as Figure 1. The design was ~ Batch + Detected_Genes_z + group, with Detected_Genes_z standardized over all 92 knees, on all 17,951 genes. Genes with zero total counts in these 38 knees were removed. Testing used the Wald test with Benjamini-Hochberg adjustment. Reported log2 fold changes were shrunk with lfcShrink(type = "normal"), and all calls used the unshrunk estimates. A gene was robust if it had adjusted $p < 0.05$ and kept the same sign at $p < 0.05$ in two refits: one with the z-scored reverse-strand share added, and one with the flagged libraries removed.

Most genes at adjusted $p < 0.05$ were large, on-off decreases. We defined the 287 genes with adjusted $p < 0.05$ and an unshrunk log2 fold change below −3 as a library-quality module. For each knee, the module score was the fraction of module genes expressed at ≥ 1 count per million (CPM, relative to all assigned reads). Its association with the number of detected genes was tested by Spearman correlation across all 92 knees (Figure S1d). The early-OA model was then refitted with the module score (z-scored within the 38 knees) as an additional covariate. For non-module genes, p values were adjusted by Benjamini-Hochberg again within the non-module genes. A non-module gene was called robust if it reached adjusted $p < 0.05$ in this refit, kept the sign of the primary model, and also met the robustness criteria above. Robust genes expressed (CPM ≥ 1) in five or fewer of the 38 knees were marked separately (Figure S1c).

#### Weighted gene co-expression network analysis

To prevent noise from low-quality libraries from forming modules, the network was built from the universe genes expressed at CPM ≥ 1 in at least 25% of the 75 high-quality libraries (14,669 genes). Counts for all 92 knees were transformed with the variance-stabilizing transformation (blind = TRUE). Batch and the standardized reverse-strand share, coding fraction and number of detected genes were removed with limma removeBatchEffect, protecting grade in the design matrix. The 5,000 genes with the highest median absolute deviation were analyzed with WGCNA. The soft-thresholding power was the lowest power (tested 1-10 and 12-20 in steps of 2) at which the signed scale-free topology fit $R^2$ reached 0.8, which was 9. Modules were detected with blockwiseModules: signed network and signed topological overlap, minimum module size 30, deepSplit 3, merge cut height 0.2, maximum block size 6,000, pamRespectsDendro = FALSE, random seed 1. This gave eight modules plus unassigned (gray) genes. Module eigengenes were correlated with grade and with arthroplasty status (Spearman; Benjamini-Hochberg across all module × trait pairs, including RNA-quality and donor covariates) in all knees, and with grade in cadaveric knees only (Spearman). For cadaveric knees with a recorded post-mortem interval, standardized eigengenes were also regressed on grade, post-mortem interval and age. Hub genes were ranked by signed module membership (kME). Module gene sets were tested for over-representation of MSigDB Hallmark, Naba matrisome (NABA_) and Reactome gene sets with fgsea fora (minimum set size 10, universe = the 5,000 network genes, Benjamini-Hochberg) (Figure S1e). Cadaveric-only estimates are reported for information and were not used as a filter, because grade 4 cartilage was obtained almost entirely at arthroplasty.

#### Single-nucleus cluster signatures in bulk cartilage

Signatures of the nine snRNA-seq clusters (c0 *PRG4*+ superficial, c1 *ACAN*-high low-signal, c2 *COL10A1*+ hypertrophic, c3 pre-hypertrophic, c4 *CYTL1*+, c5 *ANXA1*+, c6 *SLC14A1*+, c7 *CRISPLD2*+, c8 unassigned) were taken from the snRNA-seq cluster markers (Seurat FindAllMarkers on the RNA assay, Wilcoxon test, positive markers only, min.pct 0.10, log fold-change threshold 0.25, at most 2,000 nuclei per cluster). Markers were kept if their adjusted p was < 0.05, their average log2 fold change was ≥ 0.5, they were a marker of a single cluster only, and they were in the bulk gene universe. Up to 50 markers per cluster were kept, ranked by (pct.1 − pct.2) × average log2 fold change. This gave 3 (c1) to 50 genes per cluster. Bulk counts were transformed with the variance-stabilizing transformation (blind =

TRUE), batch was removed with limma removeBatchEffect, and each gene was z-scored across the 92 knees. A knee's signature score was the mean z-score of the signature genes. Each score was standardized and regressed on grade with lm(score ~ Batch + Detected_Genes_z + grade). The resulting grade coefficient is the change in standard deviations per grade unit, with Benjamini-Hochberg adjustment across the nine clusters. The same model was fitted in cadaveric knees only, with the score and detected genes re-standardized within that subset, and is reported with nominal p values (Figure S1f).

#### Matrisome composition in bulk cartilage

Matrisome genes and categories (collagens, proteoglycans, ECM glycoproteins, ECM regulators, ECM-affiliated, secreted factors) were taken from the Naba matrisome gene sets (10) in MSigDB (msigdbr). Each gene was assigned to one category, and *FN1* was separated from the other ECM glycoproteins. The gene set was the same as in the single-nucleus analysis. It was restricted to genes detected in the snRNA-seq pseudobulks and in the bulk gene universe and excluded long genes (gene span at least 8 times the 13.8-kb background) (see single-nucleus methods). Bulk expression of these genes was taken as transcripts per million (TPM; counts divided by the union exonic length of the gene), because in full-length bulk RNA-seq read-based shares (CPM) over-weight long transcripts such as FN1. For each knee, each category's share was its summed TPM as a percentage of total matrisome TPM; curated family shares (Figure S13c) were computed in the same way. Shares were regressed on grade with lm(share ~ Batch + Detected_Genes_z + grade), and grade p values were adjusted by Benjamini-Hochberg across the seven categories (Figure 6f).

#### Splice junction and exon-bin counting

Alternative splicing was quantified from the HISAT2 alignments (GRCh38) of the cartilage bulk RNA-seq libraries described above. All 99 available cartilage BAM files were processed, including the seven libraries that were not analyzed (see main Methods). The 92 knees included in the differential expression analysis were then selected for every statistical test. Because the alignments used Ensembl-style chromosome names, the GENCODE v44 comprehensive annotation was converted to the same naming (the "chr" prefix removed; chrM renamed MT). Splicing events were enumerated and counted with rMATS-turbo 4.3.0, run with all libraries as a single group (--b1 only) and without its built-in group test (--statoff). The settings were paired-end reads (-t paired), a reverse-stranded library (--libType fr-firststrand), a read length of 151 nt with --variable-read-length (to allow for adapter-trimmed reads), and --allow-clipping. rMATS returned per-library inclusion (IJC) and skipping (SJC) junction counts for 87,936 skipped-exon (SE), 9,228 mutually exclusive exon (MXE), 9,586 alternative 5′ splice site (A5SS), 14,613 alternative 3′ splice site (A3SS) and 9,975 retained-intron (RI) events. Only junction-read counts (JC) were used.

To measure exon usage independently of rMATS, every protein-coding and lncRNA gene in GENCODE v44 was split into disjoint exonic bins at all annotated exon boundaries, in the style of DEXSeq. This produced 666,320 bins in 38,912 genes. Reads were counted per bin with featureCounts (Subread 2.0.3), using the settings of the gene-level counts: -s 2 -p -B -C, plus -f -O for feature-level counting with multi-overlap assignment and -J for splice-junction counts. For the targeted analyses of *FN1*, *FGFR1*-3, *CD44* and *FGF1*, a second featureCounts run used the same settings on exonic and intronic bins of 11 genes (*CD44*, *FGF1*, *FGF2*, *FGFR1*, *FGFR2*, *FGFR3*, *FN1*, *ITGA5*, *ITGB8*, *SULF1* and *SULF2*; 694 exonic and 228 intronic bins). Junction reads for these genes were taken from the -J output, and each junction was identified by the last base of the upstream exon and the first base of the downstream exon.

#### Genome-wide splicing events against grade

For each event and knee, the percent spliced in (PSI) was calculated from the length-normalized junction counts as (IJC/IncFormLen)/(IJC/IncFormLen + SJC/SkipFormLen). Events were tested only if they had at least 10 junction reads (IJC + SJC) in at least 70% of the 92 knees, a mean PSI between 0.02 and 0.98, and a standard deviation of PSI across knees above 0.02. These criteria were met by 3,657 events: 1,340 SE, 1,069 MXE, 231 A5SS, 401 A3SS and 616 RI. Each event was modeled with one value per knee, so that a few high-coverage libraries could not dominate the estimate. Within an event, knees with fewer than 10 junction reads were excluded; tested events had 65 to 92 informative knees. PSI was clipped to [0.01, 0.99], logit-transformed and regressed by ordinary least squares on macroscopic knee grade (0 to 4, treated as continuous), with the covariates of the differential expression model: sequencing batch (two levels) and the number of detected genes (genes with at least one read in the gene count matrix, z-scored). The grade coefficient was tested with a two-sided t test. ΔPSI was the fitted PSI at grade 4 minus the fitted PSI at grade 0, with the other covariates held at their means. P values were adjusted by the Benjamini-Hochberg method separately within each event type (genome-wide adjusted p).

Each event was refitted in three sensitivity analyses. The first added library strandedness as a covariate: the RSeQC fraction of reads in the expected reverse-strand orientation, z-scored. The second excluded the 6 libraries flagged for low quality (strandedness $< 0.75$ or $< 10\%$ of reads in coding sequence; 86 knees remained). The third used the 45 cadaveric knees only. In each subset, the number of detected genes was re-standardized. An event was called robust if its adjusted p was $< 0.05$, $|\Delta PSI|$ was $\geq 0.05$, and its grade coefficient had the same sign and $p < 0.05$ in both the strandedness-adjusted and the flagged-excluded fits. The cadaveric-only fit was reported but not required, because 47 of the 51 grade 4 knees came from arthroplasty, so grade and tissue source are largely aliased at the top of the scale.

Matrix genes were defined as the MSigDB NABA_MATRISOME set (core matrisome and matrisome-associated genes, from msigdbr 7.5.1). The 309 tested events in 90 matrisome genes were also assessed separately. For this focused test, Benjamini-Hochberg adjustment was applied across those 309 events, with all event types pooled, and the same robustness criteria were applied to the matrisome-adjusted p. Apparent MXE events in *COL1A2*, *COL2A1*, *COL3A1* and *FN1* were inspected by plotting the junction reads that supported each of the two exons in every knee. In each case both exons were supported by junction reads in most knees (at least 10 reads on each in 54 to 75 of the 92 knees for the collagen events and in all 92 for *FN1*), showing that rMATS had paired two constitutive exons. These events were therefore excluded from interpretation. For display, PSI per knee was plotted against grade with an unadjusted least-squares line.

The results were compared with two alternatives. First, the coverage requirement was relaxed to at least 10 junction reads in 50% of knees (9,536 events), with the same model. Second, a quasibinomial generalized linear model was fitted to the pooled junction counts (cbind(IJC, SJC)) with the same covariates and an offset of log(IncFormLen/SkipFormLen). This model included all knees with at least one junction read, used a dispersion of max(1, Pearson $\chi^2$/residual df) and applied Benjamini-Hochberg adjustment within event type. Before the analysis was applied to real data, the per-knee model was checked on simulated counts for the 92 knees. The simulation had 300 events: 30 had a true grade effect, and 10 decoys had no grade effect but 60-fold depth and high PSI in the four highest-grade knees. The per-knee model recovered 30 of 30 true events and called none of the decoys. The pooled-read model called all 10 decoys.

#### Exon-bin usage model

Exon-bin usage was analyzed in genes that were in the differential expression universe (17,951 genes) and had more than one annotated transcript. Bins had to have a mean of at least 10 reads and nonzero counts in at least 70% of knees, and genes had to keep at least two bins. This left 62,662 bins in 5,663 genes. For each knee, the share of a bin was calculated as (bin count + 0.5)/(sum of the gene's bin counts + 1) and logit-transformed. Values were set to missing for knees with fewer than 200 reads on the gene (4.1% of values). Logit shares were regressed on batch, the number of detected genes and grade with limma (lmFit and eBayes with robust = TRUE). Benjamini-Hochberg adjustment was applied across bins, and gene-level p values were obtained by the Simes method over each gene's bins and then adjusted across genes. Because featureCounts was run with multi-overlap assignment, a read that spanned several bins was counted in each of them. Bin shares are therefore relative measures of usage rather than isoform fractions.

#### Targeted *FN1* isoform measures

Five *FN1* segments were quantified for each of the 92 knees from the targeted counts (GRCh38 coordinates on chromosome 2; *FN1* is on the minus strand):

EDB (EIIIB; 215,392,931-215,393,203) and EDA (EIIIA; 215,380,811-215,381,080). Inclusion reads were the mean of the two inclusion junctions (upstream exon to cassette exon, and cassette exon to downstream exon). Exclusion reads were the skipping junction. Inclusion was expressed as the share of junction reads.

IIICS (V) region (215,372,016-215,372,375). Read density over the region, and separately over the CS1 segment (215,372,301-215,372,375), was expressed relative to the density over two constitutive exons (215,373,322-215,373,411 and 215,375,214-215,375,393; 270 bp).

*FN1-208* 3′ end. For the 3′ end unique to the truncated transcript *FN1-208* (ENST00000426059; 176 bp, 215,414,661-215,414,836), read density was expressed relative to the adjacent shared exon (215,414,837-215,414,958). The remainder of the same intron (4,546 bp) served as a background control.

Use of the IIICS 3′ splice site that retains the 25 codons encoding the α4β1-binding CS1 segment (A3SS, 215,372,301-215,372,375) was taken from the genome-wide rMATS analysis described above.

For each segment, knees with fewer than 20 informative reads were excluded. Junction measures were transformed to the empirical logit, log((inclusion + 0.5)/(exclusion + 0.5)). Density measures were transformed to the log density ratio, computed with 0.5 reads added to each count before dividing by segment length. Each transformed measure was regressed on grade with batch and the number of detected genes as covariates, in the same way as the genome-wide events. The same three sensitivity fits were run (with strandedness as a covariate, without flagged libraries, and in cadaveric knees only). A fit required at least 12 knees and at least three distinct grades. P values were adjusted by the Benjamini-Hochberg method across the 15 targeted measures (the *FN1*, *FGFR1*-3, *CD44* and *FGF1* measures below, the CS1 read-density measure and the *FN1-208* intron background control) separately within each fit. For the CS1 site, the matrisome-adjusted p from the genome-wide analysis is reported. Its standard error was derived from the reported p value and t distribution (residual df = number of knees − 4). Confidence intervals were drawn as β ± 1.96 × standard error.

For the sashimi-style views of EDB and EDA, each bin's read density was divided by the mean density of the two flanking constitutive exons and then averaged across knees of grade 0-1 (23 knees) and grade 3-4 (54 knees). Junction arcs were labeled with the median share of junction reads that skipped or included the exon in each group. Transcript models were taken from GENCODE v44.

#### Targeted FGFR, *CD44* and *FGF1* splice measures

The same per-knee framework was applied to the following measures:

*FGFR1*, *FGFR2* and *FGFR3* IIIb/IIIc. The IIIb share was calculated as the mean of the two IIIb junctions (from the upstream immunoglobulin-like domain III exon and into the transmembrane exon) divided by the sum of that value and the corresponding mean of the IIIc junctions.

*CD44* variant exons. Inclusion of the v6-v10 region was taken as half the sum of the junctions from constant exon 5 into the variant region, and within it, against the exon 5 to exon 15 skipping junction. The v2-v5 exons had no junction reads.

*FGF1* first exons. The share of reads from each of five alternative first exons was calculated among all first-exon reads joining the shared exon at chr5:142,614,161. The main first exon starts at 142,685,957.

The numbers of knees that passed the 20-read threshold were 85 for *FGFR1*, 57 for *FGFR2*, 27 for *FGFR3*, 77 for *CD44* and 38 for *FGF1*. *FGFR3* was not displayed. The cadaveric-only fit was not estimable for *FGF1* or *FGFR3*.

#### Replication in public cartilage datasets

The grade signature (the genes at adjusted $p < 0.05$ in the continuous-grade bulk model, with their shrunk log2 fold changes) was compared with two public datasets. GSE114007 (4; knee cartilage RNA-seq; 18 normal and 20 OA knees, sequenced on two platforms) was re-analyzed from the deposited raw counts with DESeq2 (~ platform + condition), and its OA-versus-normal log2 fold changes were shrunk with a normal prior as for the bulk cohort. GSE57218 (22; RAAK study; Illumina HumanHT-12 v3 arrays of lesioned and macroscopically preserved cartilage from 33 OA joints) was analyzed from the normalized series matrix with limma as a paired design (~ donor + status), with probes collapsed to the highest-expressed probe per gene symbol. For each dataset, we report the percentage of grade-signature genes with the same sign, the percentage that were also at adjusted $p < 0.05$ with the same or the opposite sign, and the Spearman correlation of log2 fold changes over all shared genes. Signs used for these percentages were taken from the unshrunk estimates, because normal-prior shrinkage can reverse the sign of estimates close to zero.

#### Enrichment at OA genetic risk loci

The 77 high-confidence effector genes of a genome-wide association meta-analysis of OA (Table S12 of ref. 23), 72 of which were among the 17,951 genes tested in the bulk cohort, were tested for enrichment among grade-associated genes by Fisher's exact test against all tested genes and against the other genes listed at the same association signals (Table S9, genes within 1 Mb with functional genomic data; 243 non-effector genes measured here), and by a Wilcoxon test of the absolute Wald statistic. Because several effector genes share one association signal, enrichment was also tested with the signal (lead variant) as the unit: the observed number of signals with at least one grade-associated effector gene was compared with its expectation given the genome-wide rate of grade-associated genes and the number of effector genes per signal, with an exact Poisson-binomial tail probability. Within signals, effector genes were compared with the other genes at the same signal by an exact Cochran-Mantel-Haenszel test stratified by signal (40 signals with both effector and non-effector genes). For display, the per-cluster change with grade of each grade-associated effector gene was taken from the per-cluster single-nucleus grade model.

#### Shape of the grade response

To test whether gene expression changed linearly with grade, the bulk cohort was divided into four grade groups (0-1, 1.5, 2-3 and 4; 23, 10, 8 and 51 knees) and a DESeq2 likelihood-ratio test compared a model with grade group as a factor against a nested model with the group's mean grade as a linear term (both with batch and the number of detected genes), with Benjamini-Hochberg correction across the 17,951 genes. For description, each grade-signature gene was assigned a response shape from variance-stabilized expression adjusted for batch and detected genes (limma removeBatchEffect with grade group protected): the difference between each grade group's mean and the grade 0-1 mean was expressed as a fraction of the grade 4 difference, and genes were classed as early (at least half of the change at grade 1.5), mid-plateau (at least three quarters by grade 2-3), graded (between), late (less than a quarter by grade 2-3) or overshoot (an intermediate group exceeding the grade 4 change by half; in Figure 8f the four rising overshoot genes were merged with the mid-plateau class, and Figure S2e shows them separately). The stability of these classes was estimated by 200 bootstrap resamples of knees within grade groups.

#### OARSI cartilage score

The OARSI cartilage histopathology grade (0-6) (24), referred to here as the OARSI score, of the medial weight-bearing region of each knee used for bulk RNA sequencing was taken from the laboratory histology record (see main Methods). Knee identity, side, macroscopic grade and score were cross-checked against the donor tissue inventory. One knee (OA donor 7) had no score, leaving 91 knees (Normal 17, Aged 16, OA 11, arthroplasty 47; Figure S5g). Agreement with macroscopic grade is reported as the Spearman correlation.

#### Differential expression with the OARSI score

The OARSI analysis used the genes, model and thresholds of the grade analysis, with the OARSI score in place of grade: DESeq2 with the design ~ Batch + Detected_Genes_z + OARSI_Score, the score entered as a continuous variable and Detected_Genes_z computed over all 92 knees. The 17,951 genes of the grade analysis were tested, the score coefficient was tested with the Wald test, p values were adjusted with the Benjamini-Hochberg method, and log2 fold changes per OARSI unit were shrunk with lfcShrink(type = "normal"). Genes were called differentially expressed at an adjusted p value < 0.05 (662 genes; 301 rising, 361 falling) (Figure S4b); for motif enrichment and the comparison with grade, a baseMean of at least 10 was also required (660 genes; 301 rising, 359 falling). The same code reproduced the primary grade model (the same 1,346 genes at adjusted p < 0.05 and baseMean ≥ 10; log2 fold changes within 0.001). Gene set enrichment (gseGO, GO biological process) and RcisTarget motif enrichment used the ranking, settings, databases and seed described for grade. In Figure S4c, rising processes are the ten non-redundant processes with the highest normalized enrichment scores among terms of at least 30 genes, as in Figure 1c, and falling processes were selected, at most two per biological program, from those with adjusted p < 0.05. The PCA of Figure 1a was used unchanged, and the association of PC1 with the OARSI score is the squared Pearson correlation over the 91 scored knees (Figure S4a).

#### Comparison of the OARSI and grade signatures

Genes were compared between the OARSI and grade models using genes with adjusted p < 0.05 and baseMean ≥ 10 in each model; directional agreement was assessed from the unshrunk estimates, and effect-size concordance as the Pearson and Spearman correlation of shrunk log2 fold changes over the 1,461 genes significant for either measure. GO processes were compared using terms at adjusted p < 0.05 in each analysis, and their concordance is the correlation of normalized enrichment scores over the 404 terms significant for either measure, 143 of them for both (Figure S5a-f).

#### OARSI score within arthroplasty knees

To test whether the OARSI score tracked the transcriptome within end-stage tissue, the same model was fitted to the 47 arthroplasty knees, all of macroscopic grade 4 (OARSI 1-6; 44 in batch 1 and 3 in batch 2), with genes called as above. The detection of each gene called was inspected as the number of knees with zero counts. For the 660 genes associated with the OARSI score in all knees, directional agreement was assessed from the Wald statistics of the two fits, and the effect-size ratio as the slope of a regression through the origin of the within-arthroplasty on the all-knee shrunk log2 fold changes (Figure S5h). As an independent test, the model was fitted to the 44 scored cadaveric knees alone; the genes associated with the score there (adjusted $p < 0.05$, baseMean ≥ 10; 11 rising, 270 falling) formed a signature that was scored in each arthroplasty knee as the mean z-scored variance-stabilized expression (DESeq2 vst, blind, on the 47 arthroplasty knees, with batch and detected genes removed by limma removeBatchEffect) of the rising genes minus that of the falling genes. The signature score was correlated with the OARSI score by Spearman correlation (Figure S5i), with a null distribution from 1,000 random gene sets of the same size drawn within baseMean deciles.

#### Single-nucleus RNA-seq processing and quality control

Filtered gene-expression count matrices were read per sample (for multiome samples, only the gene-expression modality was used for the analyses below). Multiome and GEM-X matrices were merged by gene symbol. Because the two Cell Ranger references (GRCh38-2020-A, GENCODE v32; GRCh38-2024-A, GENCODE v44) assign different symbols to 12,913 shared Ensembl genes (mostly unnamed long non-coding RNAs; 468 protein-coding), these genes were counted under different symbols in the two chemistries; no gene named in this study is affected. Integration features were restricted to symbols present in every sample, and chemistry was included as a covariate in the regression models that combined the two chemistries. Nuclei with 1,000-25,000 unique molecular identifiers (UMIs) and less than 5% mitochondrial reads were retained (137,919 nuclei; 478-11,456 per sample). No gene-count, complexity or computational doublet filter was applied (ambient RNA and doublets were assessed in a sensitivity analysis, below); contaminating and artifactual populations were instead removed at the cluster level as described below. Each sample was normalized with SCTransform (3,000 variable features), regressing the mitochondrial fraction; cell-cycle effects were not regressed.

#### Gene-span annotation and long-gene capture

Gene spans (annotated gene-body length) were computed from the Ensembl GRCh38 release 110 annotation for 61,267 genes. Nuclear RNA over-represents long genes, so a gene was defined as long when its span was at least eight times the median span of the annotated genes in the count matrix (13.8 kb), that is at least approximately 110 kb. Capture burden was defined, for each donor-cluster pseudobulk (for each donor in the neighborhood analysis), as the fraction of UMIs from genes with known span that fell in long genes lacking any extracellular matrix, collagen, cartilage, chondrocyte, skeletal, ossification, bone, connective tissue, proteoglycan, glycosaminoglycan, elastic fiber or matrix organization GO biological process annotation (4,149 of 4,379 long genes), so that the covariate does not absorb matrix biology.

#### Integration, clustering and removal of non-chondrocyte populations

Samples were integrated with Seurat (SCT normalization, 50 dimensions, canonical correlation analysis (CCA) anchors). To identify contaminating populations, nine lineage panels (osteoblast, osteoclast, adipocyte, endothelial, myeloid, lymphoid, mast cell, mural, nerve) and a ten-gene cartilage core panel were scored per nucleus, and a cluster was removed only when its highest contaminant score exceeded 0.10 and its cartilage score was below 0.05. Fibrotic and hypertrophic chondrocyte genes (*COL1A1*, *COL1A2*, *COL3A1*, *COL11A1*, *POSTN*, *SPARC*, *IBSP*, *RUNX2*, *SPP1*, *ALPL*) were deliberately excluded from the contaminant panels. This screen was applied in the re-integrated space (below).

Because variable features selected on the SCT residual variance were biased toward long genes, features were re-selected per sample by their residual variance above a loess fit of variance on gene span, combined into a consensus of 3,000 features, and all 137,919 quality-filtered nuclei re-integrated.

Integration was limited to the top 956 of these features by the sparse-matrix size limit of the integration step. Donor mixing in the new space was checked with an observed-to-expected same-donor nearest-neighbor ratio (k = 50), which was 1.43 compared with 1.44 in the original space. In the re-integrated space, the lineage panels were averaged per cluster of a resolution-1.0 clustering (42 clusters); three clusters (endothelial 3,714, myeloid 1,638 and adipocyte 637 nuclei) were removed, and one mixed fibrocartilage cluster that the panel rule retained was subclustered (resolution 0.2) and its osteoblastic (1,390), pericyte/smooth muscle (305) and mature osteoblast (155) subclusters were removed on marker expression. Clusters of the resolution-0.1 partition of the same space were also evaluated with marker-based admissibility criteria (a single top marker exceeding the next by a log2 fold change of at least 4, or of at least 1.5 when the top 20 markers had a median span at least eight times background; more than 50% of nuclei from one donor; or fewer than 50 nuclei; cluster stability was used as a diagnostic only), and inadmissible clusters together with one cluster defined by the long gene *PCDH9* were removed, leaving 108,600 chondrocyte nuclei. These were re-clustered from scratch (principal component analysis on the integrated assay, 50 components, Louvain clustering at resolution 0.1, seed 42), giving nine clusters (c0–c8) of 43,845, 20,075, 19,036, 10,084, 6,514, 3,672, 2,074, 1,959 and 1,341 nuclei. Between 26 and 30 of the 30 donors contributed nuclei to every cluster, and no donor contributed more than 12.2% of any cluster.

#### Cluster markers and identity

Markers were identified on the SCT assay after re-correcting the SCT counts to a common median UMI (Wilcoxon test, genes detected in at least 10% of nuclei, log2 fold change ≥ 0.25, at most 2,000 nuclei per cluster, Bonferroni-adjusted $p < 0.05$). For display, markers were required to be detected in at least 20% of nuclei of the cluster, at least 5 percentage points more often than elsewhere and to be shorter than the long-gene threshold, and were assigned to the cluster in which they were most specific; where fewer than three such genes existed, the best available genes were shown and marked as not specific. Pathway enrichment of markers used over-representation analysis (clusterProfiler) against Hallmark, GO biological process and Reactome gene sets (15-500 genes), with long genes and ribosomal protein genes removed symmetrically from the marker lists and from the universe of 17,420 tested genes (genes without a span annotation were also removed). Ribosomal protein genes were removed because they made up 22% of the c0 marker list (0 to 2.6% in the other clusters) and, when retained, displaced every c0 term with translation terms; the long-gene filter removed most terms that were significant only through long genes. Cluster-level gene set enrichment used a balanced contrast of each cluster's pooled pseudobulk (log2 CPM + 1, all donors combined) against the unweighted mean of the other eight, with ribosomal protein genes removed, ranked by fold change z-scored within gene-span deciles (fgsea; 15-500 genes). The clusters were compared with the published normal-donor populations of Swahn et al. (8) by one-sided hypergeometric tests of the overlap between each cluster's 100 top markers (Wilcoxon markers on the RNA assay, ranked by log2 fold change z-scored within gene-span deciles) and each population's published markers (adjusted $p < 0.05$, log2 fold change > 0.25), against the 40,813 genes detected in at least 10 nuclei; a match required at least five shared genes and adjusted $p < 0.05$ (Benjamini-Hochberg across cluster-population pairs) and was ranked by fold enrichment. Match stability was estimated by a 2,000-replicate bootstrap over marker genes.

#### Grade-associated gene expression

For each cluster, raw counts were summed per donor (pseudobulk); donor-clusters with at least 20 nuclei were retained, clusters were tested when at least 8 donors and three grade values were available, and genes with at least 10 counts in at least 50% of the retained donors were tested. DESeq2 was fitted with ~ technology + capture burden + macroscopic knee grade, grade as a continuous variable and capture burden centered within cluster (technology was dropped where it was not estimable), and the grade coefficient was tested with the Wald test with Benjamini-Hochberg correction (adjusted $p < 0.05$; DESeq2 default independent filtering). Log2 fold changes per grade unit are maximum-likelihood estimates. Pathway analysis of the grade response used the Wald statistic z-scored within span deciles, with cytosolic and mitochondrial ribosomal protein genes (and genes without a span annotation) removed, ranked in fgsea against Hallmark, GO biological process and Reactome (15-500 genes; positive normalized enrichment score indicates a pathway rising with grade); significant GO and Reactome terms were grouped into keyword-defined themes for display (Figure 3d).

#### Cluster abundance

Cluster abundance was analyzed per donor on the complete 30 × 9 donor-by-cluster grid of nuclei counts (true zeros retained). Donor proportions were centered log-ratio transformed (pseudocount 0.5 nuclei), and each cluster was tested by the Spearman correlation of its transformed share with continuous macroscopic knee grade and by a Kruskal-Wallis test across Normal, early OA and severe OA, with significance assessed against 10,000 donor-label permutations (two-sided, $p = (x + 1)/(N + 1)$, Benjamini-Hochberg across clusters). The analysis was repeated in the 24 multiome donors. To test for shifts within clusters, differential abundance was also tested in neighborhoods of transcriptionally similar nuclei with Milo (see below).

#### Identity checks of c5 and c8

To assess whether c5 reflected a dissociation response, a 19-gene immediate-early gene score was computed for all nuclei and its donor means were related to post-mortem interval and to each donor's c5 share. To assess the lineage of c8, detection of *COL2A1*, *ACAN* and *SOX9* was compared across clusters, curated chondrocyte, fibroblast, mural, smooth muscle, endothelial, immune and basement-membrane panels were scored, and c8 was tested against MSigDB C8 cell-type signatures with fgsea.

#### Milo neighborhood differential abundance

Differential abundance with grade was tested in neighborhoods of transcriptionally similar nuclei with miloR (28). The 108,600 chondrocyte nuclei were embedded in the 50 principal components used for clustering, a k-nearest-neighbor graph was built ($k = 30$), and neighborhoods were defined around a refined random sample of 5% of nuclei (graph-based refinement), giving 3,388 neighborhoods (median 174 nuclei). Nuclei were counted per donor in each neighborhood and modeled with a negative binomial generalized linear model (edgeR; trimmed mean of M values [TMM] normalization) with ~ technology + capture burden + macroscopic knee grade, grade continuous and capture burden computed per donor with the same function-neutral long-gene definition as the per-cluster grade model and centered. Multiple testing was controlled with the graph-overlap-weighted spatial false discovery rate (threshold 0.1). Each neighborhood was assigned to the cluster of most of its nuclei. As a sensitivity analysis, the principal components, graph and neighborhoods were rebuilt on the 96,175 nuclei of the 24 multiome donors (2,897 neighborhoods) and tested with ~ capture burden + grade. To relate abundance changes to disease state, each nucleus was scored with the bulk grade signature (mean log-normalized expression of the 593 grade-signature genes rising with grade minus that of the 622 falling, among genes detected in the single-nucleus data), averaged over the nuclei of each neighborhood, and correlated with the neighborhood log fold change (Spearman), overall and within clusters; the analysis was repeated after removing the 49 signature genes that also changed in the same direction in at least six clusters.

#### Healthy reference set and gene universe

Cluster identity was defined in the 12 donors with healthy cartilage (macroscopic knee grade ≤ 1; 51,443 nuclei), so that identity was not confounded with the OA response. Markers, pairwise contrasts, signature enrichment, classification and pseudobulk recall were repeated in all 30 donors as a check, and chromatin analyses in all 24 multiome donors. All 12 healthy donors were profiled by single-nucleus multiome, so chemistry could not be modeled within this set.

Where stated, analyses were restricted to protein-coding, non-long genes. These were genes of protein-coding biotype in the gene-span annotation with a genomic span below 8 × 13.8 kb (about 110 kb), the long-gene threshold used throughout; mitochondrial and ribosomal-protein genes were excluded.

Unless stated otherwise, pseudobulk profiles in this section were raw RNA counts summed per donor and cluster, and a donor-cluster entered only if it contained at least 20 nuclei. Per-nucleus log-normalized expression was log1p(counts per 10,000); for gene programs, program scores and AUROC, counts per 10,000 were computed over the protein-coding non-long genes analyzed.

#### Cluster-versus-rest and pairwise differential expression in healthy donors

Marker genes were tested on donor × cluster pseudobulk with limma-voom. Genes were filtered with filterByExpr (grouped by cluster, min.count = 10) and normalized by TMM, which left 12,215 genes in healthy donors and 13,662 in all donors. The design was ~0 + cluster + donor, so donor was a blocking factor; technology is nested in donor. Each cluster was contrasted with the unweighted mean of the other eight (weights +1 and −1/8). Moderation used empirical Bayes with robust = TRUE, and p values were adjusted by Benjamini-Hochberg (BH). A sensitivity model added the mean-centered capture-burden covariate. This model was not used as the primary model, because burden differs between clusters in healthy donors and partly absorbs identity. Markers were genes with adjusted $p < 0.05$ and log2 fold change > 0.5.

Pairwise distinctness was assessed in the same donor-blocked model (without the burden term) using all 36 pairwise contrasts. For each pair we counted the non-long genes (annotated span below the long-gene threshold) with adjusted $p < 0.05$ and |log2 fold change| > 1 (Figure S9f).

#### Zonal and hypertrophy axes and published signatures

Zonal gene sets were taken from microdissected human knee cartilage (27): superficial-up, 60 genes (58 detected); superficial-down (middle/deep-enriched), 28 genes (27 detected); and deep-up, 19 genes.

To place the clusters on the two axes (Figure 4a), each gene was z-scored across all healthy nuclei on log-normalized expression, and genes with zero variance were dropped. For each nucleus, the zonal score was the mean z of the superficial-up genes minus the average of the mean z of the superficial-down and deep-up genes (positive = superficial). The hypertrophy score was the mean z of *RUNX2*, *COL10A1* and *IBSP*; *RUNX2* and *IBSP* are also members of the deep-up set.

Scores were averaged per donor-cluster (at least 20 nuclei). Clusters are shown as the mean ± standard error of donor means. Consistency of the order across donors was summarized as the median Spearman correlation between each donor's cluster order and the order of the averaged donor means. The c2 > c3 and c3 > other-cluster hypertrophy contrasts were tested with paired Wilcoxon tests across donors.

Enrichment of zonal and published signatures (Figure S9a-b) used gene set enrichment analysis (fgseaMultilevel, eps = 0, minSize = 2, maxSize = 500). Genes were ranked by each cluster's moderated t from the donor-blocked cluster-versus-rest model, and p values were BH-adjusted within cluster and signature family (zonal or published). The published signatures were the marker genes named in the text for five populations by Ji et al. (6); the marker genes named in the text for eleven populations by Fan et al. (9); and, from Swahn et al. (8), the top 50 genes per normal-donor population by average log2 fold change, among markers with adjusted $p < 0.05$ and log2 fold change > 0.

#### Marker expression and single-gene specificity

For the genes used to name the clusters (Figure 4b), the percentage of nuclei with non-zero counts and the mean log-normalized expression were computed per cluster. The means were z-scored across clusters for display.

Single-gene specificity (Figure S9e) was the area under the receiver operating characteristic curve (AUROC) for each cluster against all other healthy nuclei. AUROCs were computed with presto on log-normalized expression of protein-coding non-long genes detected in at least 1% of healthy nuclei. The gene with the highest AUROC among genes with positive log fold change is shown.

#### Recurrent gene programs

Gene programs were identified separately in each healthy donor, following the meta-program approach of Gavish et al. (63) (Figure 4c). For each donor, up to 4,000 nuclei were sampled (seed 1), and genes were restricted to protein-coding non-long genes detected in at least 1% of the donor's nuclei. The 2,000 most variable genes were selected (Seurat vst), and counts per 10,000 were scaled to unit standard deviation without centering. Non-negative matrix factorization (RcppML; seed 1, tol = $10^{-5}$, maxit = 300) was run at K = 4-9, and each program was represented by its 50 highest-loading genes.

A program was retained as robust if it shared at least 35 of 50 genes with a program at another K in the same donor, and at least 10 of 50 with a program from another donor; 377 of 468 programs were robust. Within each donor, programs were ranked by cross-donor support, and a program sharing more than 10 genes with a higher-ranked program was removed, leaving 65 programs.

The 65 programs were clustered by average-linkage hierarchical clustering on 1 − (shared genes/50), cut at height 0.85. Clusters supported by at least four donors were retained, which gave six meta-programs supported by 5-12 donors. Each meta-program's genes were the up-to-50 genes present in at least 25% of its member programs, ranked by frequency. Meta-programs were named from their top genes.

Per nucleus, the meta-program score was the mean log1p(counts per 10,000) of the meta-program genes minus that of control genes. Controls were 5 per gene, drawn from the same of 25 expression bins (seed 1). Scores were z-scored within donor over all of that donor's nuclei, averaged per donor-cluster (at least 20 nuclei), and then averaged over healthy donors.

#### Separability and stability

Pseudobulk recall (Figure 4d) was tested by leave-one-donor-out assignment of pseudobulk profiles. Profiles were log2(counts per million + 1), and each profile was centered on its donor's mean across clusters; only donors contributing at least five clusters were used. Each test-donor profile was assigned to the cluster whose centroid of training-donor profiles had the highest Pearson correlation. Two feature sets were used. The first was the 2,000 most variable protein-coding non-long genes, selected from the residual of a loess fit (span 0.3) of log variance on log mean among 9,013 eligible genes detected in at least 1% of healthy nuclei. The second was the 956 integration features used for clustering.

Reclustering stability was measured on all 108,600 nuclei. The 50-component PCA of the integrated assay was recomputed (seed 42); reclustering on it at resolution 0.1 reproduced the original partition (adjusted Rand index 1). Ten independent subsamples of 80% of nuclei were drawn (seed 2026), and each subsample was reclustered at resolution 0.1 (shared nearest-neighbor graph, k = 20; Louvain; seed 42). Each original cluster's stability was the median, over subsamples, of its best-match Jaccard index. On all nuclei, resolutions 0.05, 0.15 and 0.2 gave 3, 17 and 20 clusters; subsamples were also reclustered at 0.15 and 0.2.

Single-nucleus classification (Figure S9d) used a multinomial elastic net (glmnet, $\alpha = 0.5$) with log-normalized expression of the same 2,000 genes as input. Training data were capped at 1,500 nuclei per cluster (seed 7). The penalty was chosen once, as $\lambda$ at one standard error from 4-fold donor-grouped cross-validation on the first training fold ($\lambda = 0.052$; misclassification loss; 30 $\lambda$ values; minimum ratio $10^{-3}$), and held fixed. Testing was leave-one-donor-out across the 12 healthy donors. Performance is reported as balanced accuracy (mean per-cluster recall) against a chance level of 1/9.

**Chromatin accessibility in the single-nucleus multiome**

The multiome nuclei carried the snRNA-seq cluster labels by barcode: 81,670 labeled nuclei from 24 donors, including the 12 healthy donors. Peaks were the unified set of 247,013 regions (see Single-nucleus multiome processing).

Peak counts were summed per donor × cluster. Columns with at least 20,000 counts in peaks were kept (208 of 213; 105 in healthy donors).

Cluster-specific regions (Figure 4e) were tested with limma-voom. filterByExpr (min.count = 5, min.total.count = 15) left 57,739 peaks in healthy donors. Normalization was TMM, with voomWithQualityWeights. The design was ~0 + cluster + donor, contrasting each cluster with the unweighted mean of the other eight. Regions with BH-adjusted $p < 0.05$ and log fold change > 0 were counted.

Motif enrichment among the c2-specific regions used JASPAR2020 CORE vertebrate (human, taxon 9606) motifs (633 matrices), which were matched to peaks with Signac AddMotifs on BSgenome.Hsapiens.UCSC.hg38 at default settings. Each motif was tested against all tested peaks with a Cochran-Mantel-Haenszel test stratified by deciles of peak GC content. This gave a Mantel-Haenszel odds ratio and a one-sided p value, BH-adjusted within cluster. Clusters with fewer than 10 specific regions were not tested. For display, the best motif per family was shown.

For gene-body accessibility (Figure 4g), gene bodies were Ensembl (EnsDb.Hsapiens.v86) protein-coding and lincRNA genes extended by 2 kb on each side. Gene-body counts were the summed counts of overlapping peaks and were tested in the same donor-blocked design (filterByExpr min.count = 10). Concordance of c2 marker genes (for this analysis, adjusted $p < 0.05$ and log2 fold change > 1 in the healthy-donor marker model) with chromatin was tested with a one-sided Fisher's exact test against non-marker genes. A marker was concordant if its gene body was more accessible at $p < 0.05$ with log fold change > 0.

Positions between c2 and the other clusters (Figure 4f) were computed for each cluster k other than c2, using donor-adjusted cluster coefficients from the healthy fits. Each feature's position was (coefficient_k − reference)/(coefficient_c2 − reference), where the reference was the mean coefficient of the clusters other than c2 and k. Features with |coefficient_c2 − reference| ≤ 0.1 were excluded. The median over features is reported, with a percentile 95% confidence interval from 1,000 bootstrap resamples of features (seed 7). Three feature sets were used: the 539 peaks more accessible in c2 at nominal $p < 0.001$; the RNA of the 98 c2 marker genes (adjusted $p < 0.05$, log2 fold change > 1) detected in the multiome nuclei (multiome RNA pseudobulk in the same limma-voom design); and the gene-body accessibility of the same marker genes.

Motif accessibility (Figure S9c) was estimated with chromVAR on the donor × cluster peak pseudobulk, using the same 633 motifs, GC bias from peak GC content and 50 background iterations (seed 7). Deviation z-scores were computed over the 208 retained donor × cluster columns of all 24 donors. Healthy-donor columns were then tested in limma (~0 + cluster + donor; each cluster against the mean of the others; empirical Bayes; BH-adjusted across motifs within contrast).

### Shared grade response and Hallmark pathways

The shared response (Figure S10) was taken from the per-cluster pseudobulk DESeq2 grade model and its GSEA described under Grade-associated gene expression; no new model was fitted. A gene was shared if it had adjusted $p < 0.05$ in all nine clusters. All 17 such genes had the same sign in every cluster. The before-covariate counts (Figure S10a) come from the same model without the capture-burden term (~ technology + grade).

Hallmark terms (Figure S10d) are those with adjusted $p < 0.05$ in at least one cluster in the grade GSEA.

The effect of the covariate definition on pathways (Figure S10e) was assessed as the change in normalized enrichment score between the function-neutral and all-long-gene burden models. Change for matrix-themed terms (term-name regular expression) was compared with that for all other terms by a Wilcoxon rank-sum test.

### Ambient RNA and doublet sensitivity analysis

Ambient RNA and doublets were assessed as sensitivity analyses on all 137,919 quality-filtered nuclei, before cluster-level removal, with cluster labels from the integrated object so that non-chondrocyte populations informed the contamination profile. SoupX (1.6.2) estimated the ambient profile from each sample's raw droplet matrix (gene-expression modality only) and the contamination fraction with autoEstCont (median 0.045, range 0.010-0.358); for the two samples in which autoEstCont failed (estimate above 0.5), the median of the other samples was used, and counts were corrected with adjustCounts with integer rounding. DecontX (decontX 1.0.0) was run per sample on the filtered matrices with labels that distinguished chondrocytes from each non-chondrocyte lineage removed during cleaning (osteoblast, endothelial, mural (pericyte), myeloid and adipocyte), and decontaminated counts were rounded. A DecontX run with the nine chondrocyte clusters as separate labels over-corrected, assigning the highest contamination to the largest cluster (c0) and removing *PRG4* from c1 to c8, and is shown for comparison only. Doublets were predicted per sample with scDblFinder (random mode; expected rate 0.8% per 1,000 nuclei for multiome and 0.4% per 1,000 nuclei for GEM-X libraries). The per-cluster continuous-grade pseudobulk model was refitted unchanged on each corrected count set (capture burden recomputed from the corrected counts), with and without predicted doublets removed, and compared with the original model by retention of grade-associated genes, sign concordance and correlation of log2 fold changes; for the 17 shared genes, the number of clusters in which each remained significant was counted (Figure S11). Contamination and doublet fractions were related to grade by Spearman correlation across donors.

### Three-stage analysis

Donors were staged as Normal, grade ≤ 1 (12 donors); early OA, grade 1.5-2 (4 donors); and severe OA, grade 3-4 (14 donors) (Figure S12).

Per-cluster pseudobulk DESeq2 was fitted with the inclusion and gene filters of the continuous model, except that at least 10 donor-clusters were required. A cluster was skipped if Normal or severe OA had fewer than three donors. The design was ~ technology + burden + stage, and technology was estimable in all clusters. Early OA versus Normal and severe OA versus Normal were tested with Wald tests (BH, adjusted $p < 0.05$). Any stage effect was tested with a likelihood-ratio test against ~ technology + burden. The early-OA contrast was fitted when at least two early-OA donors contributed but reported only with at least three. It was therefore not reported for c6 and c8. Severe OA was also tested against early OA (Wald test on the same fit) in the seven clusters with at least three early-OA donors (c0 to c5, c7).

Severe-OA genes were classified by their early-OA effect. Genes were early onset if the effects had the same sign and |early log2 fold change|/|severe log2 fold change| ≥ 0.5, progressive if they had the same sign and a ratio < 0.5, and late only if the early-OA change had the opposite sign. Severe-OA genes were further classed as established early (early OA against Normal adjusted $p < 0.05$ in the same direction and no significant change from early to severe OA) or changing with progression (severe against early OA adjusted $p < 0.05$ in the same direction); the remainder were unresolved.

Agreement with the continuous model was the Spearman correlation of severe-OA versus Normal Wald statistics with the continuous-grade Wald statistics.

GO biological process enrichment per contrast used clusterProfiler gseGO, with ranking by −log10(p) × sign(log2 fold change); minGSSize = 10, maxGSSize = 500, BH, seed 42. Over-representation of GO biological process (clusterProfiler enrichGO with org.Hs.eg.db, simplified at a semantic similarity of 0.7), MSigDB Hallmark and NABA gene sets was tested for each class, separately for all, rising and falling genes, pooled over the seven clusters (a gene counted when it was in the class in at least two clusters, or in at least one where stated) against all genes tested in at least one cluster, and within c0 against genes tested in c0 (gene sets of 10-500 genes, NABA 10-2,000; lists of fewer than 10 genes not tested; BH). Class shares were recomputed without long genes, and the enrichments with genes and background restricted to non-long protein-coding genes; for this check, GO terms were read from the unsimplified results.

Cluster abundance by stage was tested as described under Cluster abundance.

#### Cluster-distinctive grade responses

To test whether grade responses differed between clusters (Figure 5), all clusters were analyzed jointly. Donor-clusters with at least 20 nuclei were included (239 from 30 donors). Genes were kept if they had at least 10 counts in at least 50% of all 239 donor-clusters, giving 9,051 genes. Counts were size-factor normalized and variance-stabilized (DESeq2, blind = TRUE) across all donor-clusters together.

For each gene, two linear mixed models were fitted by maximum likelihood (lme4). The full model was technology + burden + cluster + cluster × grade + (1 | donor), and the reduced model was the same with a single common grade slope. Grade was continuous, and the burden covariate (the ECM-excluded capture burden of the per-cluster grade model) was mean-centered within cluster. The cluster × grade interaction was tested by likelihood-ratio test (8 df), BH-adjusted across genes.

The deviation of each cluster from the other eight was taken from the full model as its grade slope minus the unweighted mean slope of the other eight clusters (contrast weights +1 and −1/8). The standard error came from the fixed-effect covariance matrix, giving a Wald z. A two-sided normal p value was BH-adjusted across genes within cluster.

A grade association was called cluster-distinctive when all of the following held: it was significant in that cluster in the per-cluster DESeq2 grade model (adjusted $p < 0.05$); the interaction had adjusted $p < 0.05$; the deviation had adjusted $p < 0.05$ with the same sign as the cluster's DESeq2 log2 fold change; and the cluster's own mixed-model slope also had that sign.

A gene was called opposite in direction if it was distinctive, the median log2 fold change of the other clusters had the opposite sign, and at least one other cluster was significant in the opposite direction. A pathway was called opposite if it was significant with positive normalized enrichment score in one cluster and negative in another in the per-cluster grade GSEA.

Cluster-distinctive pathways were identified by GSEA (fgsea, minSize = 15, maxSize = 500; seed 42) on each cluster's deviation z across the 9,051 genes, using the Hallmark, GO biological process and Reactome collections (MSigDB via msigdbr). P values were BH-adjusted within cluster and collection. A pathway was distinctive if its deviation had adjusted $p < 0.05$ and it was also significant (adjusted $p < 0.05$) with the same sign in that cluster's grade GSEA.

#### Display of nuclei by grade

Figure S8a introduces no statistical analysis. The uniform manifold approximation and projection (UMAP) embedding of Figure 3a is split by grade, with all nuclei drawn in gray behind each panel and that grade's nuclei colored by cluster; point size and opacity were scaled per panel by nucleus count (size = $14/\sqrt{n}$ bounded to 0.20-0.55; opacity = $25/\sqrt{n}$ bounded to 0.78-0.95). Grades contributed by a single donor are marked.

#### Matrisome annotation and curated matrix families

Matrisome genes and their categories were taken from the Naba matrisome (10) as distributed in MSigDB (msigdbr 7.5.1; gene sets NABA_COLLAGENS, NABA_PROTEOGLYCANS, NABA_ECM_GLYCOPROTEINS, NABA_ECM_REGULATORS, NABA_ECM_AFFILIATED and NABA_SECRETED_FACTORS; 1,026 genes, none assigned to more than one category), of which 1,023 were present in the single-nucleus count matrix. Because the six Naba categories group functionally unrelated genes (for example, crosslinking enzymes, proteases and protease inhibitors are all ECM regulators), genes were also assigned to 24 function-defined families curated from explicit gene lists rather than symbol patterns: fibrillar, network/basement membrane, FACIT and other collagens; hyalectan, small leucine-rich (SLRP) and basement membrane/cell-surface proteoglycans; cartilage and adhesive glycoproteins; elastic fiber; collagen crosslinking and hydroxylation; procollagen processing; matrix metalloproteinases; ADAMTS proteoglycanases; cathepsins; TIMPs; serpins and other inhibitors; glycosaminoglycan linker, chondroitin/dermatan sulfate, heparan sulfate and keratan sulfate synthesis; hyaluronan; ECM receptors; and matrix-bound growth factors (357 family-gene assignments, 356 genes; *SDC1* is in two families; 268 of the 356 genes are in the Naba matrisome). The union of the two annotations (1,023 Naba genes plus 88 family genes outside the Naba matrisome; 1,111 genes) defined the matrisome universe for gene-level analyses. Long genes were defined as in the capture-burden analysis (span at least 8 times the 13.8 kb background); 176 of the 1,023 Naba genes were long.

#### Matrisome composition of each cluster

Analyses used the donor-by-cluster pseudobulk counts of the grade model, keeping donor-clusters with at least 20 nuclei (239 donor-clusters from 30 donors; 102 from the 12 healthy donors, grade 0-1). Counts were converted to counts per million (CPM) of the column's total counts. The matrisome fraction of the transcriptome was the summed CPM of Naba genes divided by $10^6$, and the share of each category or family was its summed CPM as a percentage of the summed CPM of all genes in that annotation, computed per donor-cluster and averaged over donors. All composition results shown exclude long Naba genes; *FN1* (not a long gene) was shown as a separate slice. Disease stages were Normal (grade 0-1; 12 donors), early OA (grade 1.5-2; 4 donors) and severe OA (grade 3-4; 14 donors). For family-level change with *FN1* removed, family shares were computed over family members that were non-long Naba genes other than *FN1*, so that families made up largely of non-Naba genes (glycosaminoglycan synthesis, ECM receptors) were not scored or were represented by one gene; baseline family shares in healthy donors used all curated family genes except long Naba genes (27 long family genes outside the Naba matrisome, mainly ECM receptors and glycosaminoglycan-synthesis enzymes, were retained).

Change with grade of each share (and of the total matrisome fraction) was estimated per cluster with a donor-level linear model, share ~ technology + capture burden + grade, with grade continuous, capture burden (the function-neutral covariate of the grade model) centered within cluster, technology dropped where only one level was present and no fit with fewer than 8 donor-clusters. The grade coefficient (percentage points per grade unit) was tested with a t test and p values were adjusted by the Benjamini-Hochberg method across all cluster-by-category, cluster-by-family or cluster-by-total tests of a given analysis.

#### Matrix gene repertoire and specificity

Mean CPM per cluster over healthy donor-clusters was computed for each Naba gene. Cluster specificity in healthy donors was tested with limma-voom: genes were filtered with filterByExpr (grouped by cluster) and TMM-normalized, and for each cluster a model ~ donor + in-cluster (donor as a fixed blocking factor) contrasted that cluster against the other eight clusters pooled; p values were Benjamini-Hochberg adjusted over all filtered genes. A gene was cluster-specific at adjusted $p < 0.05$ and log2 fold change > 0.5. The displayed repertoire comprised 47 genes chosen to represent fibrillar and other collagens, proteoglycans, glycoproteins, mineralization, crosslinking and proteases/inhibitors; expression was shown as log2(mean CPM + 1) z-scored across clusters.

#### Grade response of matrisome genes in each cluster

Per-gene grade effects were the log2 fold changes per grade unit of the per-cluster pseudobulk DESeq2 model described above (~ technology + capture burden + grade; genes with at least 10 counts in at least 50% of a cluster's donors). Because this filter excludes genes expressed in few donors, matrisome genes that failed it in a cluster but had at least 10 counts in at least three donors were refitted in that cluster with the same donor-clusters, design and size factors (estimated on the genes passing the original filter), together with all originally tested genes so that the dispersion trend was anchored as in the original fit. Benjamini-Hochberg adjustment was applied within the newly admitted matrisome genes of each cluster, and a refitted gene was called changed only if adjusted $p < 0.05$, the Spearman correlation of donor CPM with grade had $p < 0.05$ and the same sign, and the gene was detected (at least one count) in at least three donors. Log2 fold changes of originally tested genes in the refit agreed with the original model (Pearson $r \geq 0.99992$ in every cluster; median absolute difference $\leq 7.7 \times 10^{-4}$). This gave 3,107 original and 1,462 refitted matrisome gene-by-cluster tests (484 and 50 significant).

Shared and cluster-specific responses were classified with the cluster-distinctive criteria applied to all genes (significant in the cluster, cluster-by-grade interaction and deviation of the cluster's slope from the unweighted mean of the other eight clusters at adjusted $p < 0.05$, deviation and the cluster's own mixed-model slope of the same sign as its DESeq2 log2 fold change); 346 matrisome genes entered the interaction model. In addition, for genes that changed in exactly one cluster, each of the other eight clusters was classed as flat (not significant, 95% confidence interval of the log2 fold change including 0 and absolute log2 fold change < 0.2), not expressed (not testable even with the relaxed filter and mean CPM at most one third of that in the changing cluster) or inconclusive. Genes were labeled restricted to the changing cluster when all eight others were not expressed, and as changing only there when all eight others were flat or not expressed. The shared group displayed comprised the six genes significant in the most clusters (at least four) that met neither criterion.

To identify matrix genes confined to one cluster irrespective of testability, the expression pattern of each matrisome gene was evaluated over all donors and within each stage from mean CPM per cluster: exclusive, at least 1 CPM in the top cluster and < 0.2 CPM in every other; enriched, at least 1 CPM and at least fourfold above the next cluster (next-cluster mean floored at 0.01). A pattern was accepted only if at least three donors had at least three counts in the top cluster.

#### Matrisome genes expressed per cluster

A matrisome gene was considered expressed in a cluster when its mean CPM over donors was at least 1 and at least three donors had at least three counts, or when it was testable in that cluster by the grade model or the relaxed refit. Expressed genes were classified by their grade call in that cluster (up, down, not changed, or too sparse to test). Because c0 contributed the deepest pseudobulks (median 7,087,538 counts per donor-cluster against 816,784 for c1-c8), the expression rule was repeated after binomial thinning of every donor-cluster to at most the median library size of the c1-c8 donor-clusters (random seed 20261009).

#### Length-matched matrix family scores and heterogeneity

Family-level grade responses were scored on the Wald statistics of the per-cluster grade model. Within each cluster, tested genes with a span annotation were divided into span deciles and the Wald statistic was z-scored within decile. A family's score was the mean within-decile z of its tested members, with a standard error from the finite-population variance of drawing that number of genes from each decile without replacement; the score was tested against zero with a two-sided normal test, required at least six tested family genes, and was Benjamini-Hochberg adjusted across all family-by-cluster scores. Whether a family's response differed between clusters was tested with a Friedman test on within-decile z (genes as blocks, the nine clusters as treatments), using family genes tested in all nine clusters and requiring at least six such genes, with Benjamini-Hochberg adjustment across families. The same scores and tests were computed on the severe OA versus Normal and early OA versus Normal contrasts of the three-stage per-cluster model and on a per-cluster Normal (11 donors) versus arthroplasty (12 donors) DESeq2 model with the same covariates.

#### Matrix gene enrichment and cluster-by-grade interaction

Enrichment of curated family genes among each cluster's grade-associated genes (adjusted $p < 0.05$) was tested with Fisher's exact test against all genes with an adjusted p value in that cluster, with Benjamini-Hochberg adjustment across clusters; a cluster was marked significant when the adjusted p was < 0.05 and the lower 95% confidence limit of the odds ratio exceeded 1. For the gene-level heatmap of curated family genes estimated in all nine clusters (103 genes), the cluster-by-grade interaction was tested per gene on variance-stabilized pseudobulk expression (DESeq2 varianceStabilizingTransformation, blind, on genes with at least 10 counts in at least 50% of all donor-clusters) with lme4 linear mixed models fitted by maximum likelihood, expression ~ technology + capture burden + cluster × grade + (1 | donor) against the model without the interaction (likelihood-ratio test), with Benjamini-Hochberg adjustment over the 346 matrisome and family genes tested.

#### Cell-cell communication inference by stage

Communication between the nine chondrocyte clusters was inferred with CellChat (2.1.2) (32,33) separately at each disease stage, using the 108,600 chondrocyte nuclei of the final snRNA-seq object. Donors were assigned to stages by macroscopic knee grade: Normal, grade 0-1 (12 donors, 51,443 nuclei); early OA, grade 1.5-2 (4 donors, 10,636 nuclei); and severe OA, grade 3-4 (14 donors, 46,521 nuclei). A cluster entered a stage's analysis only if it contained at least 25 nuclei at that stage. All nine clusters met this threshold at every stage (smallest population: c6 *SLC14A1*+ at early OA, 113 nuclei). For each stage, the log-normalized RNA expression matrix (layers joined) and the cluster labels were used to build a CellChat object. The complete human CellChatDB distributed with CellChat v2 was used without curation or added interactions: 3,234 interactions, of which 1,281 were secreted signaling, 424 ECM-receptor, 535 cell-cell contact and 994 non-protein signaling. Signaling genes were subset with subsetData. Overexpressed genes and interactions were identified with identifyOverExpressedGenes and identifyOverExpressedInteractions using default settings (Wilcoxon test, $p < 0.05$, positive markers only, no fold-change or percentage cut-off, minimum 10 cells per group), which retained 1,885, 1,798 and 2,163 ligand-receptor pairs at Normal, early OA and severe OA. No smoothing or projection onto a protein-protein interaction network was applied. Communication probabilities were computed with computeCommunProb using the truncated mean (type = "truncatedMean", trim = 0.1) as the average expression per cluster. Other settings were defaults (Hill coefficient Kh = 0.5, n = 1; raw.use = TRUE;

population.size = FALSE, so probabilities were not weighted by cluster size). Significance was assessed with 100 permutations (nboot = 100, seed 42). We used the truncated mean instead of CellChat's default trimean because the trimean is zero unless a gene is detected in roughly 25% of a cluster's nuclei, a single-cell threshold that discards lower-expressed receptors in nuclear data. A parallel trimean run was performed but was not used for any figure. Links supported by fewer than 10 nuclei in either cluster were removed (filterCommunication, min.cells = 10). Pathway-level probabilities (computeCommunProbPathway, $p < 0.05$), aggregated networks (aggregateNet) and network centrality (netAnalysis_computeCentrality, pathway level) were then computed. Significant links ($p < 0.05$) were extracted at ligand-receptor resolution and at pathway level with subsetCommunication.

#### Network summaries, pathway comparisons and receptor composition

At each stage, network size was summarized as the number of significant ligand-receptor links (one link is a ligand-receptor pair between a sender and a receiver cluster) and as the sum of their communication probabilities (Figure 7a). For the *FN1*, FGF and TENASCIN pathways, the probabilities of significant links were summed per stage (Figure 7b). Each ligand's share of FGF pathway probability (Figure 7c) and each ligand-receptor pair's share of *FN1* probability (Figure 7d) were expressed as percentages of the stage total. To compare pathways between stages, the three stage objects were merged (mergeCellChat) and rankNet was run in comparison mode (stacked = FALSE, do.stat = TRUE) for each pair of stages. In this test, each pathway's communication probabilities were compared between the two stages over all sender-receiver cluster pairs by a paired Wilcoxon signed-rank test (entries with $p > 0.05$ set to zero). The Normal versus severe OA p values are reported. Because nuclei were pooled within each stage and the test units are sender-receiver cluster pairs, these p values do not reflect donor-to-donor variation, and they were not corrected across pathways. For the three *FN1* receptor complexes whose share changed, the percentage of the Normal-to-severe OA change already present at early OA was computed as $100 \times (\text{early} - \text{Normal})/(\text{severe} - \text{Normal})$ of the receptor share (Figure 7e). For comparison, the same quantity was calculated for *FN1* mRNA as 100 × the ratio of the median early-OA log2 fold change to the median severe-OA log2 fold change against Normal. These fold changes came from the per-cluster pseudobulk stage model (DESeq2, ~ technology + capture burden + stage, Wald tests against Normal; see the differential expression section). Early-OA estimates were used for the seven clusters with at least three early-OA donors (c0 to c5, c7) and severe-OA estimates for all nine clusters. The breadth of sending and receiving for each pathway (Figure 7f) was the effective number of clusters, defined as $\exp(-\Sigma\, p_i \ln p_i)$, where $p_i$ is cluster i's fraction of the pathway's summed probability as source or target (maximum 9). Autocrine links counted toward both roles. Each cluster's share of *FN1* probability received was calculated per stage (Figure 7g). The ligand-receptor composition of the three pathways, the probability sent and received by each cluster, and the per-cluster probabilities of the three *FN1* receptor complexes (Figure S20) were obtained by summing significant-link probabilities over the relevant clusters.

#### Donor-level communication analysis

To obtain donor-level statistics, CellChat was run separately for each of the 30 donors with the settings described above (truncated mean, trim = 0.1; population.size = FALSE; full CellChatDB.human; 100 permutations; filterCommunication, min.cells = 10). A cluster entered a donor's analysis if it had at least 25 nuclei in that donor; all 30 donors were analyzed and 19 retained eight or nine clusters. For each donor, the three FN1 receptor shares, the *FGF1* share of FGF probability, the summed FN1, FGF and TENASCIN probabilities, c0's share of FN1 probability received and the effective number of FN1-receiving clusters were computed from significant links as for the stage networks; the *FGF1* share was undefined in two donors without significant FGF links. As a check independent of CellChat's permutation filter, a donor-level expression product (ligand truncated mean in the sender cluster multiplied by the geometric mean of the receptor subunits in the receiver cluster, summed over cluster pairs) was computed for the same ligand-receptor pairs. With the donor as the unit, each measure was correlated with grade (Spearman) and compared between Normal (12) and severe OA (14) donors (Wilcoxon rank-sum test), with Benjamini-Hochberg correction within each set of measures; the three FN1 receptor shares were the confirmatory tests. Early OA (4 donors) was compared with Normal and with severe OA without correction (Wilcoxon rank-sum, nominal). Sensitivity analyses repeated the tests in multiome donors only, in a rank regression on grade adjusting for technology, and in donors with at least eight clusters and 1,000 nuclei. Stage means were given 95% confidence intervals from 5,000 bootstrap resamples of donors within stage (Figure S21).

#### Network visualization

Circle plots (netVisual_circle) were drawn with the nine clusters in fixed positions. Node size was the percentage of that stage's nuclei in the cluster, on one maximum shared by all panels (43.9%). Aggregate networks (Figure S15, rows 1-2) showed the number of significant links and the summed probability (aggregateNet) for each cluster pair, with one edge-width maximum shared across stages in each row (153 links; 1.454). Differential networks (Figure S15, rows 3-4) were drawn with netVisual_diffInteraction for the Normal to early OA, early to severe OA and Normal to severe OA transitions, separately for link count and for probability. Only the 20% of edges with the largest absolute change were shown (top = 0.2). The changes for all 243 cluster-pair × transition combinations were also tabulated. Pathway networks (Figure S17, rows 1-3) used the pathway-level probability arrays after masking entries with $p > 0.05$, with a shared maximum within each pathway (*FN1*, 0.281; FGF, 0.141; TENASCIN, 0.0237). The *FN1-ITGA5*/*ITGB1*, *FN1-CD44* and *FN1-ITGAV*/*ITGB8* networks (Figure S17, rows 4-6) used the significance-masked ligand-receptor probability arrays on a single maximum shared by all nine panels (0.132).

#### MatriCom corroboration

As an independent, threshold-based method, MatriCom (1.0; GitHub izzilab/MatriCom, commit ec8fe0f) (34) was run separately at each stage on the same nuclei, clusters, stages and 25-nucleus population floor. The package's computational functions were evaluated directly from its Shiny application source, so the algorithm was identical to the web application's. Its bundled databases were used: 28,391 interactions, a 4,032-gene matrisome list and a 25-entry multimer exclusion list. Parameters were the application defaults. A gene was considered expressed by a population if at least 30% of its nuclei had non-zero expression and its mean non-zero log-normalized expression was at least 1. Further defaults were homomeric interactions removed, the multimer exclusion list applied, the model maximized (reciprocal duplicates removed), reliability score 3 only, homocellular and heterocellular communication, and the Matrisome, Surfaceome and Extracellular (non-matrisome) compartments. As a permissive sensitivity analysis, the analysis was repeated with a 10% positive-population threshold (Figure S16a). Each communication pair (sender cluster, gene 1, gene 2, receiver cluster) was classified by its presence across the three stages. Pairs gained or lost between Normal and early OA, and between Normal and severe OA, were attributed to the gene whose percentage of positive nuclei changed more. A pair was called high-confidence when that change was at least 10 percentage points and the gene changed significantly (adjusted $p < 0.05$) in the same direction in that cluster in the matched stage contrast of the pseudobulk stage model. For *FN1*-containing high-confidence pairs gained between Normal and severe OA, the gating partner was defined as the gene that crossed the 30% threshold (below 30% in Normal, at least 30% at severe OA) (Figure S16b). For all gained high-confidence pairs, attribution to the gene with the largest change was compared with attribution to the gene that crossed the threshold, for genes credited with at least 20 pairs under either rule (Figure S16c). For Figure 7h, the percentage of nuclei with non-zero expression of *FN1* and of *ITGA5*, *ITGB1*, *CD44*, *ITGAV*, *ITGB8*, *ITGB5*, *ITGA11*, *SDC4* and *LOXL3* was computed per cluster and stage. Because the two methods use different databases, counting units and inclusion rules, only the direction of change between stages was compared between CellChat and MatriCom.

#### FGF ligand balance and ERK feedback output

FGF ligands, receptors and ERK feedback genes were taken from the continuous-grade pseudobulk model (log2 fold change per grade unit; see the differential expression section) (Figure S18a). For donor-level measures (Figure S18b-c), raw counts were summed per donor and cluster. Donor-cluster pseudobulks with at least 20 nuclei were kept, and clusters with at least 8 such donors were analyzed (19 to 30 donors per cluster). Within each cluster, genes with at least 10 counts in at least 50% of donors were retained, and *FGF1*, *FGF2*, *FGFR1* and *FGFR3* were always retained. Counts were then variance-stabilized (DESeq2 vst, blind = TRUE, design ~ 1). The *FGF1*:*FGF2* and *FGFR1*:*FGFR3* ratios were the differences of variance-stabilized values. The ERK target score was the mean of per-gene z-scores across donors for those of *SPRY1*, *SPRY2*, *SPRY4*, *SPRED1*, *SPRED2*, *DUSP4*, *DUSP6*, *ETV1*, *ETV4*, *ETV5*, *IL17RD* and *FLRT3* that passed the filter. The score was computed only where at least four of these genes passed, which held in c0 to c5. Each measure was regressed on grade with a linear model including technology and the mean-centered capture-burden covariate (score ~ technology + burden + grade; technology was dropped where only one chemistry remained). Benjamini-Hochberg correction was applied across clusters within each measure, and Spearman correlations with grade were reported alongside. For the single-nucleus multiome comparison (Figure S18d; 24 donors), RNA and ATAC counts from the same nuclei were summed per donor. RNA was expressed as log2(CPM + 1). Promoter accessibility was log2(CPM + 1) of the summed counts of peaks overlapping ±1 kb of every annotated transcription start site of the gene, with all *FGF1* promoters pooled. Each was correlated with grade by Spearman correlation (nominal $p < 0.05$).

#### *FGF1* export and heparan sulfate-modifying genes

Genes for non-classical *FGF1* export (*S100A13*, *SYT1*), heparan sulfate (HS) 6-O sulfation (*HS6ST1*-3) and 6-O removal (*SULF1*, *SULF2*), 3-O sulfation (*HS3ST1*, *HS3ST3A1*), HS core proteins and release (*GPC5*, *HSPG2*, *HPSE2*) and the *FGF18*-*FGFR3* axis were taken from the same continuous-grade model (Figure S19a). *FGF18* and *FGFR3* detection was summarized as the number of donors with at least 10 pseudobulk counts and the median pseudobulk CPM per cluster. Donor-level measures used the pseudobulk, filtering and variance stabilization described above, with *FGF1* and *FGF2* always retained. The HS 6-O sulfation balance was the mean variance-stabilized expression of the HS6ST genes passing the filter minus that of *SULF1* and *SULF2* (Figure S19b). It was estimable in c0 to c6 and was tested on grade with the same covariate-adjusted linear model and Benjamini-Hochberg correction across clusters. Coupling with *FGF1* across donors within each cluster (Figure S19c-d) was the Spearman correlation between *FGF1* and *SULF2*, *SULF1* or *S100A13* (variance-stabilized), both raw and with grade partialed out. The partial coefficient was the Pearson correlation between the residuals of rank(x) and of rank(y), each regressed on rank(grade). Benjamini-Hochberg correction was applied across clusters for each gene, separately for raw and partial coefficients.

#### Single-nucleus multiome processing

Paired snRNA-seq and snATAC-seq libraries (10x Genomics Single Cell Multiome ATAC + Gene Expression, chemistry ARC-v1, barcode whitelist 737K-arc-v1) from 24 of the 30 donors were processed with Cell Ranger ARC 2.0.2 against the GRCh38 reference (refdata-cellranger-arc-GRCh38-2020-A-2.0.0). Each multiome sample was matched to its snRNA-seq donor by the MD5 checksum of its filtered feature-barcode matrix rather than by file name. Analyses used Seurat 5.1.0 and Signac 1.13.0 in R

4.3.1, with gene annotation from EnsDb.Hsapiens.v86 (UCSC chromosome names, chr1-22, X and Y) and genome sequence from BSgenome.Hsapiens.UCSC.hg38. The random seed was 42.

Nucleus quality control. For each sample, nuclei were kept if they met all of the following:

1,000 < RNA UMIs < 25,000 (the same thresholds used for the snRNA-seq reference);

mitochondrial reads < 5%;

nucleosome signal < 2;

transcription start site (TSS) enrichment > 1.

Nuclei with an undefined metric failed that criterion. This retained 119,907 of 149,339 nuclei. RNA counts were normalized per sample with SCTransform, regressing out the mitochondrial percentage.

Unified peak set. The per-sample Cell Ranger ARC peak calls were merged (GenomicRanges::reduce) into one peak set. Peaks were restricted to standard chromosomes and to widths of 20-10,000 bp, giving 247,013 peaks. Fragments from every sample were then re-counted against this set (Signac FeatureMatrix) to form a "peaks" assay. No ATAC count filter was applied before re-quantification. After re-quantification, nuclei were kept if they had more than 500 fragments in peaks, leaving 104,181 nuclei. This floor was the highest of eight candidates (0-2,000) at which cluster composition remained effectively unchanged:

1.02-fold spread in per-cluster retention;

standard deviation of the log2 composition shift 0.008.

Integration. RNA was integrated across the 24 samples with SCT-based CCA anchors:

3,000 integration features;

1-50 dimensions;

IntegrateData with features.to.integrate capped at 2,267, the largest feature number that fits within the 2^31 sparse-index limit at the binding merge of this anchor set.

Latent semantic indexing (TF-IDF, then SVD, using features with at least 5 counts) was run on the peaks assay. Component 1 correlated with sequencing depth and was excluded. LSI dimensions 2-40 were integrated across samples with IntegrateEmbeddings using the same anchors. The ATAC UMAP (Figure 9a, Figure S22a) was computed from the integrated LSI, dimensions 2-40. A weighted-nearest-neighbor graph was also built from RNA PCA dimensions 1-50 and integrated LSI dimensions 2-40. Its clusters (resolution 0.8) were not used to assign identity.

Cluster labels. The multiome RNA libraries are the same libraries as the snRNA-seq data for these donors. Cluster labels were therefore transferred by exact barcode join, not by statistical label transfer:

The reference was the 108,600-nucleus snRNA-seq chondrocyte partition (9 clusters), restricted to the 24 multiome donors (96,175 nuclei).

Each reference donor was paired with the multiome sample sharing the most barcodes. Pairing was 1:1 for all 24 donors.

Labels were joined on sample plus barcode.

81,670 of 104,181 nuclei (78.4%) received a label: c0 *PRG4*+ superficial 32,876; c1 *ACAN*-high low-signal 15,058; c2 *COL10A1*+ hypertrophic 14,475; c3 pre-hypertrophic 7,517; c4 *CYTL1*+ 4,910; c5 *ANXA1*+ 2,755; c6 *SLC14A1*+ 1,479; c7 *CRISPLD2*+ 1,559; c8 unassigned 1,041. Unlabeled nuclei, which were non-chondrocytes or nuclei absent from the snRNA-seq partition, were removed. All analyses below use these 81,670 chondrocyte nuclei.

Grade metadata. Macroscopic knee grade was taken from the donor sheet after correction against the sequencing record. Normal_5 is grade 1, and the TKA_4 library is assigned to donor TKA_5. All grade-dependent multiome analyses were re-run after this correction. The 24 donors were:

| Grade | Donors |
|---|---|
| 0 | 3 |
| 1 | 9 |
| 1.5 | 3 |
| 2 | 1 |
| 3 | 1 |
| 4 | 7 |

For display only, donors were grouped into stages: Normal (grade 0-1, n = 12), early OA (1.5-2, n = 4) and severe OA (3-4, n = 8). All statistical tests used continuous grade.

**Differential accessibility against grade**

Pooled model (Figure 9b). Fragment counts in peaks were summed per donor across all labeled nuclei (Seurat AggregateExpression). Peaks were kept if they had at least 10 counts in at least 6 donors (25% of donors), leaving 234,481 peaks. Accessibility was modeled in DESeq2 1.40.2 as ~ macro_knee_grade, with grade continuous. Because donor was the unit of replication, it could not be included as a covariate. The grade coefficient (log2 fold change per grade unit) was tested with the Wald test. Benjamini-Hochberg (BH) adjusted p values used independent filtering at $\alpha = 0.05$; 175,071 peaks were tested. DESeq2 substituted a local dispersion fit because the parametric trend did not fit. It also replaced outliers in 360 peaks (minimum 7 replicates). Peaks with adjusted $p < 0.05$ were called grade-associated: 16,746 in total, 10,003 opening and 6,743 closing.

Grade-associated peaks were annotated with ChIPseeker against TxDb.Hsapiens.UCSC.hg38.knownGene, with promoters defined as ±3 kb of a TSS and nearest-gene symbols from org.Hs.eg.db. Their genomic context was compared with all 158,325 tested peaks that were not associated with grade (Figure S22b).

Leave-one-donor-out robustness. The model was refitted 24 times, each time leaving out one donor. Log2 fold changes were compared with the full model by Pearson correlation on the peaks tested in both fits ($r \geq 0.985$ in all refits). Tissue source could not be included as a covariate, because it is aliased with grade 4: all six arthroplasty knees are grade 4, and only one grade-4 knee is cadaveric.

Per-cluster model (Figure 9c, Figure S22c-e). Counts were summed per donor and cluster. A donor-cluster unit was used if it contained at least 20 nuclei, and a cluster was tested if at least 8 donors qualified. The same filter (at least 10 counts in 25% of that cluster's donors, rounded to the nearest whole donor), the same design (~ macro_knee_grade) and the same threshold (adjusted $p < 0.05$, BH within cluster) were applied, fitting DESeq2 to every peak that passed this filter in the cluster. Donors and peaks tested per cluster:

| Cluster | Donors | Peaks tested |
|---|---|---|
| c0 | 24 | 136,471 |
| c1 | 24 | 70,605 |
| c2 | 24 | 75,187 |
| c3 | 24 | 44,300 |
| c4 | 23 | 26,398 |
| c5 | 23 | 12,366 |
| c6 | 19 | 4,451 |
| c7 | 18 | 10,504 |
| c8 | 17 | 5,275 |

Peaks significant in at least one cluster were cross-tabulated by the number of clusters in which they were significant and by overlap with the pooled set (Figure S22c). For c0-c4, pairwise overlap was computed as the fraction of one cluster's hits that were also significant in the other (Figure S22d).

**Peak-gene links**

Peaks were linked to genes with Signac LinkPeaks:

expression from the SCT assay;

peak counts from the peaks assay;

peaks within 500 kb of the gene TSS;

genes and peaks detected in more than 10 nuclei.

Signac defaults applied otherwise:

Pearson correlation;

200 background peaks matched on GC content (RegionStats on BSgenome.Hsapiens.UCSC.hg38), total accessibility and peak width;

a link was retained at absolute score > 0.05 and z-test $p < 0.05$, with no correction for multiple testing.

Links were computed across all 81,670 nuclei without reference to grade or cluster. This gave 5,712 links between 4,939 peaks and 1,355 genes. Links are correlations of accessibility with expression and do not demonstrate regulation.

Links crossed with grade-associated peaks (Figure 9e-f, Figure S22e). Pooled grade-associated peaks were intersected with the links (922 links, 479 genes). Genes with two or more such peaks were classified as all-opening, all-closing or mixed. The same intersection was made with each cluster's significant peaks.

**Motif accessibility (chromVAR)**

Motif positions for the JASPAR2020 CORE collection were annotated in all peaks with Signac AddMotifs (motifmatchr). The collection was restricted to human (taxonomy 9606) and to the latest version of each matrix, giving 633 matrices. Per-nucleus deviation z-scores were computed with Signac RunChromVAR (chromVAR 1.24.0) using its default GC- and accessibility-matched background peaks.

Grade association (Figure 9d). z-scores were averaged per donor within each cluster. A donor contributed if it had at least 20 nuclei in that cluster, and a cluster was tested if at least 8 donors qualified (17-24 donors per cluster). For each motif and cluster, the donor means were correlated with grade by Spearman correlation, with BH correction across the 633 motifs within each cluster. Of 5,697 cluster-motif tests, 625 reached nominal $p < 0.05$ and three reached adjusted $p < 0.05$ (*RUNX2* in c0 and c5; *PITX2* in c4). Figure 9d therefore shows the 21 motifs with nominal $p < 0.05$ in at least seven of the nine clusters. Correlations from individual nuclei were not used for inference, because they are pseudoreplicated.

Cluster identity in motif space (Figure S22f). Donor-by-cluster mean z-scores were calculated from units with at least 25 nuclei, in clusters with at least 100 nuclei and at least 4 such donors. They were fitted with limma (~ 0 + cluster + donor). Each cluster was contrasted with the mean of the other clusters, using empirical-Bayes moderated t statistics and BH correction within each contrast. One representative motif per transcription-factor family was shown: the motif with the largest absolute difference.

SMAD motif families (Figure 11c). Two sources of SMAD motifs were used.

JASPAR motifs: the JASPAR SMAD3 (MA0795.1) and SMAD2::SMAD3::SMAD4 (MA0513.1) motifs were taken from the per-cluster analysis above (BH across 633 motifs).

cisTarget motifs: 111 cluster-buster files from the cisTarget v10nr_clust collection were converted to position frequency matrices. They covered eight SMAD2/3 motifs and every motif directly annotated to *SMAD1*/5, *CEBPA*/B/D, *RUNX2*, *SOX9*, *DLX5*, *DLX3* or *SP7*. These were scored on the same peaks with AddMotifs and RunChromVAR.

A metacluster's score was the mean z of its member matrices.

The SMAD2/3 family score was the mean of eight motifs: homer CTGTCTGG_Smad2, HOCOMOCO SMAD2_MOUSE.H11MO.0.A, JASPAR MA1964.1, homer TWGTCTGV_Smad3, SwissRegulon hs SMAD3, hdpi SMAD3, metacluster_91.2 and metacluster_50.1.

Donor-by-cluster family means (at least 10 nuclei per unit and at least 8 donors per cluster) were correlated with grade by Spearman correlation, with BH correction across the nine clusters within the family.

**Chromatin coverage at selected loci**

*FN1* and *TNFSF11* (Figure S23). Coverage was drawn with Signac CoveragePlot from the fragments of all labeled nuclei, grouped by stage:

60 kb added on either side of the gene;

called peaks and peak-gene links shown beneath;

violin plots of SCT expression of the gene in the same nuclei.

*ITGA5*, *ABCA1* and *GPX3* (Figure S24). For each gene, the two linked grade-associated peaks with the lowest adjusted p were plotted per stage, using the peak ±1.5 kb, each panel scaled to its own window. For all of the gene's linked grade-associated peaks (six, seven and eight peaks, all closing), a donor accessibility score was calculated:

1. log2(CPM + 1) of each peak in the per-donor pseudobulk, with CPM relative to total peak counts;
2. z-scored across donors per peak;
3. averaged across peaks.

The score was correlated with grade across 24 donors by Spearman correlation. Stage grouping was used for display only.

***FN1* production, assembly machinery and *HTRA1***

Per-cluster grade association (Figure 10a). Per-cluster log2 fold changes per grade unit were taken from the snRNA-seq continuous-grade pseudobulk model (see snRNA-seq grade model): DESeq2 ~ technology + capture burden + grade on summed raw counts per donor and cluster. Units had at least 20 nuclei, clusters needed at least 8 donors and 3 distinct grades, and genes needed at least 10 counts in at least 50% of the cluster's units. Log2 fold changes were not shrunk, and adjusted p values were BH within cluster with α = 0.05. Donors per cluster: c0-c2 30, c3 29, c4 28, c5 28, c6 24, c7 21, c8 19. The genes shown were:

*FN1*;

the fibril-assembly genes *ITGA5*, *ITGB1*, *ITGAV*, *ITGB3*, *TNS1*, *TLN1*, *FERMT2*, *ILK*, *PXN* and *SDC4*;

the fibronectin-cleaving proteases *HTRA1*, *MMP2*, *MMP3* and *MMP14*.

Genes below the expression filter in a cluster were not tested.

Donor-level coupling (Figure 10b-c). Within each cluster, pseudobulk counts of qualifying donors were filtered as above, with *FN1* always retained, and variance-stabilized with DESeq2 vst (blind = TRUE). *FN1* and *HTRA1* were correlated across donors by Spearman correlation. The partial correlation given grade was computed by regressing the ranks of each gene on the rank of grade (linear regression) and correlating the residuals (Pearson), and is referred to here as partial Spearman. P values were BH-adjusted across the nine clusters, separately for raw and partial coefficients.

In a sensitivity analysis, the rank of the SMAD3 core score (see TGF-β/SMAD signaling) was added to the covariates.

Chromatin and RNA at *FN1*, *HTRA1*, *ITGA5* and *CD44* (Figure 10d-f). Peak counts and raw RNA counts were summed per donor over all 81,670 labeled nuclei.

Figure 10d: for every peak linked to *FN1* (23 peaks), *HTRA1* (11), *ITGA5* (21) or *CD44* (17), accessibility in CPM (relative to the donor's total peak counts) was correlated with grade across the 24 donors by Spearman correlation. These coefficients are descriptive and were not tested. Distance was measured from the peak center to the linked TSS, with upstream negative relative to gene strand. Peaks in the pooled grade-associated set (adjusted $p < 0.05$) were marked.

Figure 10e: linked-region accessibility of a gene was the sum of counts over all of its linked peaks, expressed as log2(CPM + 1). Multiome RNA was log2(CPM + 1) of the gene's counts relative to the donor's total RNA counts. Each was expressed as the difference from the mean of the Normal stage and summarized as mean ± standard error per stage.

Figure 10f: *FN1* linked-region accessibility was correlated with *FN1* RNA across the 24 donors by Spearman correlation.

#### TGF-β/SMAD signaling, *FN1* receptors and the *CD44* axis

Pathway genes (Figure 11a). Grade associations of TGF-β ligands and latent-complex genes, receptors and activators, and transducers and target genes were taken from the same per-cluster snRNA-seq model as Figure 10a:

ligand and latent complex: *TGFB1*-3, *LTBP1-3*;

receptors and activators: *TGFBR1*-3, *ACVR1*, *ITGAV*, *ITGB8*, *MMP14*, *ITGA5*, *ITGB1*, *CD44*;

transducers and targets: *SMAD3*, *SMAD7*, *SKIL*, *PMEPA1*, *SERPINE1*, *CCN2*, *COMP*, *TGFBI*.

Target scores (Figure 11b). Within each cluster, pseudobulk counts were filtered and variance-stabilized as for Figure 10b. Each gene was z-scored across donors, and a set score was the mean z over the set's expressed genes; a set needed at least 4 expressed genes. Three sets were scored:

SMAD3 early-response ("SMAD3 core") set: *SERPINE1*, *SMAD7*, *SKIL*, *PMEPA1*, *TGFBI*, *CCN2*, *JUNB*, *BHLHE40*, *TGIF1*, *KLF10*, *SNAI2*, *SKI*, *LTBP1*, *COL1A1*, *TIMP3*, *CDKN2B* and *BAMBI*. *FN1*, *COMP* and *LTBP2* were excluded to avoid circularity. Between 11 and 17 genes per cluster passed the expression filter.

Hallmark TGF-β signaling (MSigDB via msigdbr 7.5.1).

BMP target set: *ID1*-4, *SMAD6*, *SMAD9*, *MSX2*, *BAMBI*, *NOG*, *GREM1*.

Scores were regressed on grade with technology and capture burden as covariates (linear model), with BH correction across clusters within each set.

TGF-β genes with linked grade-associated peaks (Figure 11d). For TGF-β pathway genes and the *FN1*-related genes *FN1*, *HTRA1*, *ITGA5* and *CD44*, linked peaks were counted as opening (log2 fold change > 0, adjusted $p < 0.05$ in the pooled model), closing, or not grade-associated. Genes with at least one grade-associated linked peak were shown, except *JUNB*, an immediate-early gene (1 opening and 5 closing of 27 linked peaks).

*FN1* receptor genes (Figure 11e, Figure S25a).

Clusters: per-cluster estimates for *ITGA5*, *CD44*, *ITGAV* and *ITGB8* were taken from the snRNA-seq model above, with 95% confidence intervals of log2 fold change ± 1.96 × standard error.

Bulk: bulk estimates came from the bulk continuous-grade DESeq2 model ~ batch + detected genes (z) + grade in 92 knees (see bulk RNA-seq section). The bulk log2 fold change was shrunk with lfcShrink (type "normal"), and its adjusted p value came from the unshrunk Wald test.

Hyaluronan-axis genes and *CD44* effectors (Figure S25a): *HAS2*, *HAS3*, *CEMIP*, *HYAL1*, *HYAL2*, *HMMR*, *TNFAIP6*, *HAPLN1*, *EZR*, *MSN*, *CCN1*, *CCN2*, *ADAMTS5* and *MMP3*, shown in the same format.

Coupling beyond grade (Figure 11f, Figure S25b). Within each cluster, the following were correlated across donors with *CD44*, *ITGA5* and *ITGB8*:

variance-stabilized expression of single genes, which were always retained regardless of the expression filter;

mean-z program scores, each requiring at least 2 genes:

adhesion core (14 genes);

ERM-Rho (9);

catabolic (10);

YAP targets (MSigDB CORDENONSI_YAP_CONSERVED_SIGNATURE);

Hallmark TNF-α signaling via NF-κB;

KEGG focal adhesion (extracellular matrix ligands and integrins removed);

hyaluronan synthesis (*HAS1*-3);

pericellular hyaluronan (*ACAN*, *VCAN*, *HAPLN1*, *PRG4*, *TNFAIP6*).

*FN1*, *CD44*, *ITGA5*, *ITGB1*, *ITGAV*, *ITGB8*, *HAS1*-3, *COL1A1*, *COL1A2* and *COMP* were excluded from all program sets. Partial correlations were computed by regressing the ranks of both variables on rank(grade), capture burden and technology, then correlating the residuals (Pearson). P values were BH-adjusted across clusters for each target-measure pair. Figure 11f shows the median partial correlation across the nine clusters and the number of clusters with adjusted $p < 0.05$.

For Figure S25b, *CD44* and *ITGA5* expression were each residualized within cluster on grade, capture burden and technology (linear model). All donor-cluster pairs were then pooled ($n = 239$) and correlated by Spearman correlation. Because the pairs are not independent, this correlation is descriptive.

#### Candidate transcriptional regulators of *FN1*

Prior regulatory edges. *FN1* regulators were taken from two sources:

CollecTRI: human interactions downloaded from the OmniPath web service (omnipathdb.org, CollecTRI dataset) on October 7, 2026; OmniPath does not assign a release number to this resource. AP-1 and NF-κB dimers were collapsed to AP1 and NFKB, other complexes were removed, and the mode of regulation was set to −1 for inhibitory-only edges and +1 otherwise, with edges unweighted. This gave 70 *FN1* regulators.

DoRothEA: edges of confidence A-C (dorothea 1.14.1), giving 9 *FN1* edges.

Transcription-factor activity. Activity was inferred with the decoupleR 2.8.0 univariate linear model (run_ulm, minimum 5 targets) using CollecTRI regulons. *FN1* was removed from every regulon, and transcription factors with at least 5 targets were retained (758). Activity was inferred in three datasets:

snRNA-seq (30 donors): vst-transformed donor-by-cluster pseudobulks. Units had at least 20 nuclei, clusters at least 8 donors, and genes at least 10 counts in at least 50% of donors, with *FN1* and candidate transcription factors always retained.

Bulk (92 knees): vst-transformed bulk counts. Genes needed at least 10 counts in at least 25% of knees (17,350 genes, with candidate transcription factors always retained); duplicate gene symbols were resolved by highest total count.

Multiome (24 donors): vst-transformed RNA pseudobulks. Genes needed at least 10 counts in at least 12 donors, with candidate transcription factors always retained.

The SMAD3 core score and other target-set scores were computed as mean z, with *FN1* excluded and at least 3 genes required.

Association with grade was tested with linear models (*FN1* regulators, evidence category E3 below):

snRNA-seq: ~ technology + capture burden + grade, per cluster;

bulk: ~ batch + detected genes (z) + grade.

Coupling with *FN1* beyond grade was tested by partial correlation on rank residuals (Pearson on residuals of rank(variable) regressed on the covariates):

snRNA-seq: rank(grade), technology and capture burden;

bulk: rank(grade), batch and detected genes;

multiome: rank(grade) only. Two *FN1* measures were used here, *FN1* RNA and *FN1* accessibility (the 19 linked opening peaks).

These coupling statistics are nominal and were not corrected for multiple testing.

Motif occurrence in *FN1*-linked opening regions. The 19 *FN1*-linked peaks that opened with grade (pooled adjusted $p < 0.05$) were scanned for motifs:

motifs: the JASPAR2020 CORE vertebrate collection, latest versions (746 matrices);

scanning: motifmatchr defaults ($p < 5 \times 10^{-5}$; base composition from each scanned set).

The scanned regions were compared with a GC- and width-matched background, built by assigning all peaks to bins of 20 GC-content quantiles × 5 width quantiles and sampling 50 non-*FN1* peaks per *FN1* peak from the same bin (seed 7). Enrichment was tested with a one-sided Fisher's exact test, with BH correction across the 746 motifs. cisTarget v10nr_clust family motifs were tested in the same way, counting a family hit if any member matched.

Candidate ranking (Figure S26a). Each candidate (33 curated factors and pathways and the data-driven additions below) was scored on four evidence categories:

| Category | Criterion |
| --- | --- |
| E1 | positive CollecTRI or DoRothEA (A-C) FN1 edge |
| E2 | JASPAR motif enrichment at nominal $p < 0.05$ with ≥4 of 19 regions hit, or cisTarget family $p < 0.05$ |
| E3 | activity, target-set score or the factor's own RNA rising with grade in ≥3 of 9 clusters, and in bulk, at nominal $p < 0.05$ |
| E4 | positive coupling of activity, target-set score or the factor's own RNA with *FN1* at nominal $p < 0.05$ in ≥3 clusters, in bulk and in the multiome (*FN1* RNA or accessibility) |

Candidates were ranked by the number of categories met, then by coupling percentile. Transcription factors not among the curated candidates were added as data-driven rows if their activity met E3 and E4 and ranked at or above the 90th percentile of *FN1* coupling among all 758 activities in both the snRNA-seq and bulk data.

SMAD3-conditioned coupling (Figure 11g). Partial correlations with *FN1* were recomputed with rank(SMAD3 core score) added as a covariate; for the SMAD3 core score itself, *SOX5* activity was added instead. In this sensitivity analysis, the snRNA-seq covariates were rank(grade) and technology. The median across clusters is shown.

Figure 11h. This panel shows bulk *FN1* against *SOX5* activity, *FOSL1* activity and the SMAD3 core score, each as residuals of ranks on rank(grade), batch and detected genes.

Footprinting (Figure S26b-c). Footprints were computed from Tn5 insertions read directly from the Cell Ranger ARC fragment files. These files are already Tn5-shift-corrected, so the two insertions per fragment were taken at start + 1 and end. Only nuclei with a cluster label were used, and analyses were run on all labeled chondrocytes and on c0 separately.

Motif sites: sites for 13 JASPAR2020 motifs (*CTCF* MA0139.1, *SOX5* MA0087.1, *FOSL2* MA0478.1, *RUNX2* MA0511.2, *SOX9* MA0077.1, *FOSL1* MA0477.2, SMAD3 MA0795.1, SMAD2::SMAD3::SMAD4 MA0513.1, *TEAD1* MA0090.3, *TEAD4* MA0809.2, *FOXD1* MA0031.1, *NFKB1* MA0105.4, *RELA* MA0107.1) were located within the multiome peaks on chromosomes 1-22 and X with motifmatchr ($p < 5 \times 10^{-5}$) and centered on the motif midpoint with strand orientation.

Profiles: insertions were counted per donor in ±250 bp windows around each site, with minus-strand sites reversed.

Tn5 bias model:

The insertion hexamer (positions −3 to +2) was tabulated from $10^6$ random insertions per donor, pooled across donors.

Its frequency was divided by the hexamer frequency in all peak sequences to give a per-hexamer bias.

An expected profile per motif was the mean bias at each offset over up to 20,000 sampled sites (seed 7).

Correction: observed and expected profiles were each normalized to their mean at 200-250 bp from the center, and the expected profile was subtracted from the observed.

Footprint depth: mean corrected flank signal (from the motif half-width to half-width + 50 bp on each side) minus mean corrected core signal (within the motif half-width).

Depth was correlated with stage (Normal 0, early OA 1, severe OA 2) across the 24 donors by Spearman correlation. The same test was run in the 18 non-arthroplasty donors, and in a linear model adjusting for log10 insertions in peaks. The depth and flank stage correlations in all donors were BH-adjusted across the 13 motifs; the non-arthroplasty and insertion-adjusted tests were nominal. Aggregate profiles were summed per stage before normalization and correction (Figure S26c).

**Statistics and reproducibility**

Unless stated otherwise, tests were two-sided, multiple testing was controlled by the Benjamini-Hochberg procedure, and the significance threshold was adjusted $p < 0.05$. Donors, not nuclei, were the unit of replication for all single-nucleus inferences. For bulk RNA-seq, reported log2 fold changes are shrunk with a normal prior (DESeq2 lfcShrink, type "normal") and p values come from the unshrunk Wald test; calls that depend on the direction of change (robustness to library quality, replication and response shapes) used the unshrunk estimates. Single-nucleus log2 fold changes are unshrunk maximum-likelihood estimates. Because grade 4 cartilage was obtained almost entirely at arthroplasty, analyses in cadaveric knees only are reported as sensitivity analyses and were not required for significance. Random seeds were fixed (42 unless stated otherwise).

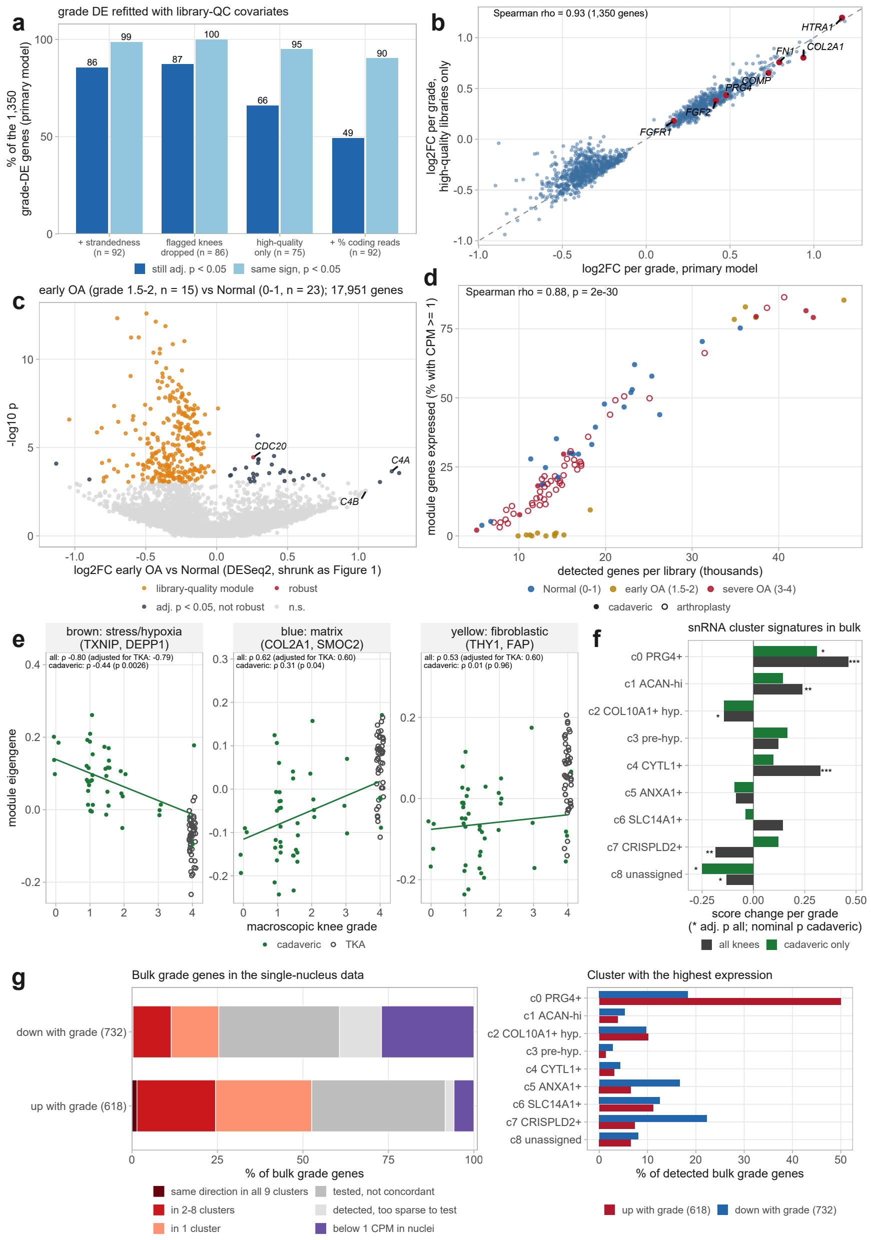


**Figure S1. Robustness of the grade analysis to RNA quality, comparison of early OA with Normal cartilage and other analyses of the bulk cohort.** (a) The 1,350 genes associated with grade (Figure 1b) refitted with library-quality covariates (strandedness, fraction of coding reads), after excluding libraries flagged for low quality, or restricted to high-quality libraries (reverse-strand share ≥ 0.85 and ≥ 30% coding reads; 75 knees): percentage still significant (adjusted $p < 0.05$) and with the same sign at $p < 0.05$. (b) Log2 fold change per grade unit in the primary model against the refit restricted to high-quality libraries, both shrunk with a normal prior. **(c)** Early OA (grade 1.5-2, n = 15) against Normal (grade 0-1, n = 23) with the model of Figure 1 (DESeq2, ~ batch + detected genes + group, all 17,951 genes; log2 fold change shrunk with a normal prior, as in Figure 1). Orange, the library-quality module (287 genes at adjusted $p < 0.05$ detected together in proportion to library complexity); dark gray, other genes at adjusted $p < 0.05$ that did not remain significant with the module as a covariate or with strandedness as a covariate or flagged libraries removed; red, robust. **(d)** Percentage of the module genes expressed (counts per million ≥ 1) in each of the 92 knees against the number of detected genes, colored by grade group (filled, cadaveric; open, arthroplasty). **(e)** Co-expression modules (WGCNA after regressing out library-quality measures with grade protected) against grade; Spearman correlation with grade in all knees, with arthroplasty status, and in cadaveric knees only. The cadaveric-only estimates are shown for information and were not used as a filter, because grade 4 knees were obtained almost entirely at arthroplasty. **(f)** Scores of the snRNA-seq cluster marker signatures in bulk cartilage: change per grade unit in all knees (stars, adjusted p) and cadaveric knees only (stars, nominal p). **(g)** The bulk grade genes (Figure 1b) in the single-nucleus data. Left, share of the genes rising and falling with grade that changed in the same direction in all nine, two to eight or one chondrocyte subset (per-cluster grade model, adjusted $p < 0.05$), were tested without a concordant change, were detected but too sparse to test, or were barely detected in chondrocyte nuclei (mean counts per million below 1 in every subset). Right, for the detected genes, the subset with the highest donor-averaged expression. * $p < 0.05$, ** $p < 0.01$, *** $p < 0.001$.

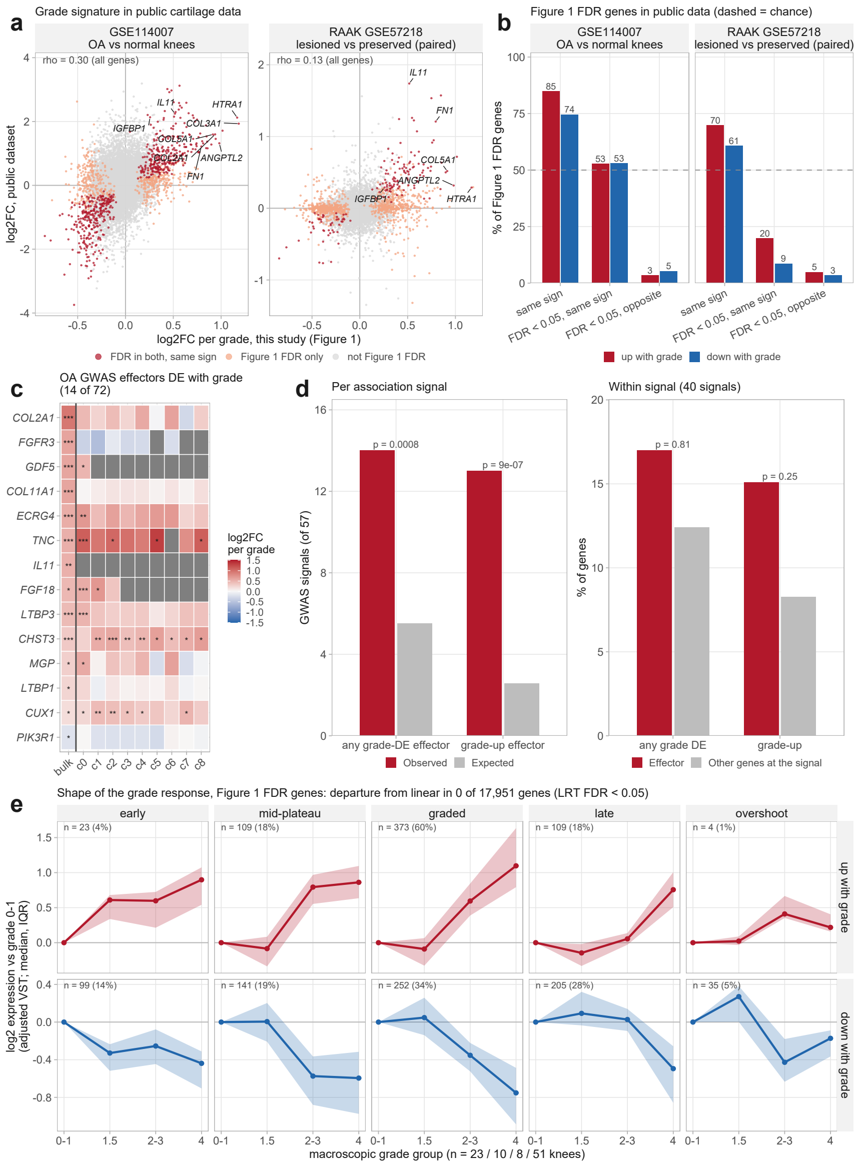


**Figure S2. The bulk grade signature is reproduced in direction in public cartilage data, is enriched at OA risk loci and is consistent with a linear change with grade. (a)** Log2 fold change per grade unit in this study (Figure 1) against log2 fold change in GSE114007 (knee cartilage RNA-seq, 20 OA against 18 normal; DESeq2 with sequencing platform as a covariate) (4) and in RAAK GSE57218 (33 pairs of lesioned and macroscopically preserved cartilage from the same OA joint; limma, paired) (22). Red, genes at adjusted $p < 0.05$ in both with the same sign; orange, adjusted $p < 0.05$ in Figure 1 only; rho, Spearman correlation over all shared genes. **(b)** Percentage of the Figure 1 genes at adjusted $p < 0.05$ that changed in the same direction in each dataset, that were also at adjusted $p < 0.05$ there with the same sign, or with the opposite sign, for genes up and down with grade; dashed line, 50% expected for direction by chance. GSE114007 comes from an earlier cohort of our group and is not fully independent. **(c)** High-confidence OA effector genes from genome-wide association (23) that changed with grade in the bulk cohort (14 of the 72 tested), with their change per grade in the bulk cohort and in each chondrocyte cluster (per-cluster grade model); gray, not tested in that cluster. **(d)** Left, association signals (lead variants) with at least one effector gene at adjusted $p < 0.05$ in Figure 1, observed and expected from the genome-wide rate given the number of effector genes per signal (exact Poisson-binomial p), for any direction and for genes up with grade. Right, within signals: percentage of effector genes and of the other genes at the same signals that changed with grade (Cochran-Mantel-Haenszel test stratified by signal). **(e)** Expression of the Figure 1 genes at adjusted $p < 0.05$ in grade groups 1.5, 2-3 and 4 relative to grade 0-1 (n = 10, 8, 51 and 23 knees): difference in mean variance-stabilized expression after adjustment for batch and detected genes (median and interquartile range over genes), grouped on the same values by the fraction of the grade 4 difference reached at intermediate grades: early, at least half at grade 1.5; mid-plateau, at least three quarters by grade 2-3; graded, between; late, less than a quarter by grade 2-3; overshoot, an intermediate group exceeding grade 4 by half. Groups are descriptive (genes kept their group in a median of 44% to 72% of bootstrap resamples of knees); a likelihood-ratio test of grade groups against a linear grade term found no gene departing from linear (adjusted $p < 0.05$) among 17,951. Log2 fold changes from DESeq2 (this study and GSE114007) are shrunk with a normal prior as in Figure 1; RAAK values are limma estimates. Directions used for replication in (a-b) are assigned from the unshrunk estimates. * $p < 0.05$, ** $p < 0.01$, *** $p < 0.001$ (adjusted).

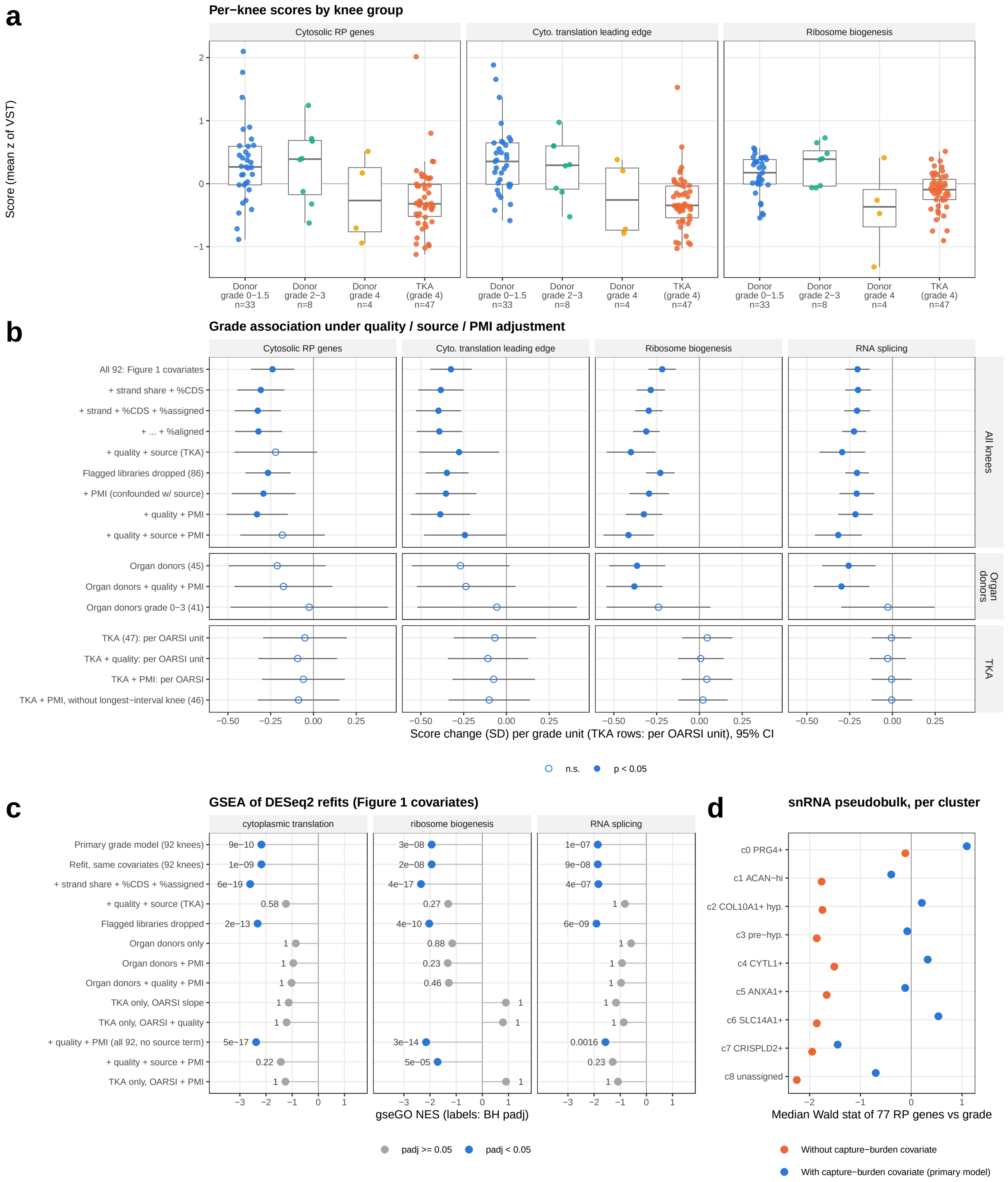


**Figure S3. The decline of translation-related transcripts with grade is a library-level feature of end-stage cartilage.** Bulk RNA-seq of 92 knees. **(a)** Per-knee scores (mean z-scored variance-stabilized expression) of cytosolic ribosomal-protein genes, the leading-edge genes of cytoplasmic translation and ribosome biogenesis, by group: organ donors of grade 0-1.5 (33 knees), 2-3 (8) and 4 (4), and arthroplasty knees (TKA, all grade 4; 47). **(b)** Change in each score per grade unit (standard deviations; 95% confidence interval) under increasing adjustment for RNA quality (RSeQC strand share, coding, assigned and aligned reads), tissue source and post-mortem interval (PMI) in all knees, within organ donors, and within arthroplasty knees per OARSI unit (with and without the knee with the longest interval); filled, $p < 0.05$. The all-knee PMI term largely re-encodes tissue source. **(c)** Normalized enrichment scores of cytoplasmic translation, ribosome biogenesis and RNA splicing in gseGO of the DESeq2 grade model refitted with each adjustment or subset; labels, Benjamini-Hochberg adjusted p. With a source term, the grade effect rests on the four organ-donor grade 4 knees. **(d)** Median Wald statistic against grade of the 77 ribosomal-protein genes in each chondrocyte cluster, in the per-cluster pseudobulk model without and with the capture-burden covariate used in the manuscript.

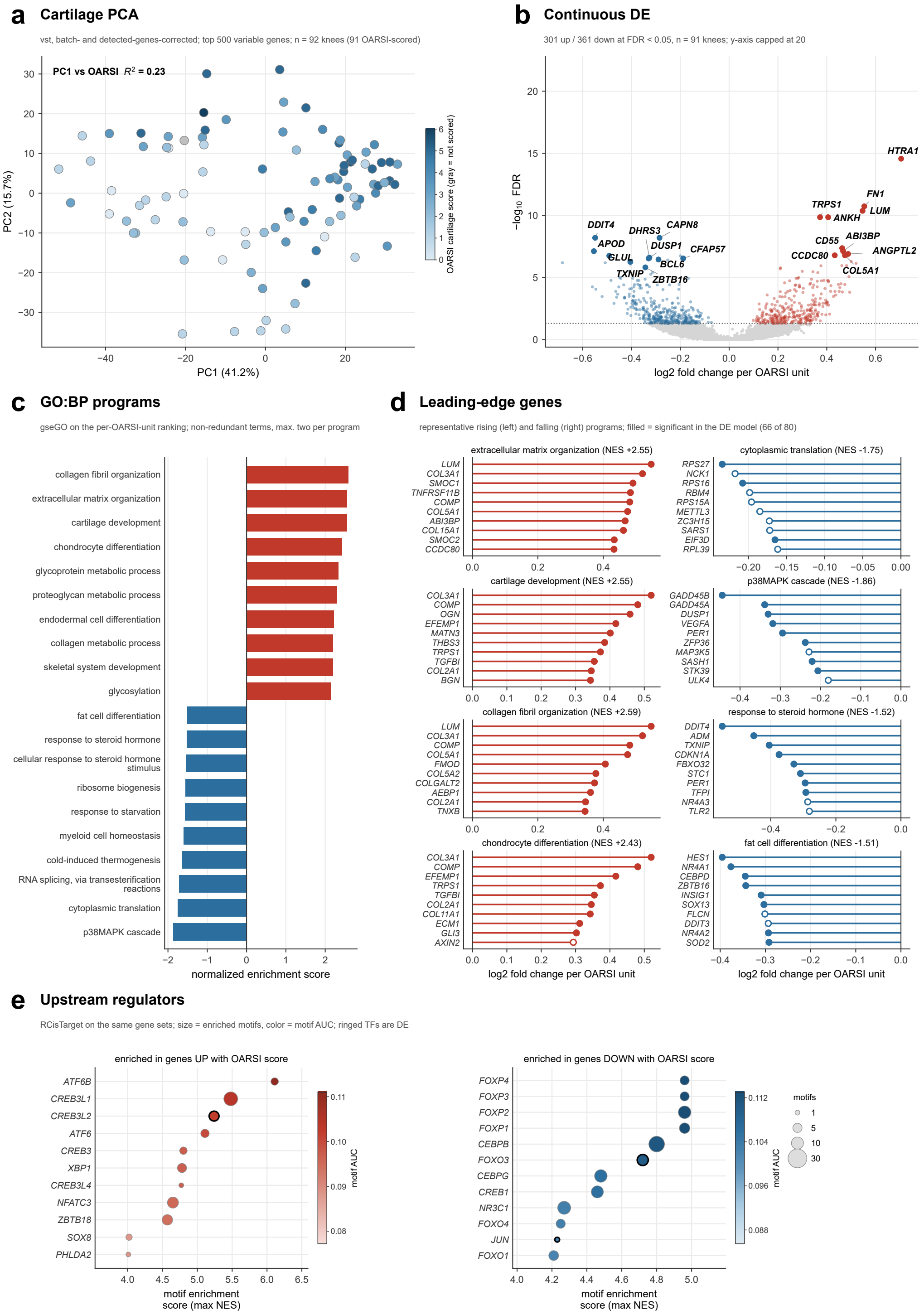


**Figure S4. The cartilage transcriptome changes with the histological OARSI score as with macroscopic grade.** Bulk RNA-seq of articular cartilage analyzed with the OARSI cartilage score of the medial weight-bearing region in place of macroscopic grade (91 knees with a score), with the model and covariates of Figure 1. **(a)** Principal component analysis of Figure 1a, colored by OARSI score; the unscored knee is gray. PC1 correlates with the OARSI score ($R^2$ = 0.23, 91 knees; $R^2$ = 0.36 for grade). **(b)** Genes changing with the continuous OARSI score; 301 rise and 361 fall at adjusted $p < 0.05$ (660 with baseMean ≥ 10), and the most significant in each direction are labeled. The y axis is capped at 20. **(c)** GO biological processes enriched on the same ranking, ten in each direction (rising, the highest normalized enrichment scores among terms of at least 30 genes; falling, at most two per biological program), as normalized enrichment score. **(d)** Leading-edge genes of four rising and four falling processes; filled points are also significant in the gene-level analysis. **(e)** Transcription factor motifs enriched within 10 kb of the transcription start sites of genes rising and falling with the OARSI score (RcisTarget); bubble size is the number of enriched motifs, fill the motif AUC, and a black ring marks a transcription factor that is itself differentially expressed with the OARSI score.

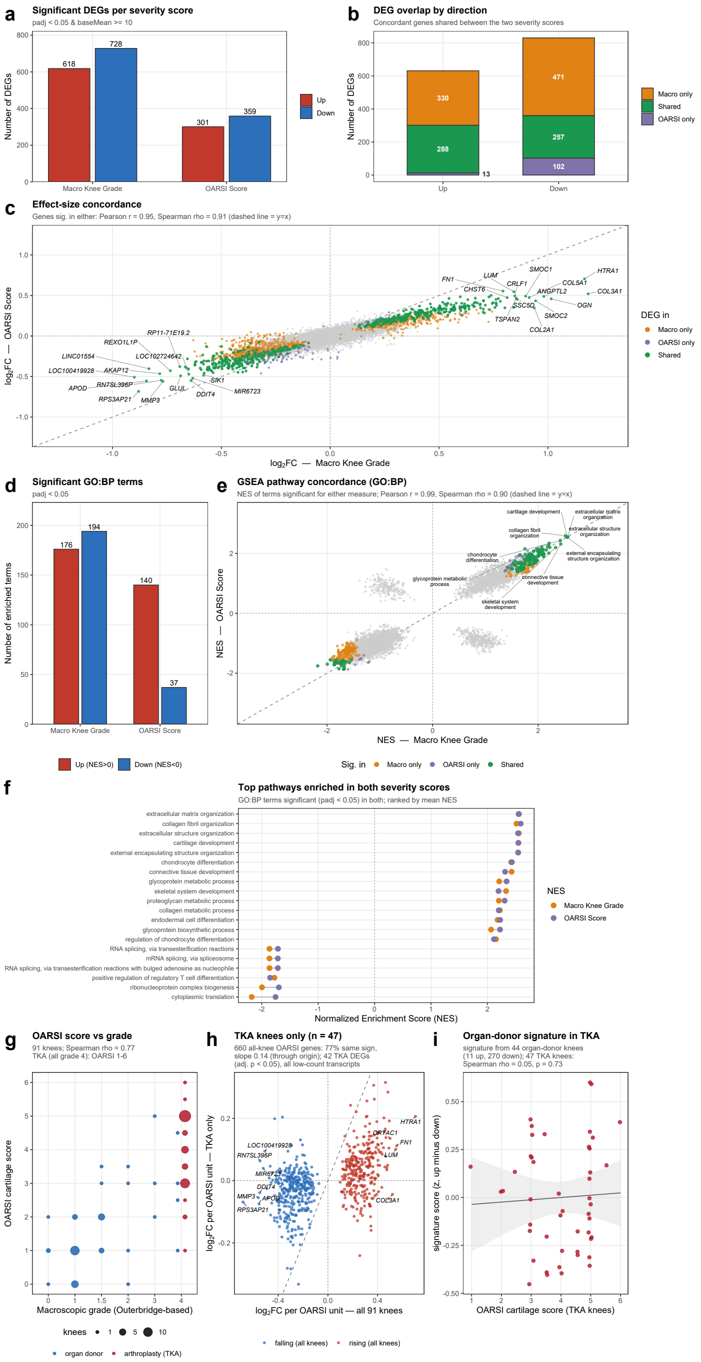


**Figure S5. Macroscopic grade and OARSI score signatures, and the OARSI score within arthroplasty knees.** Bulk RNA-seq of articular cartilage; continuous DESeq2 models with the covariates of Figure 1 (grade, 92 knees; OARSI score, 91 knees). **(a)** Genes associated with each measure (adjusted $p < 0.05$, baseMean ≥ 10). **(b)** Overlap of the rising and falling genes; no shared gene changes in opposite directions. **(c)** Log2 fold changes per unit of grade and of OARSI score for all genes, colored by the measure for which they are significant; dashed line, y = x. Correlations are over the 1,461 genes significant for either measure. **(d)** GO biological processes enriched for each measure (gseGO, adjusted $p < 0.05$). **(e)** Normalized enrichment scores of all processes, colored by the measure for which they are significant; correlations are over the 404 processes significant for either measure (143 shared). **(f)** Normalized enrichment scores of shared processes for both measures: the 14 rising and 6 falling processes with the highest mean absolute score. **(g)** OARSI score of each knee against its macroscopic grade, colored by tissue source, with the two sources plotted side by side; point size, number of knees. Arthroplasty knees, all of grade 4, span OARSI scores 1 to 6. **(h)** Shrunk log2 fold change per OARSI unit of the 660 OARSI-associated genes in all 91 knees and in the 47 arthroplasty knees alone, colored by direction in all knees; dashed line, y = x. Agreement in direction is from the Wald statistics, and the slope from a regression through the origin. The 42 genes significant within arthroplasty knees (adjusted $p < 0.05$) were all low-count transcripts. **(i)** Score of an OARSI signature derived from the 44 organ-donor knees alone (11 rising and 270 falling genes; mean z-scored expression of rising minus falling genes) in each of the 47 arthroplasty knees, against its OARSI score; line, linear fit with 95% confidence band.

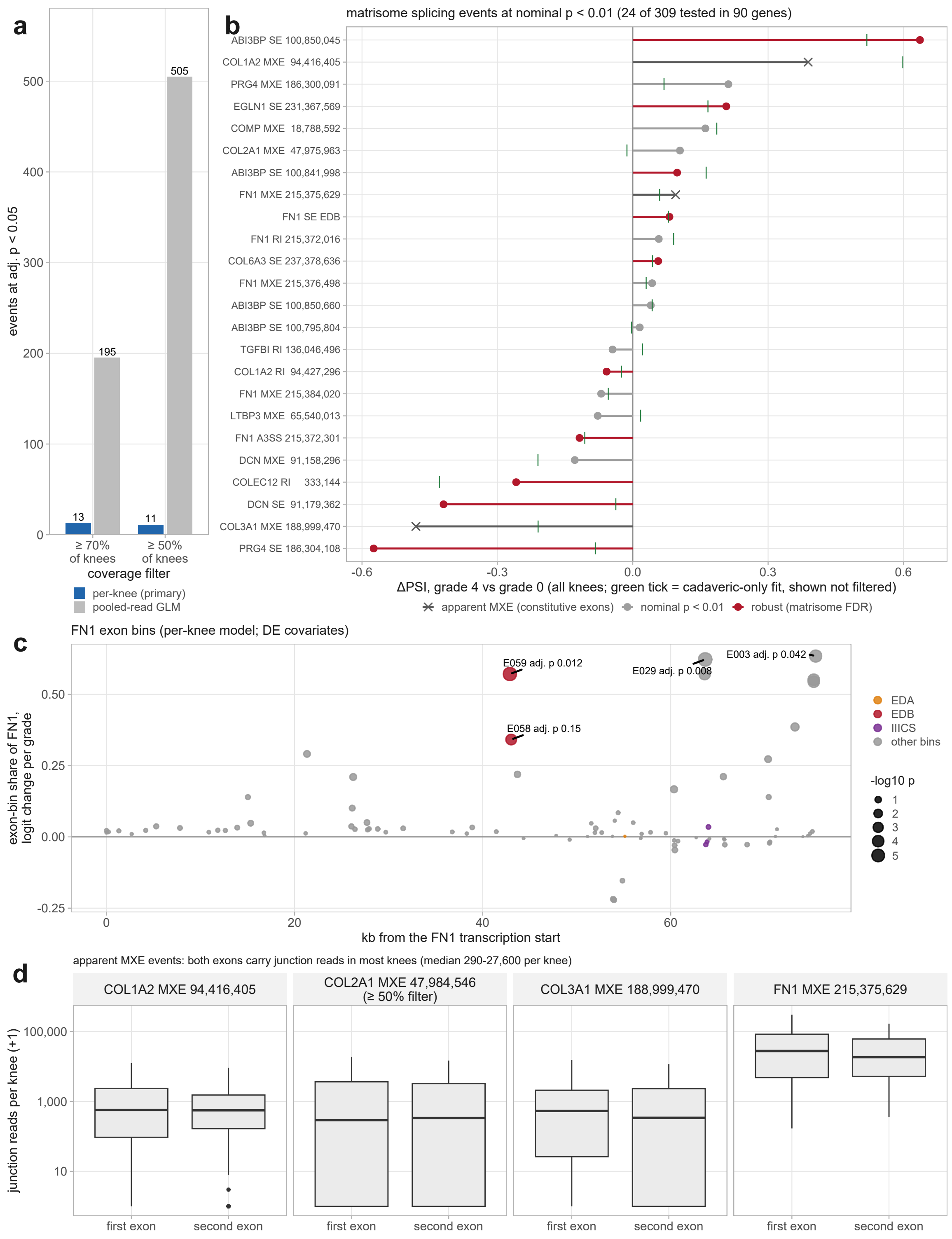


**Figure S6. Genome-wide splicing in bulk cartilage: model comparison, matrix events and *FN1* exon usage. (a)** Number of events at adjusted p < 0.05 with the per-knee model and with a quasibinomial model of pooled junction reads, at two coverage filters; the pooled model is dominated by high-coverage knees. **(b)** All alternative splicing events in matrisome genes with nominal p < 0.01 (70% coverage filter): ΔPSI from grade 0 to grade 4 with the adjusted p computed within the 309 matrisome events; the estimate in cadaveric knees only is shown for information; crosses, the apparent mutually exclusive exon events shown in (d). **(c)** Per-knee exon-bin share along *FN1* (logit change per grade unit, differential expression covariates); the EDB bins are shown in red, EDA and IIICS bins in color; point size, –log10 p. **(d)** Read support for both exons of the apparent mutually exclusive exon events in *COL1A2*, *COL2A1* (from the 50% coverage filter), *COL3A1* and *FN1*: junction reads supporting each exon per knee. Both exons carried at least 10 junction reads in 54 to 75 of the 92 knees for the collagen events and in all 92 for *FN1*, indicating that these events pair two constitutive exons rather than mutually exclusive ones.

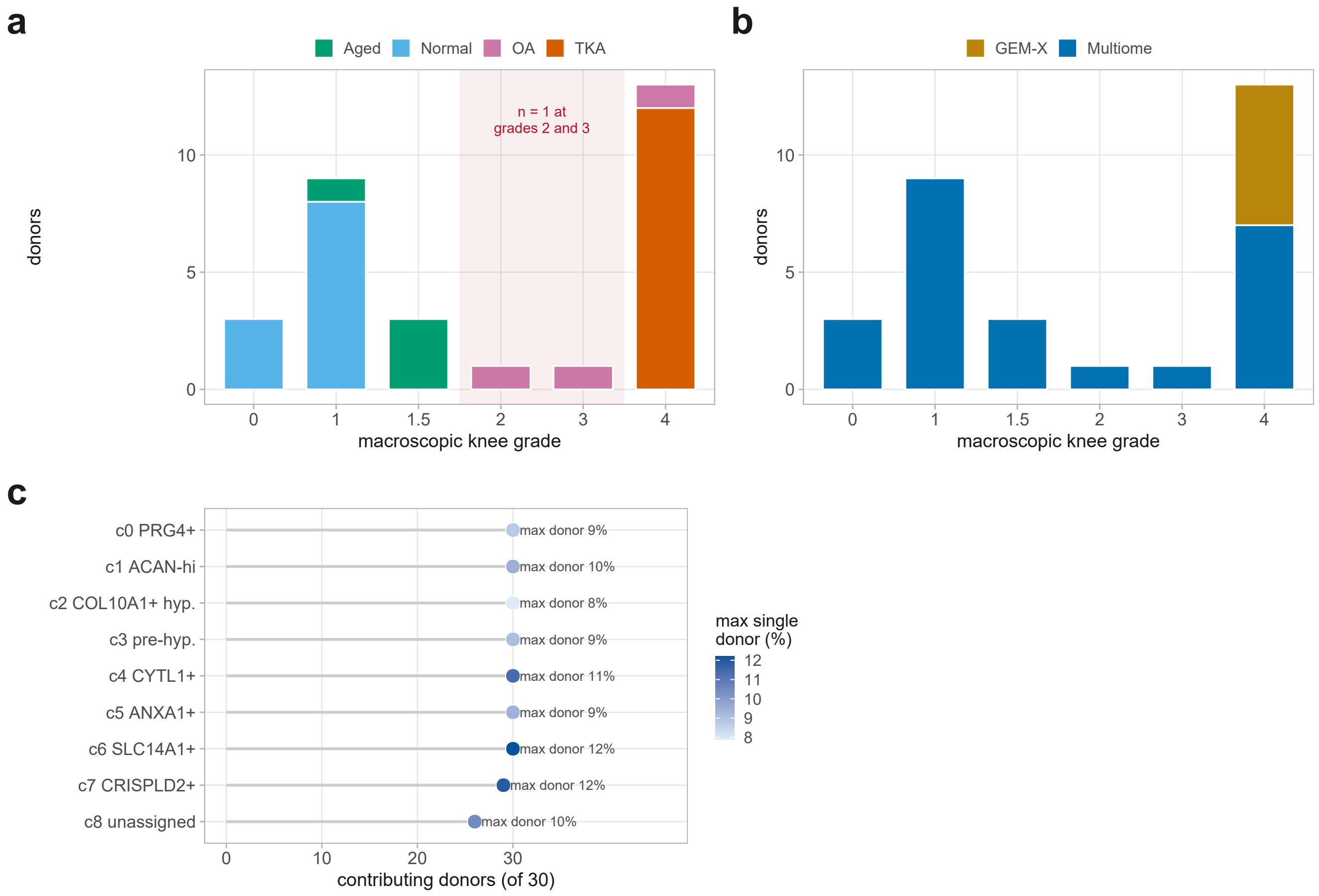


**Figure S7. Cohort composition. (a)** Donors per macroscopic knee grade, colored by clinical group; the shaded band marks grades 2 and 3, where n = 1 each. **(b)** The same distribution by sequencing chemistry. **(c)** Contributing donors per cluster, of 30, with each cluster's maximum single-donor share.

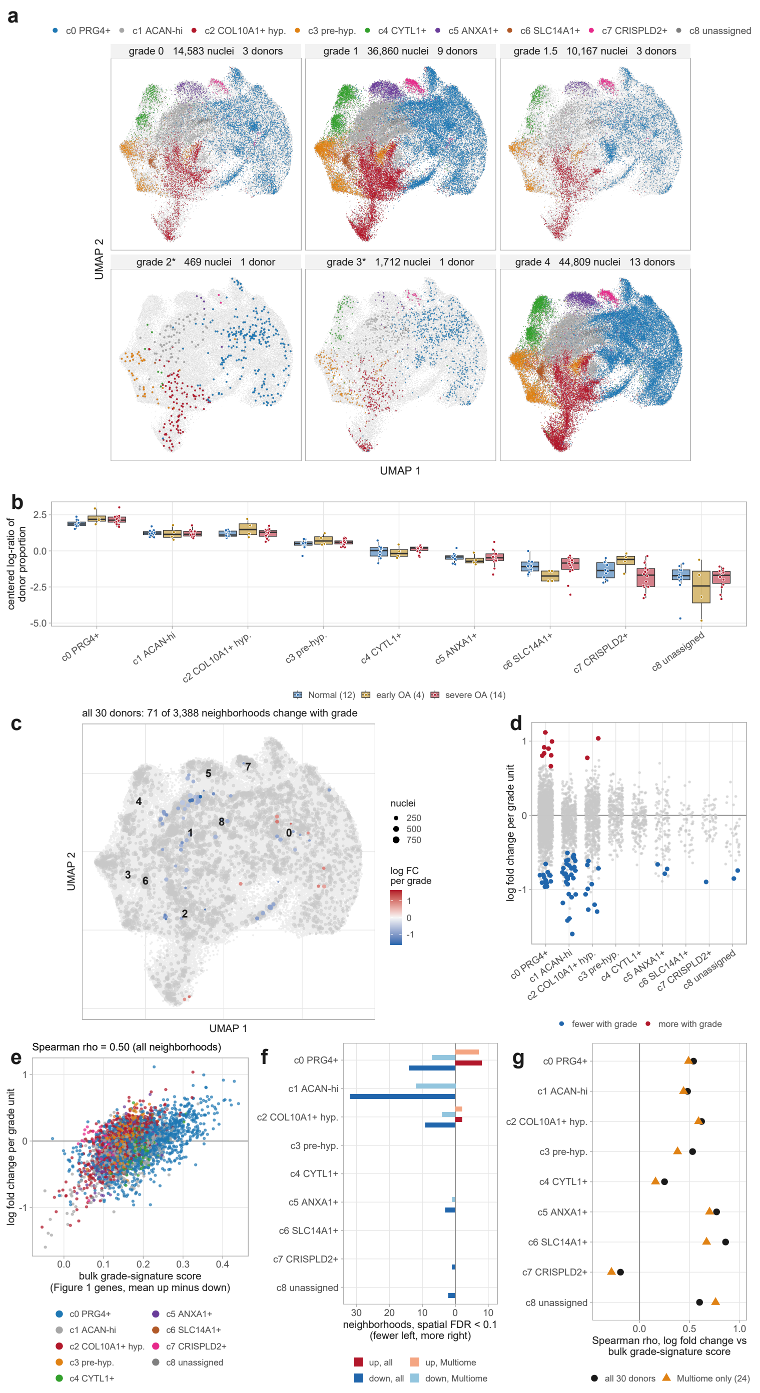


**Figure S8. Cluster abundance by grade and neighborhood differential abundance. (a)** The UMAP of Figure 3a split by grade; gray shows all 108,600 nuclei in every panel and color shows the nuclei of that grade by cluster. Point size and opacity are scaled per panel by nuclei count, and * marks a grade contributed by a single donor. **(b)** Cluster abundance by stage, as the centered log-ratio of each donor's proportion over the complete nine-cluster composition (Normal, grade 0-1, 12 donors; early OA, grade 1.5-2, 4 donors; severe OA, grade 3-4, 14 donors); no cluster's abundance changed with grade (permutation tests, all adjusted $p \geq 0.48$). **(c)** Neighborhoods of transcriptionally similar nuclei ($k = 30$ nearest neighbors in the 50 principal components used for clustering; 5% of nuclei sampled as neighborhood centers) shown at their index nucleus on the UMAP of Figure 3a, all 30 donors; color, log fold change in abundance per grade unit where spatial false discovery rate < 0.1 (negative binomial model of nuclei per donor, ~ technology + capture burden + grade, with the capture burden of the per-cluster grade model computed per donor); size, nuclei per neighborhood. **(d)** Log fold change per neighborhood by majority cluster; colored neighborhoods change with grade. **(e)** Log fold change against each neighborhood's mean score for the bulk grade signature of Figure 1 (expression of the genes up with grade minus those down, from the independent cohort of 92 knees). **(f)** Neighborhoods changing with grade per cluster in all 30 donors and in the 24 donors profiled with the multiome chemistry only (graph and neighborhoods rebuilt on those nuclei; ~ capture burden + grade); fewer with grade to the left, more to the right. **(g)** Spearman correlation between log fold change and the bulk grade-signature score within each cluster, in both analyses.

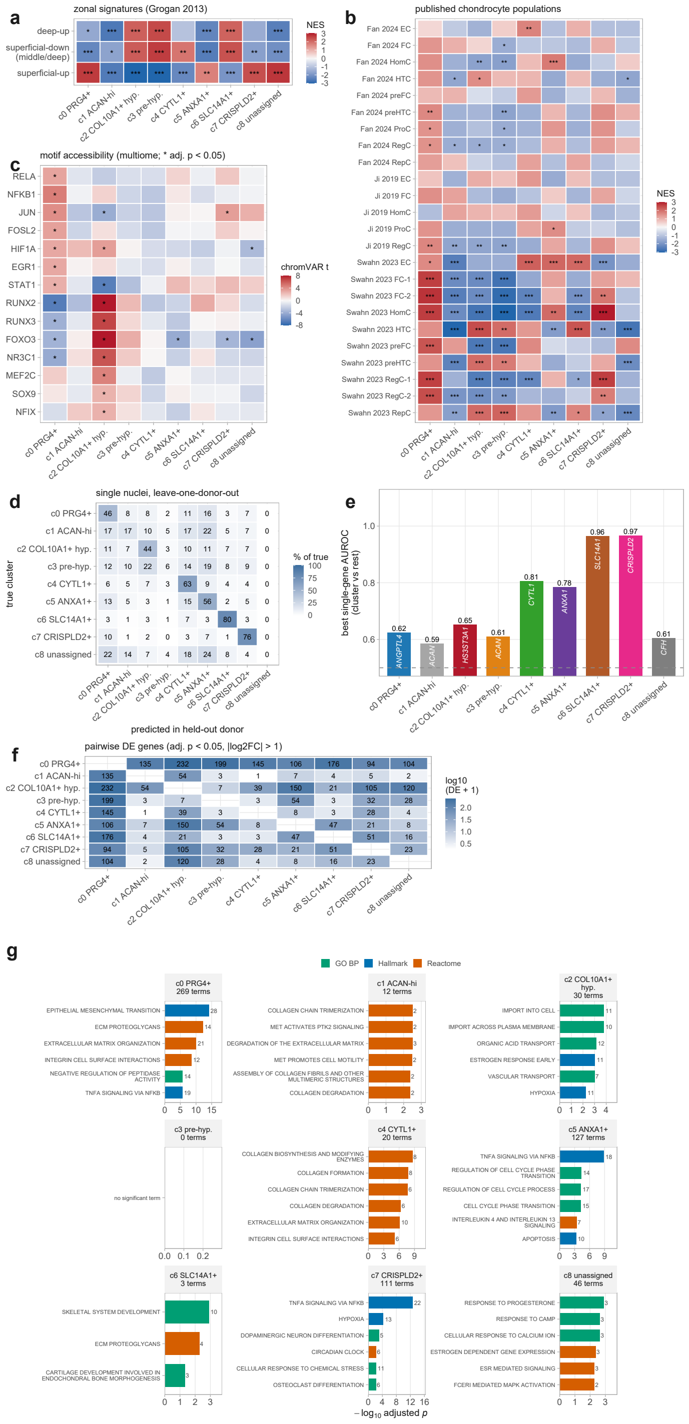


**Figure S9. Cluster identity: zonal, published-population, motif, separability and marker-pathway analyses.** Healthy donors (grade 0-1) as in Figure 4. **(a)** Enrichment of zonal gene sets (ref. 27) among genes ranked by each cluster's difference from the other clusters (normalized enrichment score, NES). **(b)** Enrichment of chondrocyte population signatures from published single-cell studies of OA cartilage (Ji 2019, Fan 2024, Swahn 2023; refs. 6, 9 and 8), as NES. **(c)** Motif accessibility (chromVAR) of each cluster against the others in the single-nucleus multiome, donor-blocked t statistic; stars, adjusted $p < 0.05$. **(d)** Single-nucleus classifier (multinomial elastic net trained on protein-coding genes excluding long genes, leave-one-donor-out): percentage of each cluster's nuclei assigned to each cluster in the test donor, which was not used for training; balanced accuracy 0.43 against 0.11 expected by chance. **(e)** Most specific single gene of each cluster (area under the receiver operating characteristic curve, AUROC, cluster against all other nuclei). **(f)** Number of genes differentially expressed between each pair of clusters (genes excluding long genes, adjusted $p < 0.05$ and absolute log2 fold change > 1). In (a-b) stars mark adjusted * $p < 0.05$, ** $p < 0.01$, *** $p < 0.001$. **(g)** Pathway enrichment of the cluster markers (over-representation analysis, long genes and ribosomal protein genes removed; Supplementary Methods): the six most significant gene sets per cluster from Hallmark, GO biological process and Reactome. Bars are $-\log_{10}$ adjusted p value and the number at each bar end is the count of marker genes in that set; note the free x scale per panel.

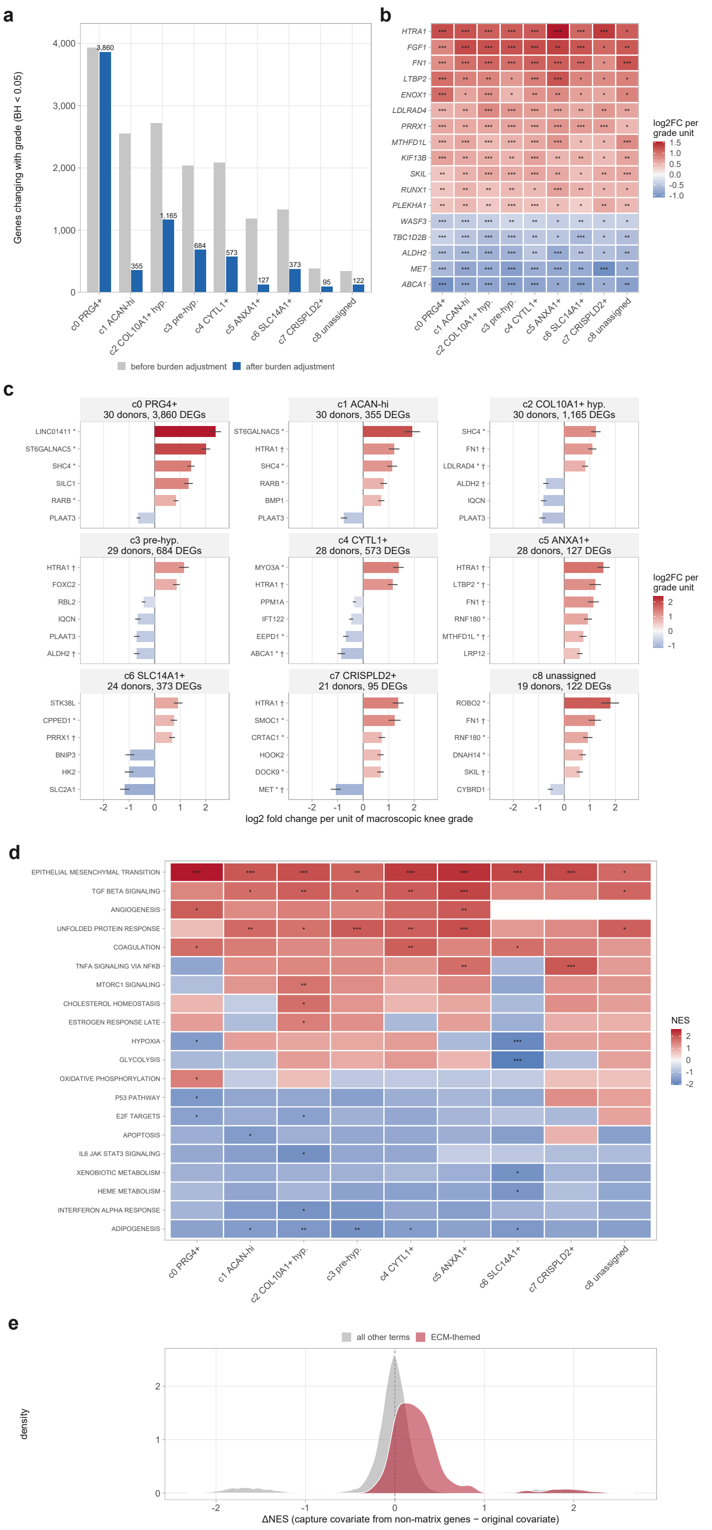


**Figure S10. Advancing OA is associated with a response shared by all nine chondrocyte clusters. (a)** Genes changing with grade in each cluster, before and after the capture-burden covariate. **(b)** The 17 genes changing significantly with grade in all nine clusters, all in the same direction in every cluster, as log2 fold change per grade unit; asterisks give that cluster's own adjusted p value, * below 0.05, ** below 0.01 and *** below 0.001. **(c)** The six genes with the smallest adjusted p value in each cluster, sorted within cluster by effect size; bars right of zero increase with grade and whiskers are one standard error. * marks genes longer than 8 times the 13.8 kb background span and † marks genes that also appear in panel b. **(d)** Grade-associated Hallmark pathways: all 20 Hallmark terms significant in at least one cluster, as normalized enrichment score; a blank tile means the term was not returned for that cluster. **(e)** Change in normalized enrichment score when the capture covariate is rebuilt from non-matrix genes only, split by whether the term is matrix-themed.

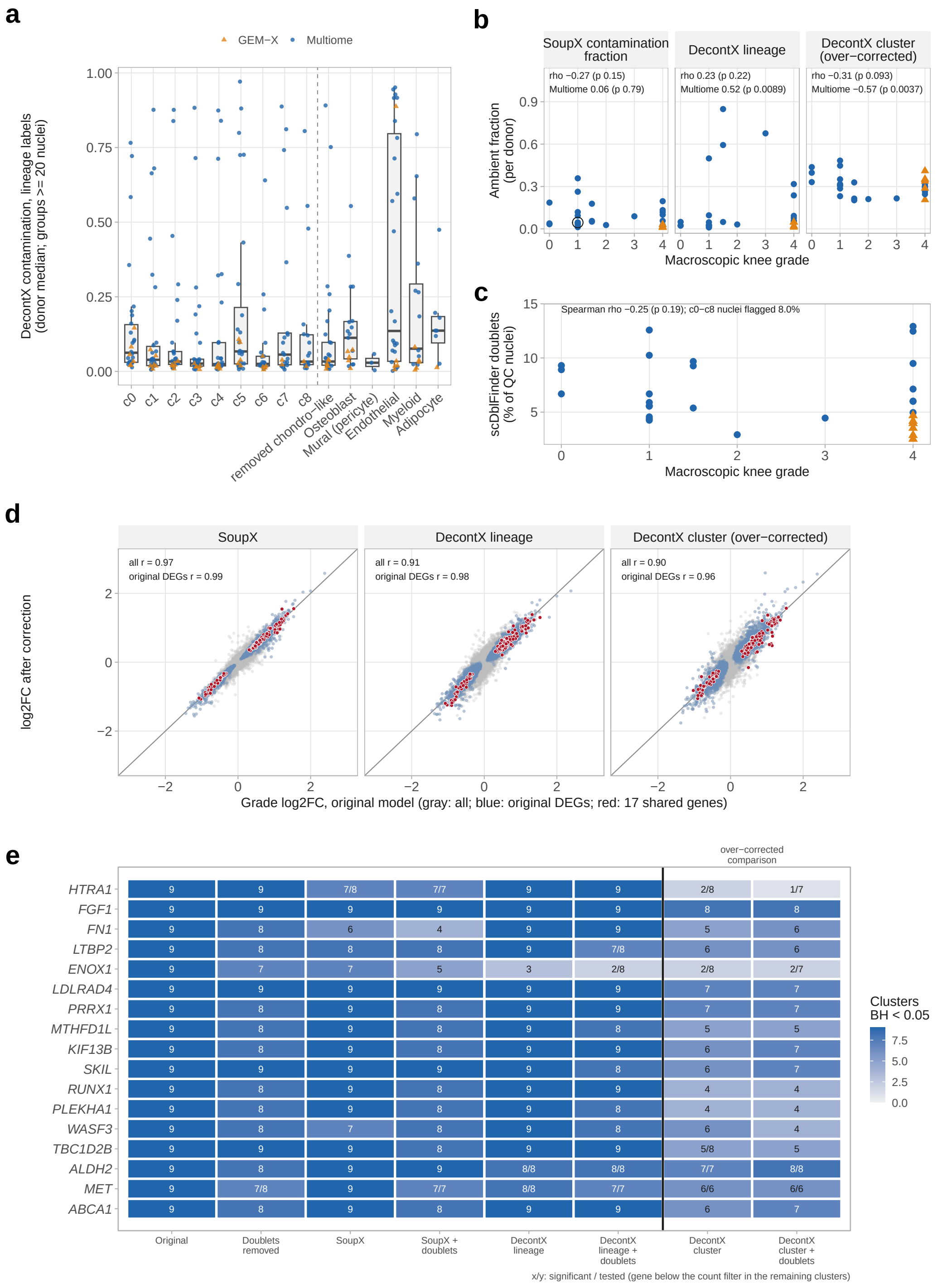


| | Original | Doublets removed | SoupX | SoupX + doublets | DecontX lineage | DecontX lineage + doublets | DecontX cluster | DecontX cluster + doublets |
|---|---|---|---|---|---|---|---|---|
| *HTRA1* | 9 | 9 | 7/8 | 7/7 | 9 | 9 | 2/8 | 1/7 |
| *FGF1* | 9 | 9 | 9 | 9 | 9 | 9 | 8 | 8 |
| *FN1* | 9 | 8 | 6 | 4 | 9 | 9 | 5 | 6 |
| *LTBP2* | 9 | 8 | 8 | 8 | 9 | 7/8 | 6 | 6 |
| *ENOX1* | 9 | 7 | 7 | 5 | 3 | 2/8 | 2/8 | 2/7 |
| *LDLRAD4* | 9 | 9 | 9 | 9 | 9 | 9 | 7 | 7 |
| *PRRX1* | 9 | 8 | 9 | 8 | 9 | 9 | 7 | 7 |
| *MTHFD1L* | 9 | 8 | 9 | 8 | 9 | 8 | 5 | 5 |
| *KIF13B* | 9 | 8 | 9 | 8 | 9 | 9 | 6 | 7 |
| *SKIL* | 9 | 9 | 9 | 9 | 9 | 8 | 6 | 7 |
| *RUNX1* | 9 | 8 | 9 | 8 | 9 | 9 | 4 | 4 |
| *PLEKHA1* | 9 | 8 | 9 | 8 | 9 | 8 | 4 | 4 |
| *WASF3* | 9 | 8 | 7 | 8 | 9 | 8 | 6 | 4 |
| *TBC1D2B* | 9 | 9 | 9 | 8 | 9 | 9 | 5/8 | 5 |
| *ALDH2* | 9 | 8 | 9 | 9 | 8/8 | 8/8 | 7/7 | 8/8 |
| *MET* | 9 | 7/8 | 9 | 7/7 | 8/8 | 7/7 | 6/6 | 6/6 |
| *ABCA1* | 9 | 8 | 9 | 8 | 9 | 9 | 6 | 7 |

**Figure S11. The shared grade response persists after ambient RNA correction and doublet removal. (a)** DecontX contamination estimate per donor (median over the nuclei of each group with at least 20 nuclei), with labels distinguishing chondrocytes from each non-chondrocyte lineage, for the nine chondrocyte clusters, chondrocyte-like nuclei removed during cleaning and the non-chondrocyte lineages; circles, multiome; triangles, GEM-X. **(b)** Ambient RNA fraction per donor against grade: SoupX contamination fraction (open circle, samples given the fallback estimate) and donor median DecontX contamination of c0-c8 nuclei with lineage labels and with cluster labels (over-corrected comparison); Spearman correlations over all donors and over multiome donors. **(c)** Percentage of nuclei called as doublets by scDblFinder in each donor against grade. **(d)** Log2 fold change per grade unit in the per-cluster pseudobulk model before and after correction with SoupX, DecontX with lineage labels and DecontX with cluster labels, each point a gene in a cluster; gray, all genes; blue, genes significant in the original model; red, the 17 shared genes; line, y = x. **(e)** Number of clusters in which each of the 17 shared genes remained significant (adjusted $p < 0.05$) under each correction, with and without removal of predicted doublets; the DecontX cluster-label arm is shown at the right for comparison. x/y, significant/tested where the gene fell below the expression filter in some clusters.

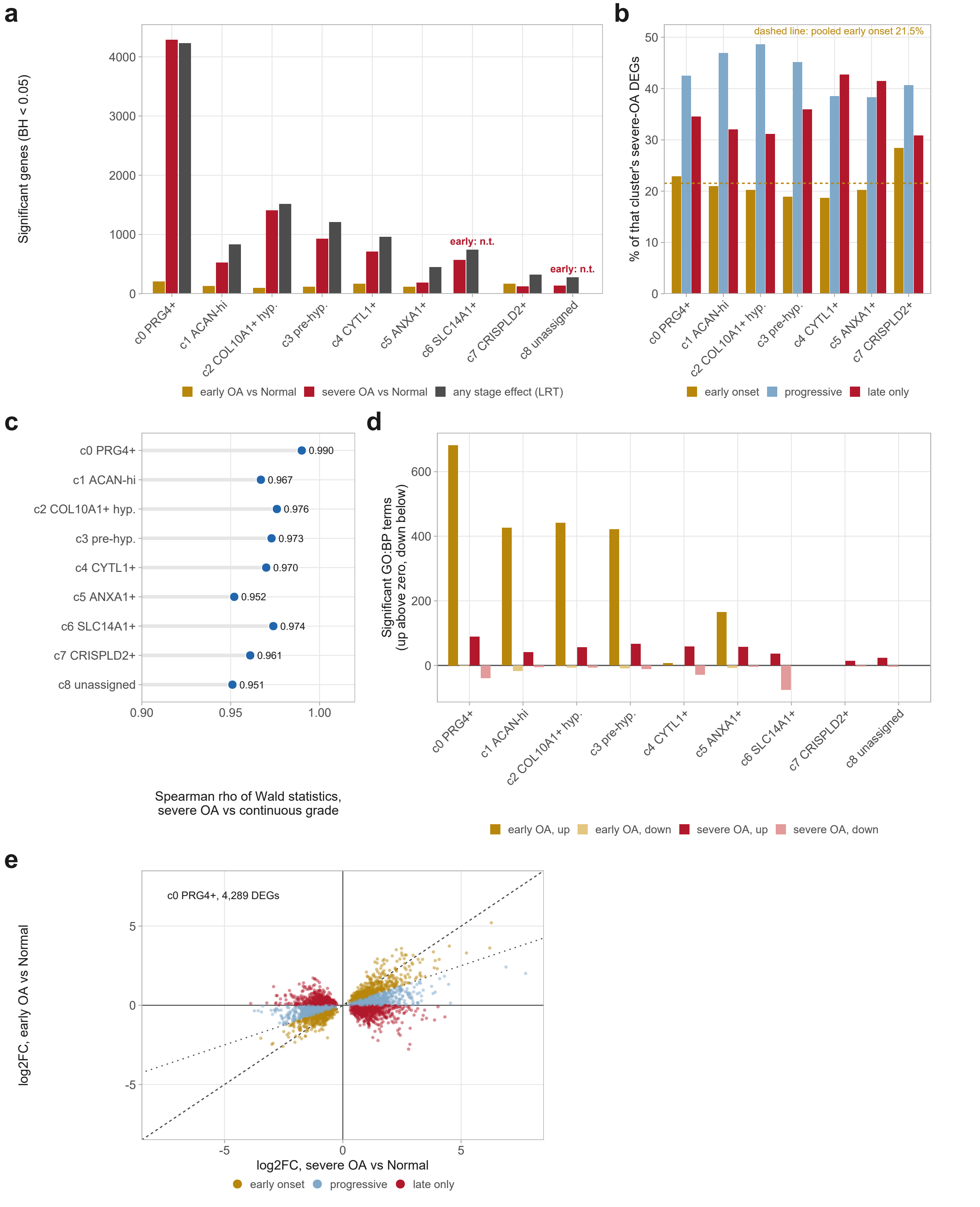


**Figure S12. Staging donors reproduces the continuous grade result. (a)** Genes changing in each stage contrast and in the test of any stage effect; n.t. marks a cluster (c6, c8) with only two early-OA donors, in which the early-OA contrast was fitted but not reported. **(b)** Each cluster's severe-OA genes classified by their early-OA effect as early onset, progressive or late only, as a percentage of that cluster's total; c6 and c8 are not shown (see a). **(c)** Spearman correlation of the severe-OA-versus-Normal statistics with those of the continuous grade model, per cluster. **(d)** Significant GO biological process terms per stage contrast, plotted upward for terms rising with stage and downward for terms falling. **(e)** Early-OA against severe-OA log2 fold change in c0; the dashed line is equality and the dotted line the half-effect boundary.

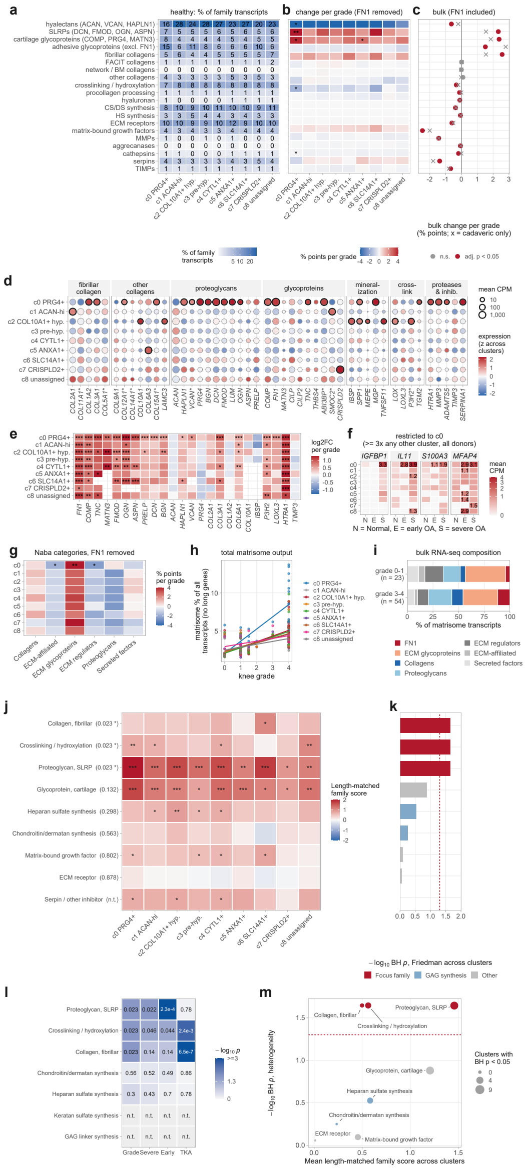

**Figure S13. Matrix composition per cluster: families, categories, cluster-restricted genes, bulk detail and between-cluster differences.** Single-nucleus donor-by-cluster pseudobulks with long genes excluded, as in Figure 6. **(a)** Share of each curated matrix family in each cluster's matrisome transcripts in healthy donors (grade 0-1; long Naba genes excluded, long family genes outside the Naba matrisome retained), with percentages in the tiles. **(b)** Change per grade unit in the share of each curated family with *FN1* removed (the 20 families shown; 21 of 24 contain non-long Naba matrisome genes), per cluster (stars, adjusted p). **(c)** Bulk cartilage RNA-seq (92 knees; transcripts per million): change per grade unit in the share of each curated family including *FN1*, so that the adhesive glycoprotein family includes *FN1* (share ~ batch + detected genes + grade); red, adjusted $p < 0.05$; crosses, the fit in cadaveric knees only, shown for information and not used as a filter because grade 4 knees were obtained almost entirely at arthroplasty. **(d)** Full matrix gene repertoire in healthy donors, as in Figure 6c. **(e)** Change per grade unit of selected matrix genes in each cluster (log2 fold change, per-cluster pseudobulk model; blank, not tested; stars, adjusted p). **(f)** Matrix genes restricted to *PRG4*+ superficial chondrocytes (c0) that change with grade (Figure 6d, double dagger): mean counts per million in each cluster in Normal (N, grade 0-1), early OA (E, grade 1.5-2) and severe OA (S, grade 3-4) donors; values of at least 1 are printed. Means in small clusters and in early OA (four donors) can reflect a single donor. **(g)** Change per grade unit in each matrisome category's share with *FN1* removed, per cluster (stars, adjusted p). **(h)** Total matrisome as a percentage of all transcripts against macroscopic knee grade, one point per donor and cluster, with a linear fit per cluster. **(i)** Bulk cartilage RNA-seq: composition of matrisome transcripts (transcripts per million) in grade 0-1 (23) and grade 3-4 (54) knees. **(j)** Length-matched response score for each extracellular matrix family in each cluster, for the nine families scored in all nine clusters; rows are ordered by the test in panel k and each row label carries that family's adjusted p value from it. Asterisks inside tiles are that cluster's own test; n.t., too few genes for the test in panel k. **(k)** Test of whether a family's response differs across the nine clusters; the dashed line marks adjusted $p = 0.05$. **(l)** The same test on four disease axes: continuous grade, severe OA versus Normal, early OA versus Normal, and Normal versus arthroplasty. Gray marks a family with too few genes tested in all nine clusters to test. **(m)** Mean family score against the heterogeneity test, with point size the number of clusters in which the family itself is significant. * $p < 0.05$, ** $p < 0.01$, *** $p < 0.001$.

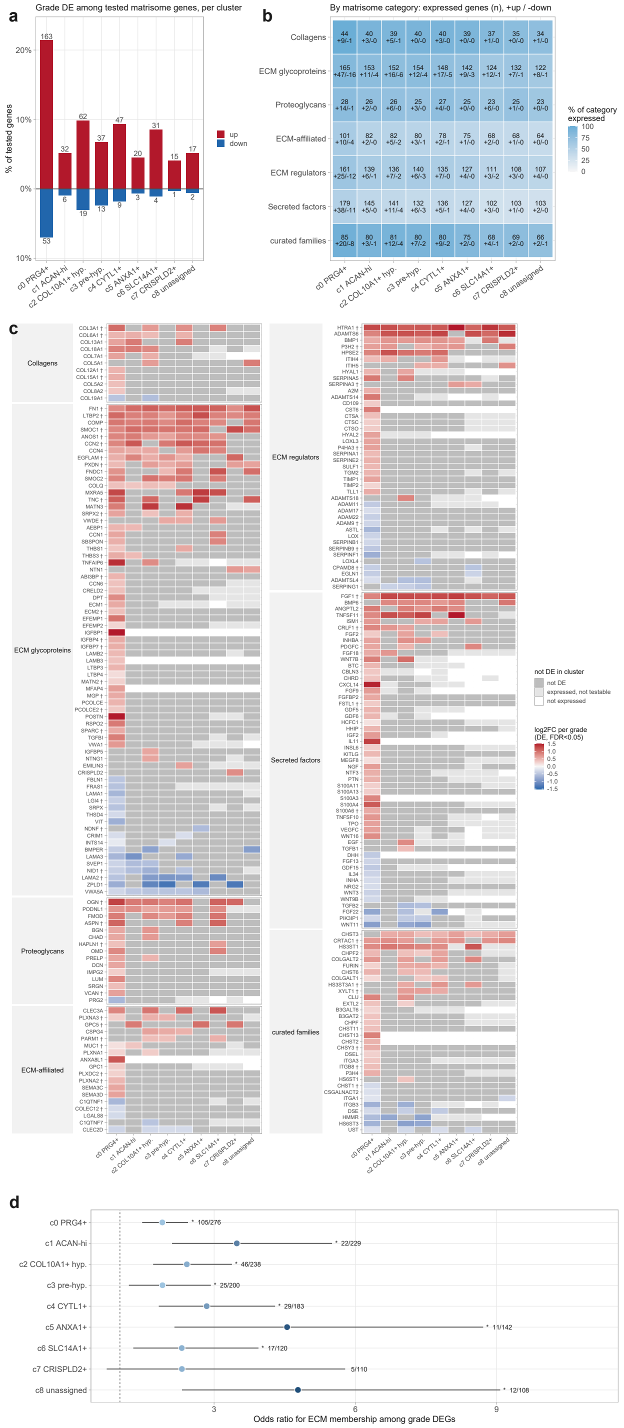


**Figure S14. Matrisome genes expressed in each chondrocyte cluster and their change with grade. (a)** Percentage of the tested matrisome genes that rose (red) or fell (blue) with grade in each cluster (per-cluster pseudobulk model, adjusted $p < 0.05$); numbers are genes. **(b)** Matrisome genes expressed in each cluster by category: number expressed, with the numbers that rose (+) and fell (-) with grade; fill, percentage of the category expressed. Expressed: mean counts per million of at least 1 over donors and at least three counts in at least three donors, or testable in the per-cluster grade model (at least 10 counts in half of the donors) or in a refit requiring at least 10 counts in at least three donors. **(c)** Every matrisome gene that changed with grade in at least one cluster (241 genes): log2 fold change per grade unit where adjusted $p < 0.05$; dark gray, expressed but not changed; light gray, expressed but too sparse to test; white, not expressed. 137 genes changed in a single cluster, 126 of them in c0, and none changed in opposite directions in different clusters. † marks genes whose grade response differs between clusters (cluster-by-grade interaction in a donor-random-intercept model over all clusters, Benjamini-Hochberg across the 346 matrisome genes in the interaction model; 111 of 346 genes, 55 of the 241 shown). **(d)** Enrichment of the 276 curated matrix-family genes among each cluster's grade-associated genes, as a Fisher odds ratio with 95% confidence interval; the number at each interval is that cluster's significant matrix genes over matrix genes tested. * adjusted $p < 0.05$ with a confidence interval excluding 1.

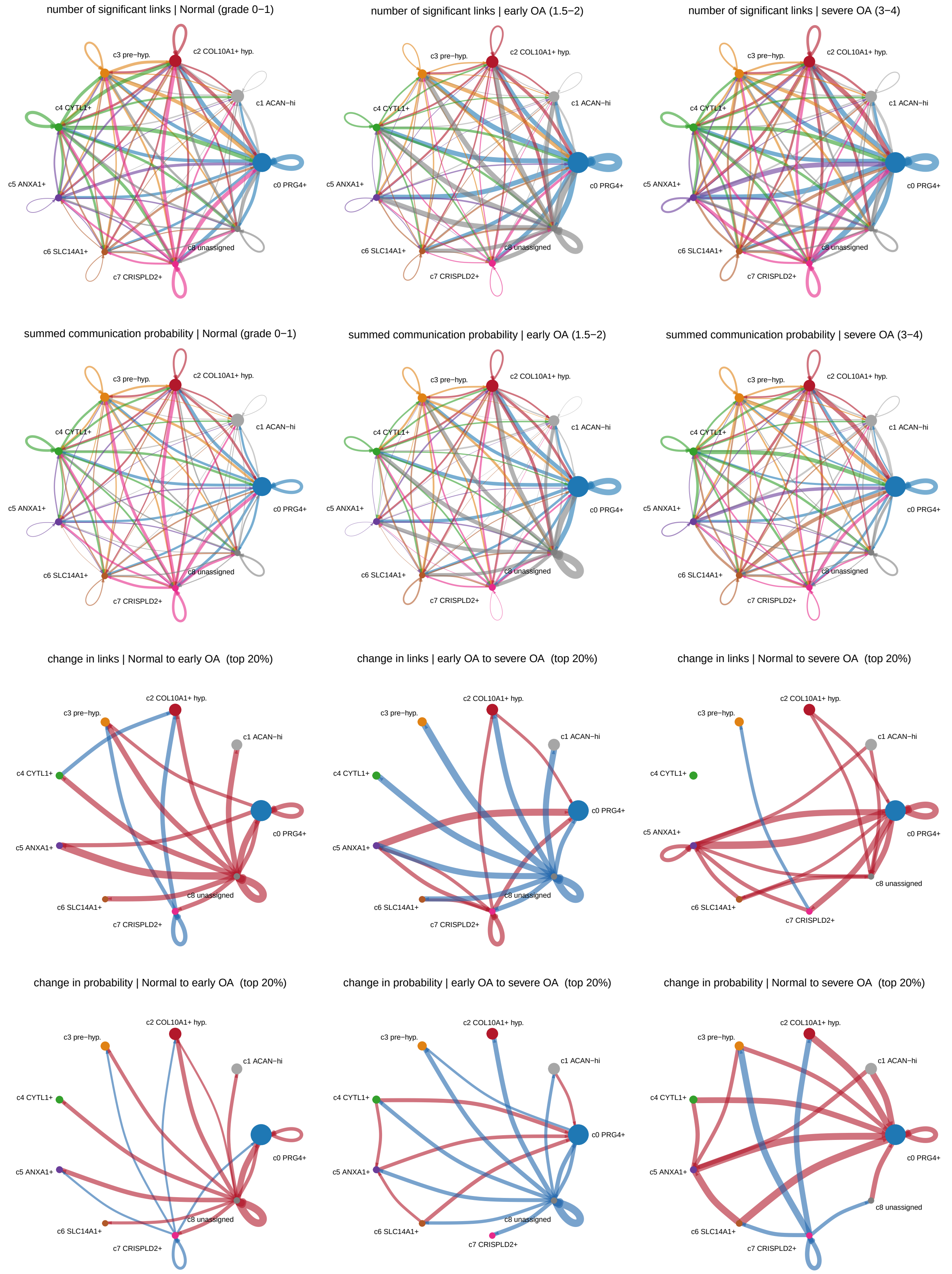


**Figure S15. CellChat network topology across stage.** Rows 1-2, aggregate networks over the nine clusters at each stage, as number of significant links and as summed communication probability; rows 3-4, the change in each measure across the three stage transitions, red increasing and blue decreasing in the later stage. Edge width shares one maximum within each row, so panels in a row are directly comparable. Node size is the percentage of that stage's nuclei in the cluster, and self-loops are autocrine links. Differential rows show only the strongest 20% of changes by absolute magnitude.

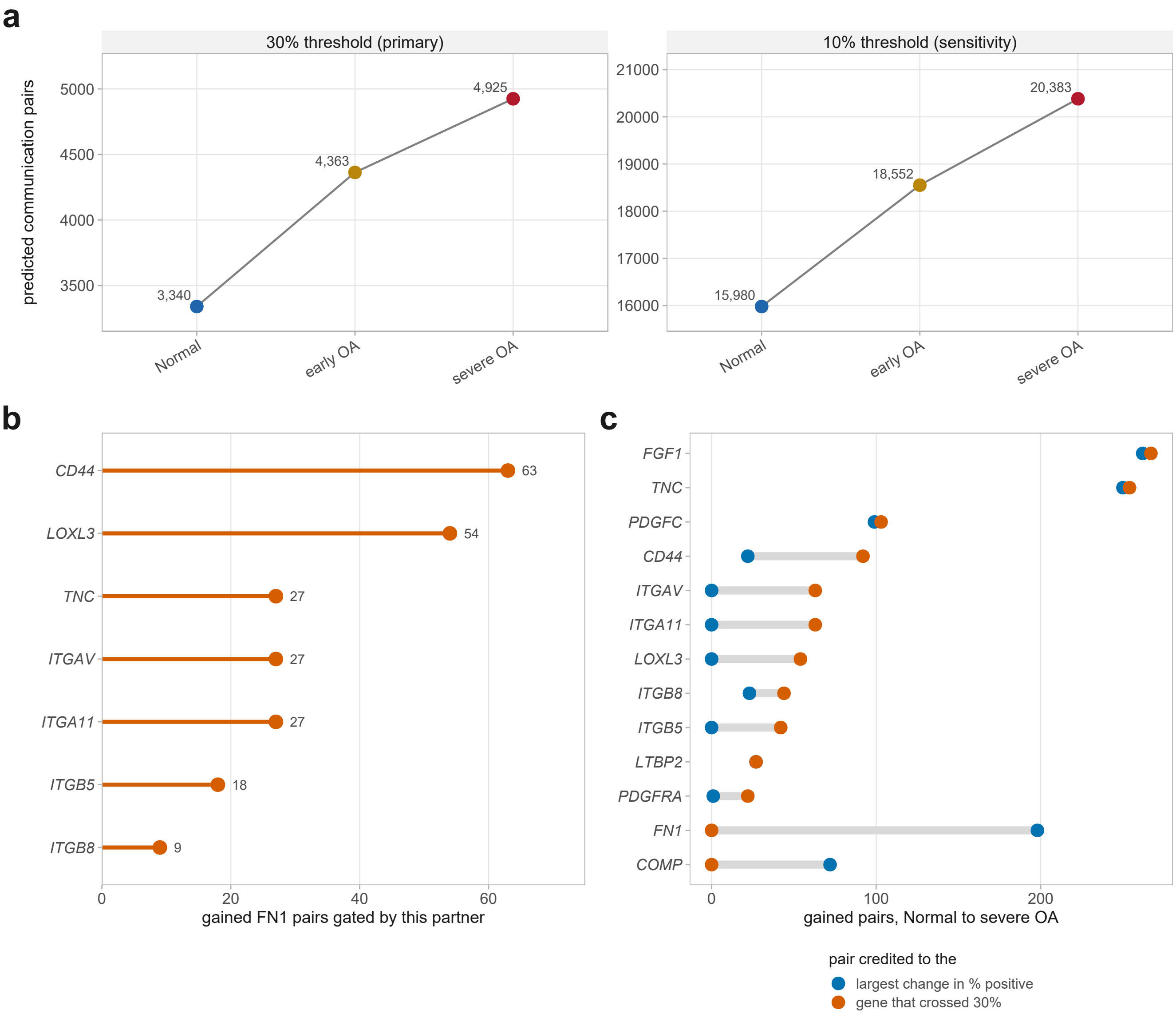


**Figure S16. MatriCom analysis supporting Figure 7. (a)** MatriCom predicted communication pairs at each stage, at the default 30% and a permissive 10% positive-population threshold. **(b)** The gating partner of each *FN1*-containing high-confidence pair gained between Normal and severe OA, meaning the partner gene that crossed the 30% threshold; no pair was gated by *FN1*. **(c)** Gained pairs credited to each gene under two rules, the gene whose percentage of positive nuclei changed most and the gene that crossed the 30% threshold, for genes with at least 20 pairs under either rule; gray bars join the two counts. Because a pair appears only when both of its genes clear the threshold, the second rule is the more appropriate attribution.

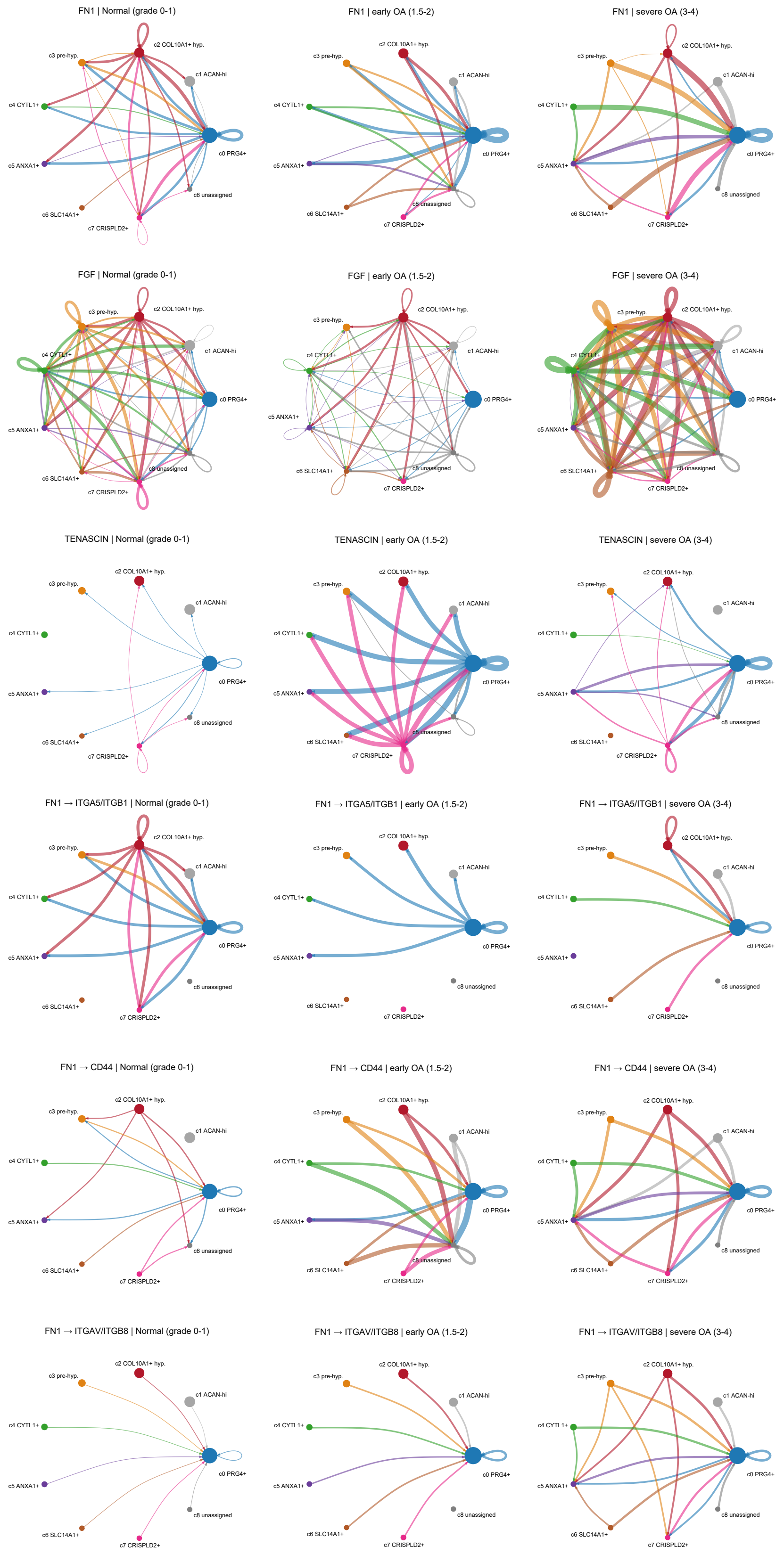


**Figure S17. Pathway and *FN1*-receptor networks by chondrocyte cluster.** Rows 1-3, the *FN1*, FGF and TENASCIN pathways at each stage, scaled to a shared maximum within each row, so stages are comparable within a pathway but rows are not comparable to one another. Rows 4-6, the three *FN1* receptors of Figure 7d whose share changed at each stage, on a single edge scale shared by all nine panels. An absent edge is a non-significant one.

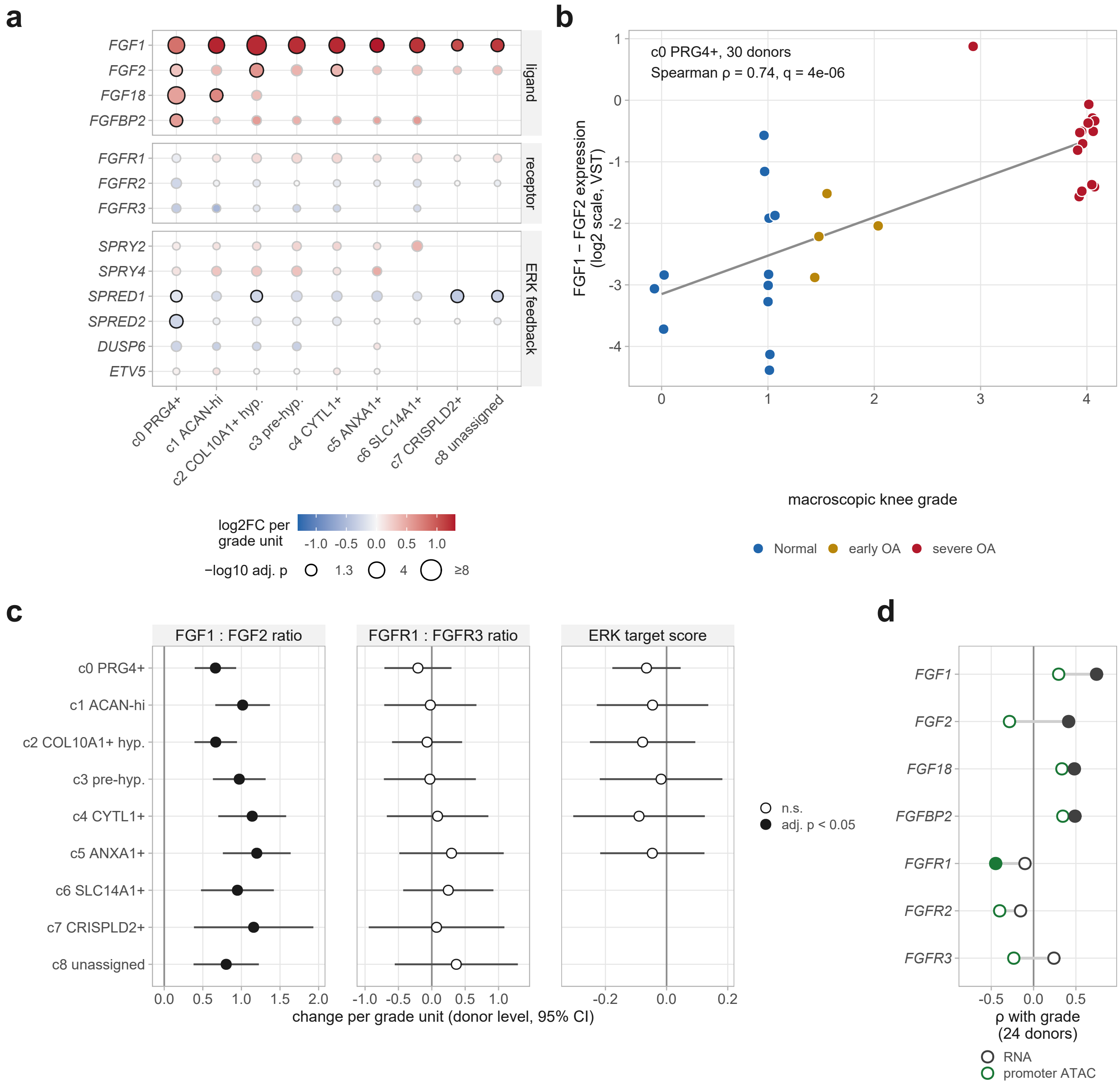


**Figure S18. *FGF1* rises relative to *FGF2* with grade without a rise in ERK output. (a)** Log2 fold change per grade unit (continuous-grade model of Figure S10) of FGF ligands, receptors and ERK feedback genes in each cluster; color is capped at ±1.3, size shows −log10 adjusted p and a black outline marks adjusted $p < 0.05$. A blank position means the gene was below the expression filter in that cluster. **(b)** *FGF1* relative to *FGF2* (difference of variance-stabilized expression) in each donor's c0 pseudobulk against macroscopic knee grade, 30 donors; the line is a linear fit for display. **(c)** Donor-level measures in each cluster regressed on grade with technology and capture burden as covariates, as change per grade unit with 95% confidence interval: the *FGF1*:*FGF2* and *FGFR1*:*FGFR3* expression ratios, and an ERK target score (mean z-scored variance-stabilized expression of *SPRY1*, *SPRY2*, *SPRY4*, *SPRED1*, *SPRED2*, *DUSP4*, *DUSP6*, *ETV1*, *ETV4*, *ETV5*, *IL17RD* and *FLRT3*, those with at least 10 counts in half of the cluster's donors, at least four genes required; not estimable in c6 to c8). Filled points are adjusted $p < 0.05$ (Benjamini-Hochberg across clusters within each measure). **(d)** Single-nucleus multiome, 24 donors: Spearman correlation with grade of each gene's RNA and of its promoter accessibility (transcription start sites ±1 kb, all *FGF1* promoters pooled), measured in the same nuclei; filled points are $p < 0.05$.

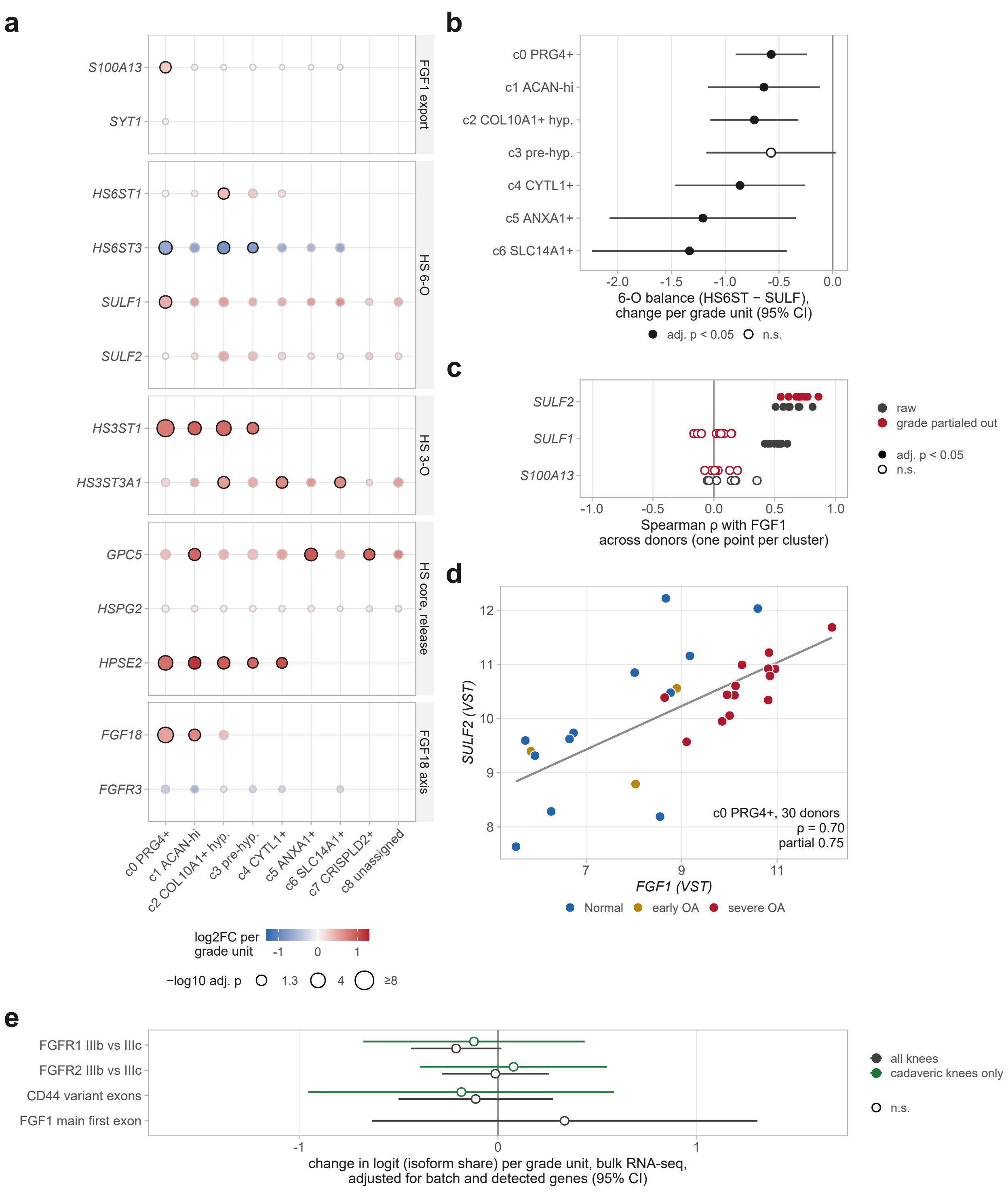


**Figure S19. Expression of *FGF1* export and heparan sulfate-modifying genes with grade. (a)** Grade association of genes for non-classical *FGF1* export (*S100A13*, *SYT1*), heparan sulfate (HS) 6-O and 3-O sulfation and 6-O removal (*SULF1*, *SULF2*), HS core proteins and heparanase-2, and the *FGF18*-*FGFR3* axis in each cluster, as in Figure 10a. **(b)** HS 6-O sulfation balance per donor (mean variance-stabilized expression of HS6ST1-3 minus that of *SULF1* and *SULF2*, in each donor's cluster pseudobulk), regressed on grade with technology and capture burden as covariates; bars are 95% confidence intervals and filled points adjusted $p < 0.05$ (Benjamini-Hochberg across clusters); c7 and c8 lack HS6ST expression. **(c)** Spearman correlation across donors between *FGF1* and *SULF2*, *SULF1* or *S100A13* within each cluster (one point per cluster), before (gray) and after (red) partialing out grade; filled points are adjusted $p < 0.05$. **(d)** *SULF2* against *FGF1* in each donor's c0 pseudobulk (30 donors), colored by stage. **(e)** Bulk RNA-seq (92 knees): change per grade unit in the share of IIIb among IIIb and IIIc junction reads for *FGFR1* and *FGFR2*, *CD44* variant-exon inclusion and the share of *FGF1* reads from its main first exon, adjusted as in Figure 2f, in all knees and in cadaveric knees only (not estimable for *FGF1*). *FGFR3* IIIb/IIIc (27 knees; $p = 0.50$) is not shown.

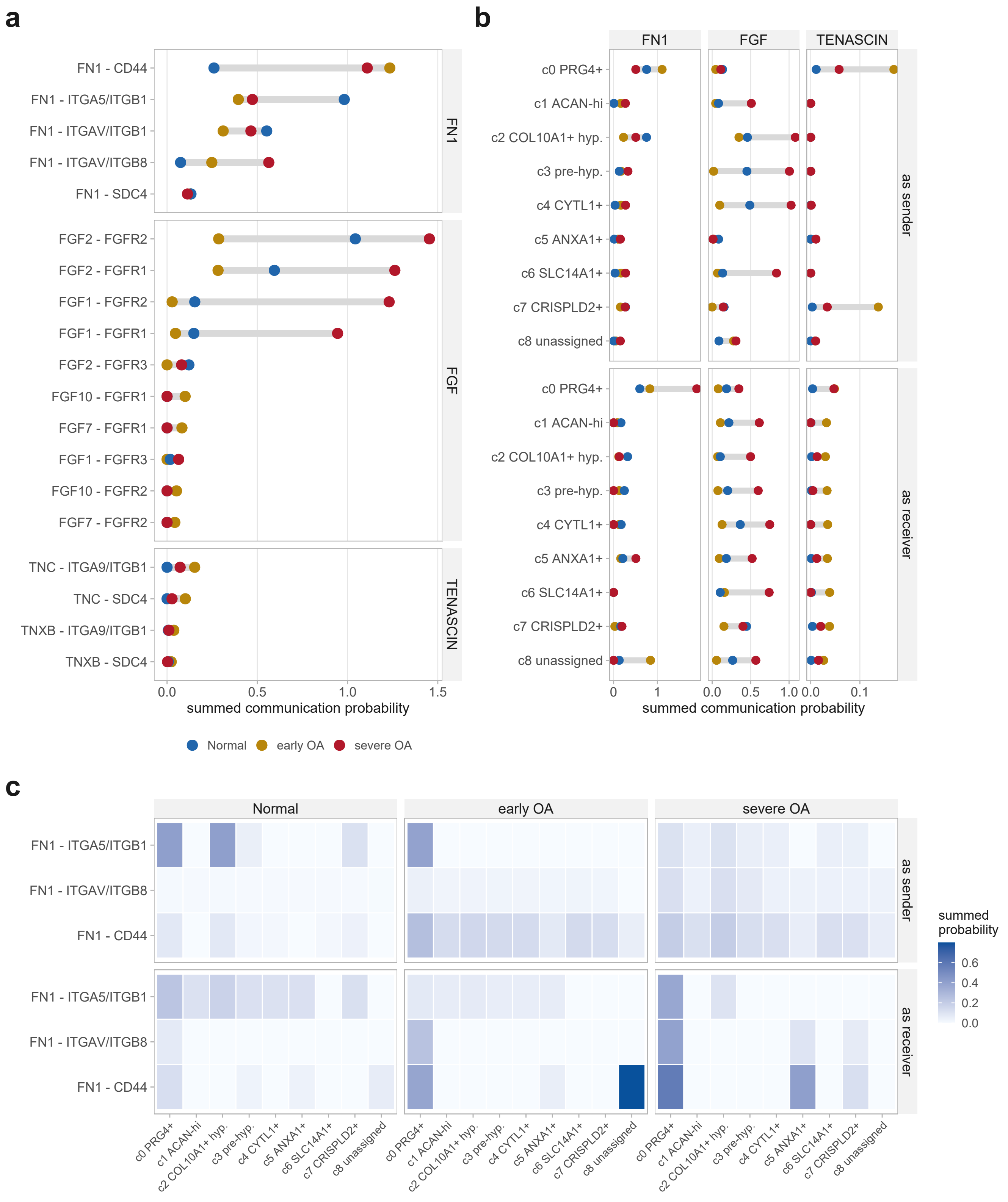


**Figure S20. CellChat ligand-receptor composition and sender/receiver detail. (a)** Summed probability of every ligand-receptor pair in the three pathways of Figure 7b at each stage; gray bars span the stages. **(b)** The same probability aggregated per cluster as the source (top) and target (bottom) of a link; autocrine links contribute to both, and the x axis is scaled per pathway. **(c)** Summed probability by cluster, role and stage, of the three *FN1* receptors of Figure 7d whose share changed.

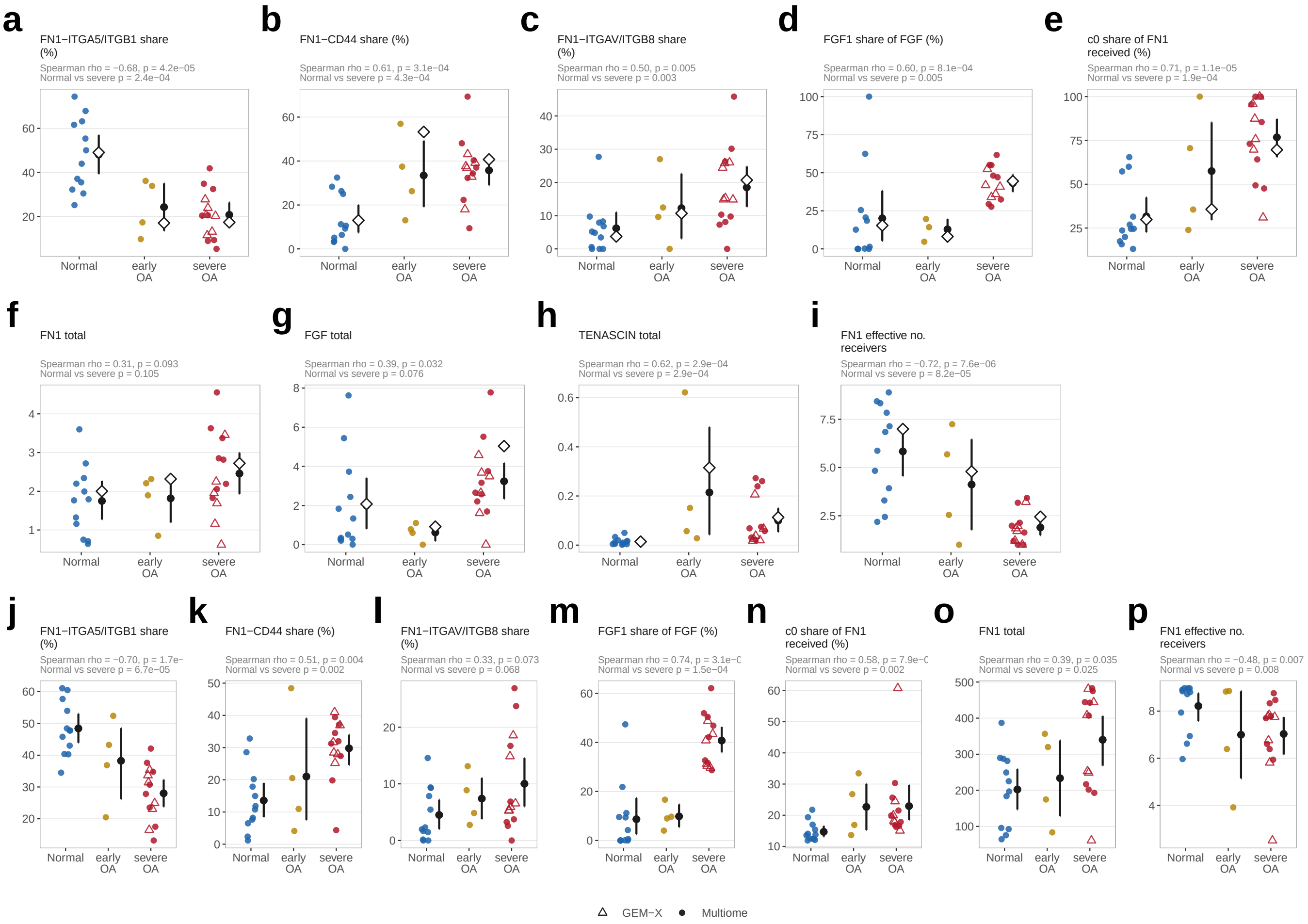


**Figure S21. Donor-level replication of the stage-level communication results.** CellChat was run separately in each of the 30 donors with the settings of Figure 7 (a-i), and a permutation-free expression-product score was computed for the same ligand-receptor pairs (j-p). Points are donors (circles, single-nucleus multiome; triangles, GEM-X); black point and bar, donor mean with 95% bootstrap confidence interval (5,000 resamples within stage); open diamond, pooled stage estimate of Figure 7. Panel subtitles give the Spearman correlation with grade (30 donors) and the Wilcoxon test of Normal (12) against severe OA (14) donors, unadjusted (adjusted values in the text). **(a-c)** Share of *FN1* probability through *ITGA5*/*ITGB1*, *CD44* and *ITGAV*/*ITGB8*. **(d)** *FGF1* share of FGF probability (28 donors with significant FGF links). **(e)** c0 share of *FN1* probability received. **(f-h)** Summed *FN1*, FGF and TENASCIN probability. **(i)** Effective number of FN1-receiving clusters. **(j-p)** Expression-product versions of the *FN1* receptor shares (j-l), *FGF1* share (m), c0 share of *FN1* received (n), total *FN1* (o) and effective number of FN1-receiving clusters (p).

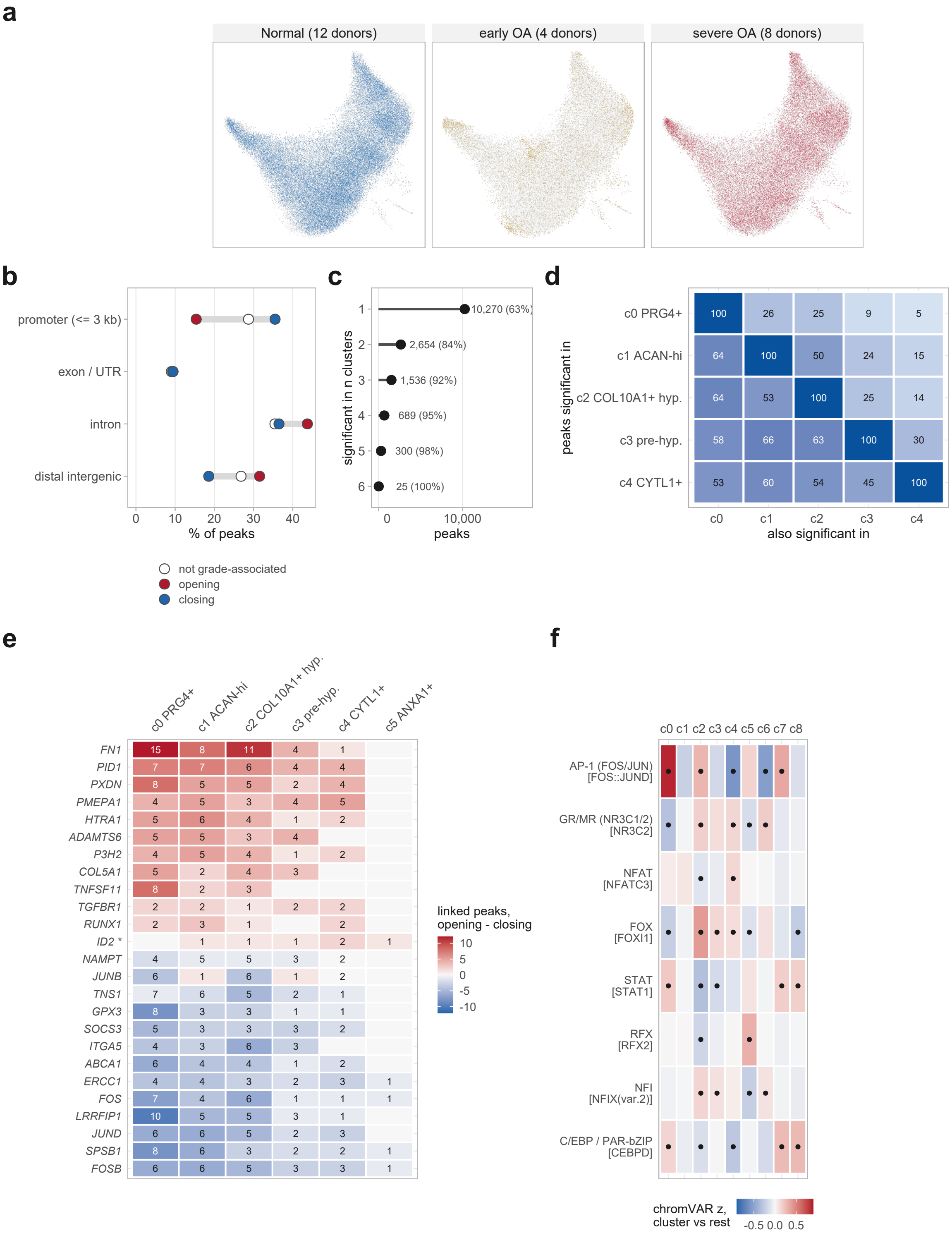


**Figure S22. Chromatin accessibility detail. (a)** The ATAC UMAP of Figure 9a split by stage, over all nuclei in gray. **(b)** Genomic context of opening and closing regions against all 158,325 tested regions not associated with grade. **(c)** Regions significant in at least one cluster (Figure 9c) by the number of clusters in which they are significant, with the percentage also significant in the pooled test in brackets. **(d)** Percentage of each row cluster's grade-associated regions also significant in the column cluster, for the five clusters with more than 1,000. **(e)** Linked grade-associated regions per gene and cluster (clusters c0-c5, those with significant linked regions); numbers are linked regions significant in that cluster, color is opening minus closing, and * marks genes not reached by the pooled analysis of Figure 9e. **(f)** Cluster identity in motif space, independent of grade: donor-mean chromVAR deviation of each cluster against the mean of the others (limma, donor blocked), one representative motif per transcription factor family in brackets; dots mark adjusted p value < 0.05.

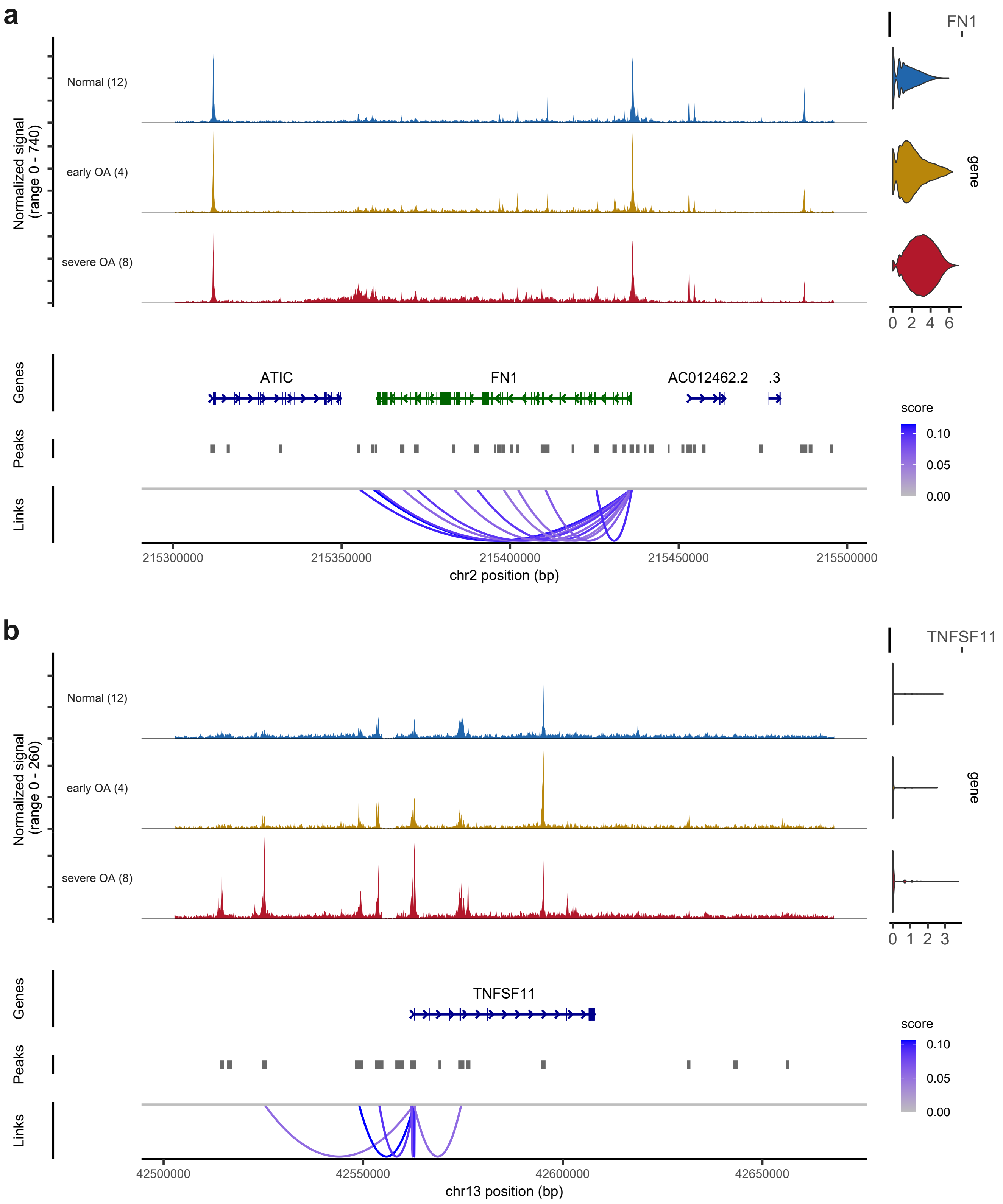


**Figure S23. Chromatin accessibility at *FN1* and *TNFSF11* by stage.** Normalized ATAC signal aggregated per stage across **(a)** *FN1* and **(b)** *TNFSF11* with 60 kb flanks, number of donors in brackets, with gene models, called peaks and peak-gene links colored by link score beneath; violins show the gene's RNA expression in the same nuclei. Coverage is shown per stage for display; the statistics are the continuous-grade tests of Figure 9.

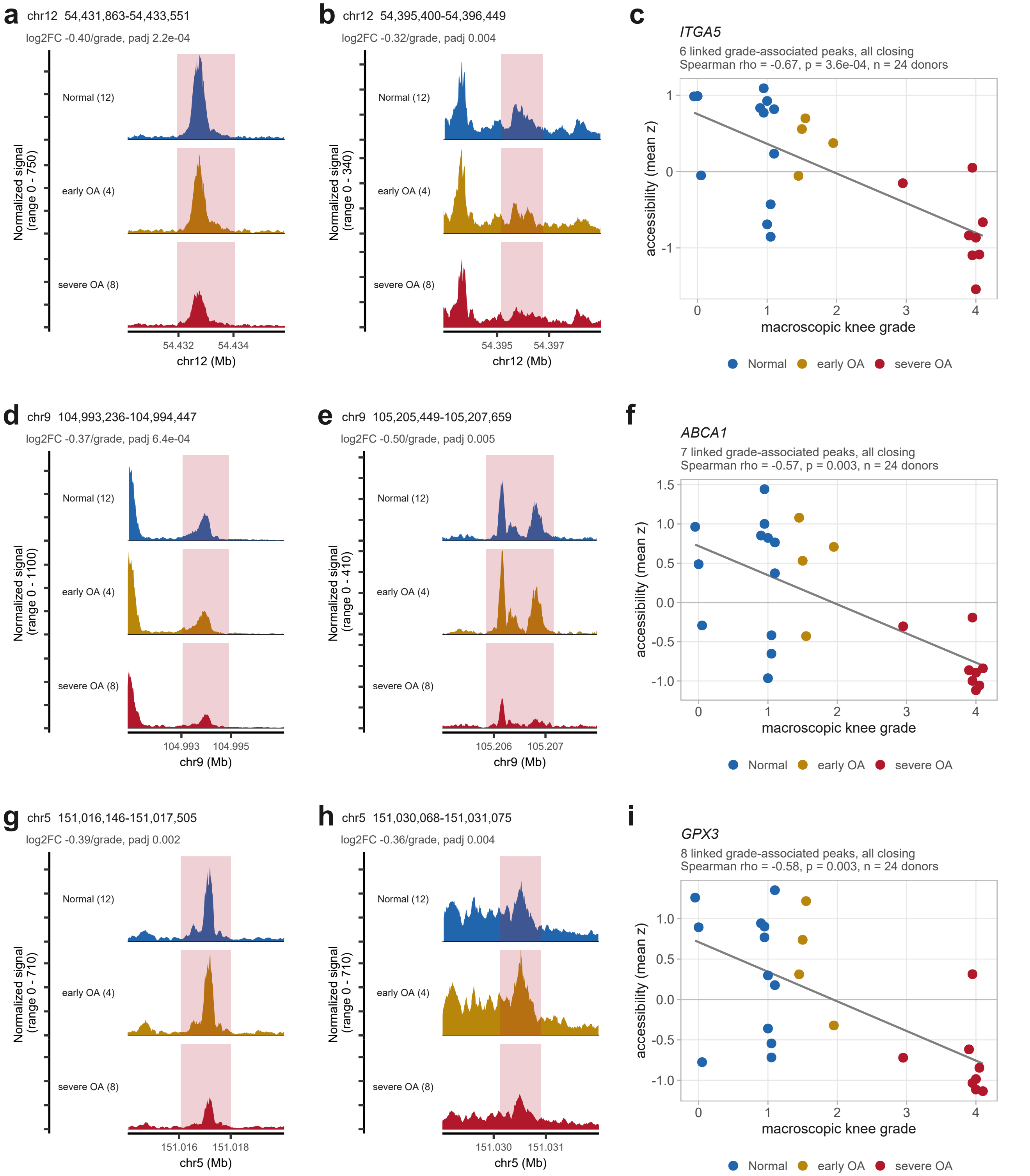


**Figure S24. Accessibility at the regions linked to *ITGA5*, *ABCA1* and *GPX3* falls with grade.** (a, b) *ITGA5*, (d, e) *ABCA1* and (g, h) *GPX3*: the two most significant of each gene's grade-associated linked regions, as normalized ATAC signal aggregated per stage (number of donors in brackets) over the region and 1.5 kb either side, with the region highlighted and its log2 fold change per grade unit and adjusted p value from Figure 9b above; each panel is scaled to its own window. (c, f, i) Donor-level accessibility of all of that gene's grade-associated linked regions (six for *ITGA5*, seven for *ABCA1* and eight for *GPX3*, all closing), as the mean across regions of per-region z-scored log2 counts per million in each donor's pseudobulk, against macroscopic knee grade; the line is a linear fit for display and the statistic is the Spearman correlation across the 24 donors. Coverage is shown per stage for display; the tests are the continuous-grade models of Figure 9.

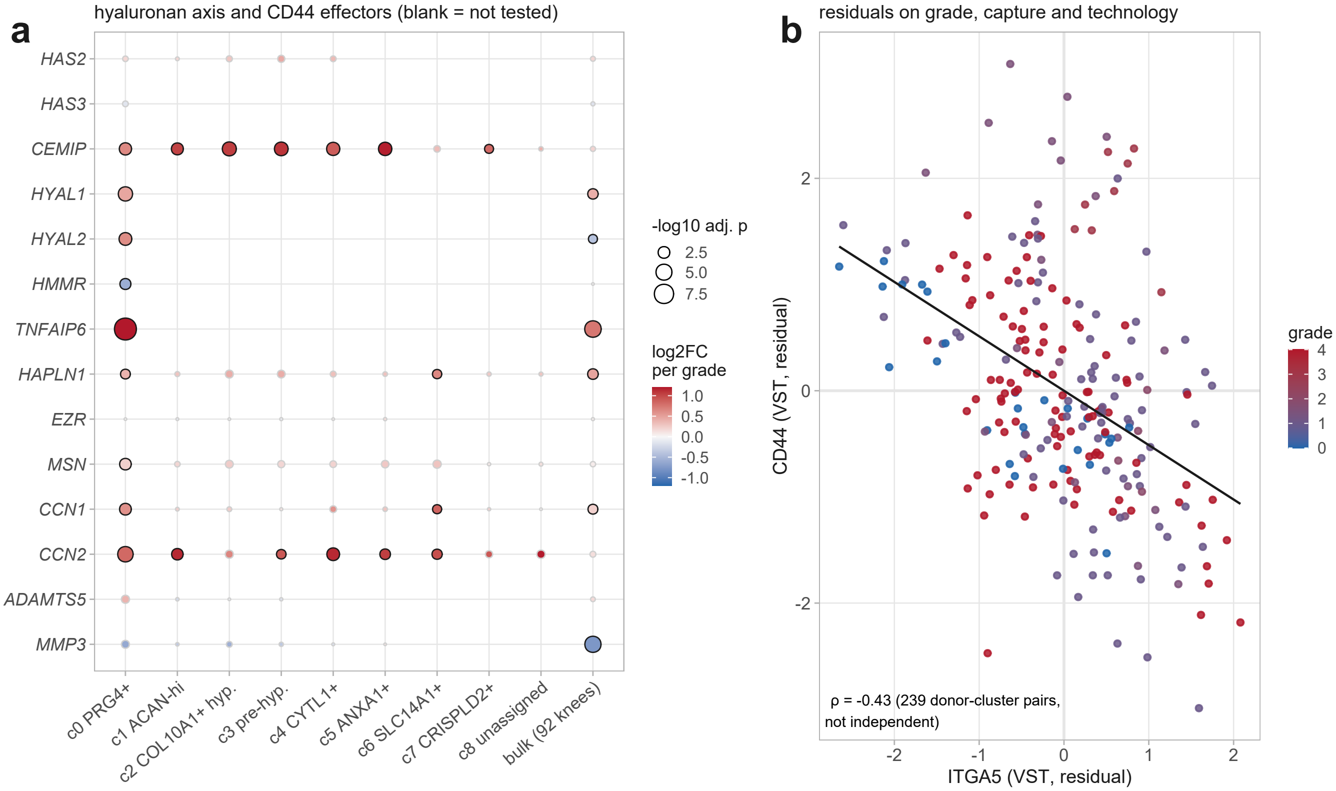


**Figure S25. Hyaluronan-axis genes and the donor-level *CD44-ITGA5* axis. (a)** Grade association of hyaluronan-axis genes and *CD44* effectors in each cluster and in bulk, displayed as in Figure 10a but with color capped at ±1.2; blank, not tested. **(b)** *CD44* against *ITGA5* (variance-stabilized expression, residuals on grade, capture burden and technology) for every donor and cluster; the pairs are not independent and the correlation is descriptive.

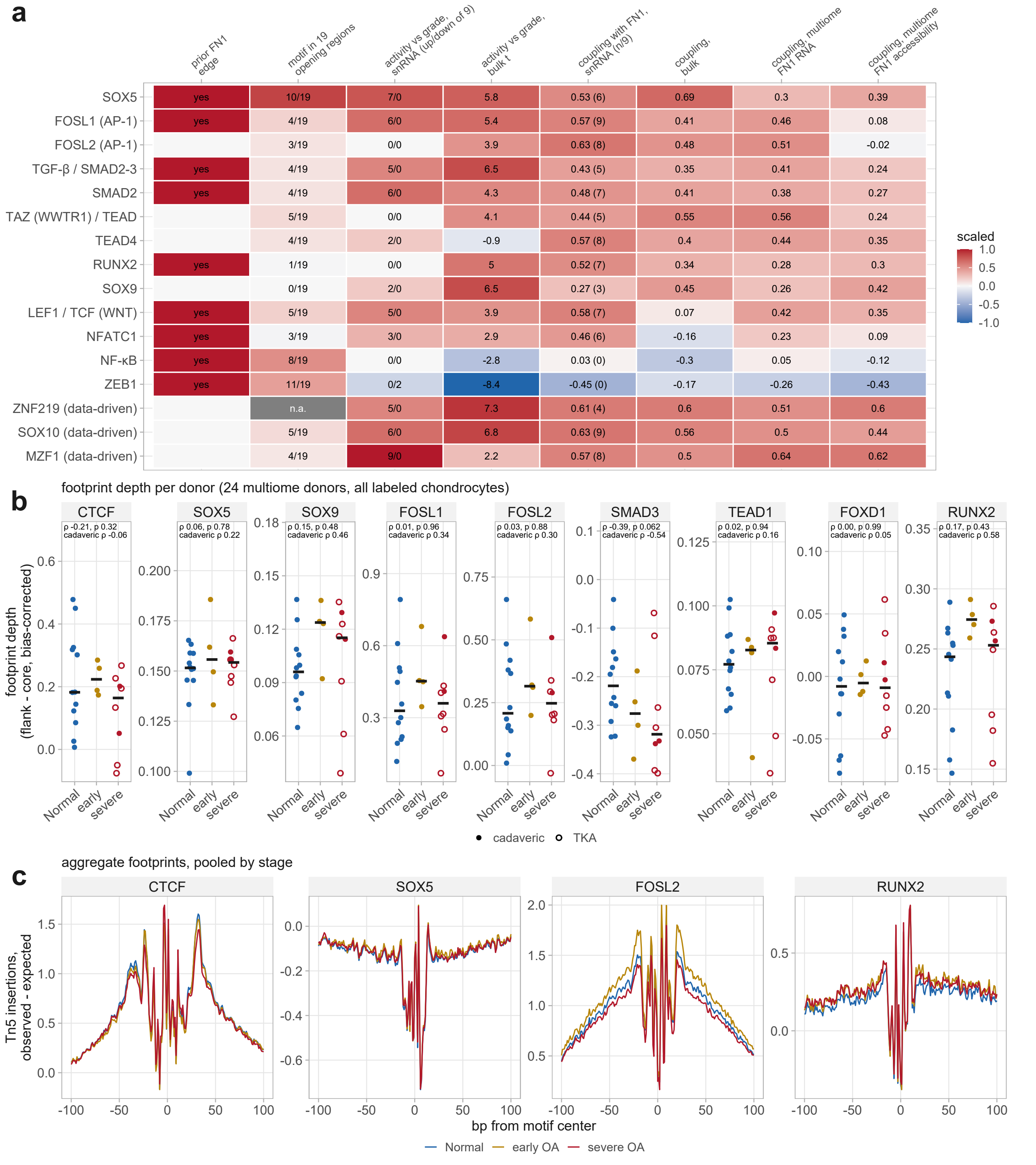


**Figure S26. Candidate transcriptional regulators of *FN1*: evidence and footprinting. (a)** Evidence for each candidate: prior *FN1* regulatory edge (CollecTRI/DoRothEA); occurrence of the motif in the 19 *FN1*-linked regions that opened with grade (JASPAR; the strongest among the factors shown, *SOX5*, nominal p = $4\times10^{-4}$, adjusted p = 0.14; n.a., gray, no JASPAR motif for the factor); activity (inferred from target genes) against grade in the snRNA-seq clusters (clusters up/down of nine at nominal $p < 0.05$) and the bulk cohort (t); and coupling of activity with *FN1* beyond grade (partial Spearman correlation; median over clusters with the number of clusters at nominal $p < 0.05$, bulk, and multiome *FN1* RNA and *FN1* linked-region accessibility). Data-driven rows are factors without prior *FN1* edges that coupled as strongly. **(b)** Footprint depth (flank minus core accessibility, corrected for Tn5 insertion bias) per donor by stage in all labeled chondrocytes of the 24 multiome donors; Spearman correlation with stage across all donors and in cadaveric donors only (filled, cadaveric; open, arthroplasty). **(c)** Aggregate bias-corrected Tn5 insertion profiles around *CTCF* (control), *SOX5*, *FOSL2* and *RUNX2* motifs, pooled by stage.

**Table S1. Identity, distinctness and grade response of the nine chondrocyte clusters.** For each cluster: nuclei, summary verdict, position on the zonal and hypertrophy axes, defining genes and programs in the 12 healthy donors, cross-donor recognition and stability on reclustering (Figure 4, Figure S9), cluster-specific chromatin, the matrix produced in healthy donors (Figure 6), cluster-distinctive grade-associated genes, pathways and matrix genes (Figure 5, Figure 6d), abundance against grade (centered log-ratio share, Spearman correlation with 10,000 donor-label permutations) and the share of the bulk grade genes most highly expressed in the subset (Figure S1g). Provided as an Excel file.
